# Evaluation of the genotoxic potential of dietary metabolites to support the renewal of an active substance under European Commission Regulation (EC) 1107/2009 – fludioxonil case study

**DOI:** 10.1101/2025.09.17.676779

**Authors:** Zofi A. McKenzie, Anne-Sophie Parant, Katy L. Bridgwood, Ewan D. Booth

## Abstract

The evaluation of the genotoxic potential of dietary metabolites of active substances is an important aspect for the (re)registration of pesticides within the European Union. This paper presents a case study evaluating the genotoxic potential of three dietary metabolites (CGA335892, SYN518580 and CGA227731) of the active substance fludioxonil under Regulation (EC)1107/2009. This tiered process started with (Q)SAR and read across, to enable a “grouping” approach, the three metabolites were identified as “exemplar compounds” and evaluated for gene mutation, clastogenicity and aneugenicity. Bacterial reverse mutation tests (*S. typhimurium*. and *E. coli* strains) were conducted on CGA335892 (negative), SYN518580 (negative) and CGA227731 (positive; TA1537 ±S9; 500-1500 µg/plate). *In vitro* micronucleus assays in TK6 cells on CGA335892 and SYN518580 were positive at ≤7 and ≤25 µg/mL respectively whereas CGA227731 was clearly negative in human lymphocytes. *In vivo* genotoxicity assays were conducted to follow up the positive *in vitro* findings. Mouse micronucleus studies (CGA335892, SYN518580) were negative up to the maximum tolerated / limit dose respectively; CGA335892 and SYN518580 were concluded to be not clastogenic and not aneugenic *in vivo*. CGA227731 was negative in a rat Comet assay (liver, duodenum); however, following regulatory review a transgenic rodent gene mutation assay was conducted (BigBlue® rats, up to limit dose) to investigate the *in vivo* mutagenicity of this metabolite. This study was negative (liver, duodenum, bone marrow); CGA227731 is concluded to be non-mutagenic *in vivo*. Based on this extensive battery of genotoxicity tests, CGA335892, SYN518580 and CGA227731 are concluded to be of no genotoxic concern.

**Highlights:**

- Genotoxicity evaluation of dietary metabolites under European data requirements
- Tiered dietary metabolite evaluation using (Q)SAR, read across and genotoxicity testing
- Use of apical endpoint genotoxicity studies to follow up an indicator assay
- Demonstration of lack of genotoxicity for fludioxonil dietary metabolites

## 1. Introduction

Fludioxonil is a broad-spectrum contact fungicide with residual activity, demonstrating high efficacy against diseases caused by Ascomycetes, Basidiomycetes, and Deuteromycetes. Used extensively worldwide, it is available both as a solo formulation and in ready-mix combinations with other active substances. Its applications include foliar sprays for preventive treatment on grapes, specialty crops, and vegetables, as well as post-harvest treatments for various fruit crops using diverse application methods. In seed treatment, fludioxonil serves as a cornerstone ingredient in multiple combinations, controlling a wide range of soil- and seed-borne diseases including snow mold (*Monographella nivalis*), *Fusarium spp*., common bunt (*Tilletia caries*), *Septoria spp*., leaf stripe (*Pyrenophora graminea*), and numerous other pathogens affecting major arable crops and vegetables.

In Europe, the active substance fludioxonil recently underwent a comprehensive review as part of the Active Ingredient Renewal (AIR3) process, which resulted in the publication of EFSA’s peer review (EFSA Conclusions, 2024). Under Regulation (EC) 1107/2009, this assessment framework requires rigorous evaluation of the active substance and its metabolites. This assessment requires the evaluation of the genotoxic potential of all identified metabolites in plants or animals, hereafter referred to as ‘dietary metabolites’. The process begins with *in silico* tools, including (Q)SAR predictions and structural alerts analysis, followed by read-across approaches and genotoxicity testing where applicable.

The (Q)SAR process should use recognised validated (Q)SAR models that follow the 5 OECD principles of a robust (Q)SAR model addressing relevant genotoxicity endpoints (OECD 2024). An assessment should use at least two models based on different approaches, for example the combination of statistical model(s) and knowledge-based model(s) in an orthogonal process. The combined output from the models may then be reviewed using human expert knowledge (Powley 2015).

Read across is a technique to predict toxicity of one chemical using data from similar chemicals and is based on the principle that similar chemicals have similar biological effects. An assessment of the chemical and/or biological similarity between substances is performed to determine whether data for the data-rich “source” substance can be used to predict the toxicity of the “target” substance. There are a variety of guidance documents that describe the process in more detail (ECHA 2017, OECD 2017b, EFSA 2025)

For metabolites requiring experimental evaluation, EFSA requires the genotoxicity endpoints of gene mutation, structural and numerical chromosome aberrations to be addressed (EFSA, 2011). The standard testing strategy involves a core battery of *in vitro* tests, typically the bacterial reverse mutation assay (OECD 471) and the *in vitro* micronucleus test (MNvit; OECD 487). Exceptionally, a mammalian gene mutation assay (OECD 490 or 476) may be required if there is a mammalian gene mutation specific (Q)SAR alert. When *in vitro* results are positive or inconclusive, appropriate *in vivo* follow-up testing is conducted, guided by the Scientific Opinion on genotoxicity testing strategies (EFSA, 2011) for assay selection and result interpretation. Studies conducted to support the (re)registration of an active substance should be conducted under the appropriate Good Laboratory Practice (GLP) and in accordance with the relevant OECD test guideline. Literature data may be used in the assessment of the genotoxic potential of dietary metabolites only when it is evaluated to be of sufficient quality and robustness (Klimisch score of 1 or 2 required (Klimisch *et al.,* 1997)).

This tiered strategy, encompassing (Q)SAR, read-across, and data generation, aims to minimize unnecessary testing while maintaining robust safety assessment standards. Any read-across applications must be scientifically justified based on structural similarities, common functional groups, and the preservation or loss of structural alerts relative to the parent or exemplar compound. Regulatory acceptance of read-across is higher within the European framework if the exemplar compound for each group has experimental data covering all the required genotoxicity endpoints (i.e. (Q)SAR alone may not be sufficient to exclude the genotoxic potential of a compound or group of compounds).

Fludioxonil itself, and the majority of its dietary metabolites have been evaluated for their genotoxic potential within the Regulation (EC)1107/2009 framework and have been concluded to be non-genotoxic (ANSES 2023). Genotoxicity data on two metabolites (CGA335892 and SYN518580) was not available at the time of the regulatory deadline. A third metabolite, CGA227731, was concluded to be genotoxic by the EU regulators, based on their review during the renewal process of *in vitro* studies and a follow up *in vivo* Comet assay. Subsequently, a transgenic rodent gene mutation assay (TGR) has been conducted to investigate the *in vivo* mutagenicity of this compound. This publication presents a case study of the genetic toxicity evaluation for these three dietary metabolites of fludioxonil and demonstrates the application of this tiered strategy. The chemical structures and descriptions of chemical changes between fludioxonil and these metabolites are presented in Table 1.

**Table 1:**
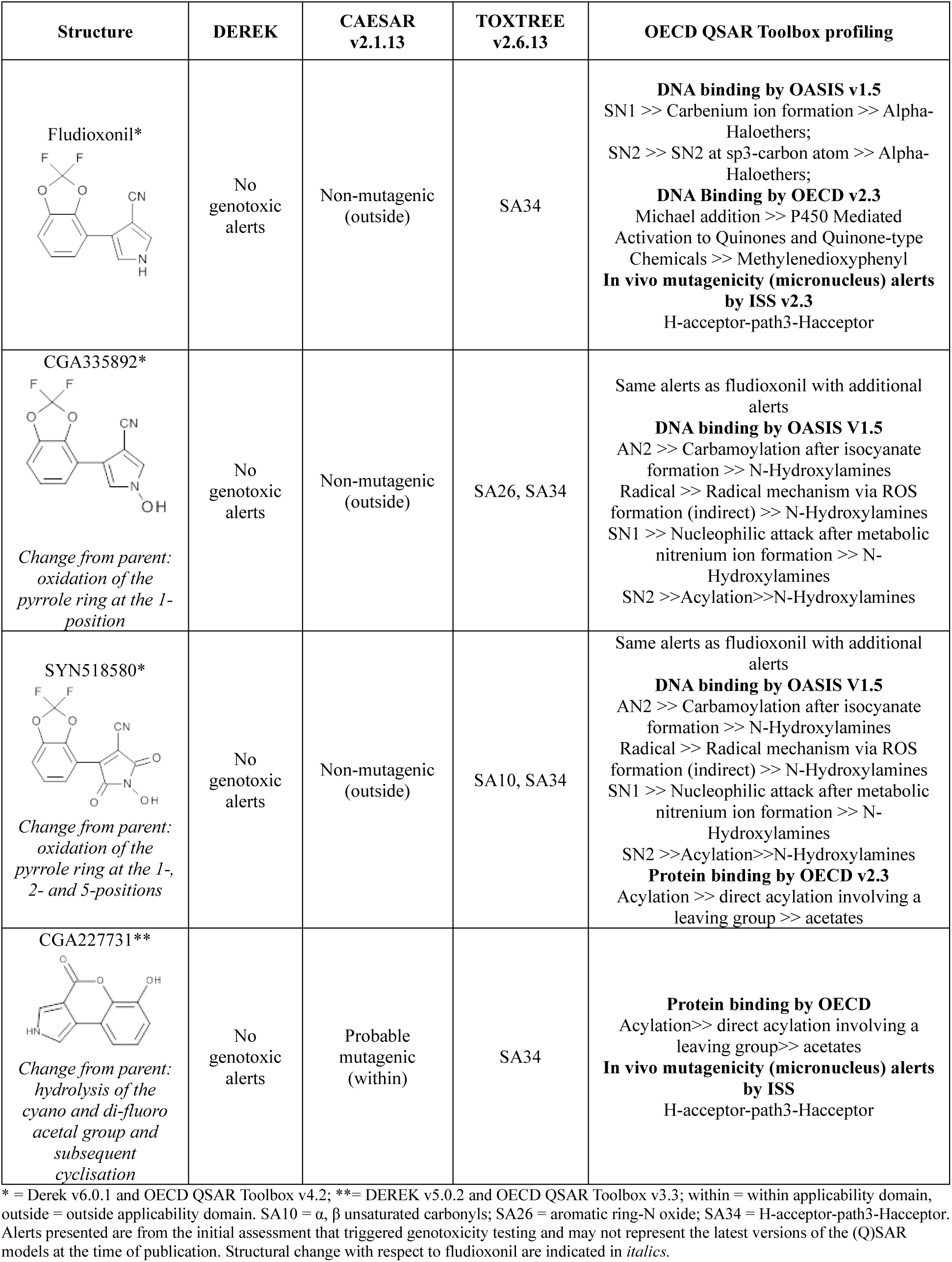
Summary of (Q)SAR Predictions and Alerts.

| Structure | DEREK | CAESAR v2.1.13 | TOXTREE v2.6.13 | OECD QSAR Toolbox profiling |
| --- | --- | --- | --- | --- |
| Fludioxonil* | No genotoxic alerts | Non-mutagenic (outside) | SA34 | **DNA binding by OASIS v1.5**<br>SN1 >> Carbenium ion formation >> Alpha-Haloethers;<br>SN2 >> SN2 at sp<sup>3</sup>-carbon atom >> Alpha-Haloethers;<br>**DNA Binding by OECD v2.3**<br>Michael addition >> P450 Mediated Activation to Quinones and Quinone-type Chemicals >> Methylenedioxyphenyl<br>**In vivo mutagenicity (micronucleus) alerts by ISS v2.3**<br>H-acceptor-path3-Hacceptor |
| CGA335892*<br>*Change from parent: oxidation of the pyrrole ring at the 1-position* | No genotoxic alerts | Non-mutagenic (outside) | SA26, SA34 | Same alerts as fludioxonil with additional alerts<br>**DNA binding by OASIS V1.5**<br>AN2 >> Carbamoylation after isocyanate formation >> N-Hydroxylamines<br>Radical >> Radical mechanism via ROS formation (indirect) >> N-Hydroxylamines<br>SN1 >> Nucleophilic attack after metabolic nitrenium ion formation >> N-Hydroxylamines<br>SN2 >> Acylation >> N-Hydroxylamines |
| SYN518580*<br>*Change from parent: oxidation of the pyrrole ring at the 1-, 2- and 5-positions* | No genotoxic alerts | Non-mutagenic (outside) | SA10, SA34 | Same alerts as fludioxonil with additional alerts<br>**DNA binding by OASIS V1.5**<br>AN2 >> Carbamoylation after isocyanate formation >> N-Hydroxylamines<br>Radical >> Radical mechanism via ROS formation (indirect) >> N-Hydroxylamines<br>SN1 >> Nucleophilic attack after metabolic nitrenium ion formation >> N-Hydroxylamines<br>SN2 >> Acylation >> N-Hydroxylamines<br>**Protein binding by OECD v2.3**<br>Acylation >> direct acylation involving a leaving group >> acetates |
| CGA227731**<br>*Change from parent: hydrolysis of the cyano and di-fluoro acetal group and subsequent cyclisation* | No genotoxic alerts | Probable mutagenic (within) | SA34 | **Protein binding by OECD**<br>Acylation >> direct acylation involving a leaving group >> acetates<br>**In vivo mutagenicity (micronucleus) alerts by ISS**<br>H-acceptor-path3-Hacceptor |

\* = Derek v6.0.1 and OECD QSAR Toolbox v4.2; \*\* = Derek v5.0.2 and OECD QSAR Toolbox v3.3; within = within applicability domain, outside = outside applicability domain. SA10 = $\alpha$ , $\beta$ unsaturated carbonyls; SA26 = aromatic ring-N oxide; SA34 = H-acceptor-path3-Hacceptor. Alerts presented are from the initial assessment that triggered genotoxicity testing and may not represent the latest versions of the (Q)SAR models at the time of publication. Structural change with respect to fludioxonil are indicated in *italics*.

## 2. Materials and Methods

All studies were conducted in accordance with GLP and the relevant OECD test guideline. OECD guideline versions quoted in this publication represent the guidelines to which the studies were conducted. Subsequent versions of the guidelines may have been adopted since the conduct of the studies described herein. All test substance *in vitro* concentrations and *in vivo* dose levels were selected to meet the relevant OECD test guideline requirements and are not reflective of human exposure concentrations.

### 2.1 (Q)SAR methodology

Four (Q)SAR software applications were used based on expert knowledge rules and statistical methods for the assessment of genotoxicity endpoints. These were; DEREK Nexus v.5.0.2/v.6.0.1 ((Judson *et al.,* 2003; 2013; Marchant *et al.,* 2003; Sanderson *et al.,* 1991), CAESAR v.2.1.13 (mutagenicity model) (Ferrari and Gini, 2010) and ToxTree v.2.6.13 (structural alerts for *in vivo* micronucleus formation) (Benigni *et al.,* 2009; Benigni *et al.,* 2010). Additionally, the OECD QSAR Toolbox 3.3.5.17/v.4.2 was used to assess DNA binding, protein binding and for functional group profiling. Overall, genotoxicity endpoints of *in vitro* and *in vivo* mutagenicity, chromosome damage/micronucleus formation and DNA / protein binding were considered using these tools. Due to the time-period the work described covered, different versions of the software were used as they became available. Details of the software versions used for each structure are given in Supplementary Table 1.

### 2.2 Test material

SYN518580 (4-(2,2-difluoro-1,3-benzodioxol-4-yl)-1-hydroxy-2,5-dioxopyrrole-3-carbonitrile; 96% w/w, yellow solid), CGA335892 (4-(2,2-difluoro-1,3-benzodioxol-4-yl)-1-hydroxypyrrole-3-carbonitrile; 97% w/w, beige powder) and CGA227731 (6-hydroxy-2H-chromeno[3,4-c]pyrrol-4-one; 95%w/w (bacterial reverse mutation, MNvit and *in vivo* Comet assay; brown solid), 100% w/w (bacterial reverse mutation; light pink to orange solid) and 99% w/w (TGR; brown solid)) were supplied by Syngenta.

A correction factor was applied to test material with a purity of <98%. All test substances were provided with a GLP certificate of analysis and used within the re-certificate date

### 2.3 CGA227731 *in vitro* studies

A bacterial reverse mutation test and *in vitro* micronucleus test using human lymphocytes on CGA227731 were conducted as detailed in ANSES 2023.

### 2.4 CGA335892 and SYN518580 *in vitro* studies

#### 2.4.1 Cells

*Salmonella typhimurium* LT2 bacteria of strains TA1535, TA1537, TA98 and TA100, and *Escherichia coli* WP2 strains *uvrA*/pKM101 and pKM101 were used in the bacterial reverse mutation test. The bacterial strains were obtained from the AstraZeneca Genetic Toxicology Department, Alderley Park, UK. The bacterial strains were originally sourced from Professor B N Ames, Biochemistry Department, University of California, Berkeley, California, USA (TA1535, TA1537, TA100), BioReliance Corporation, Rockville, Maryland, USA (TA98) and Dr S Venitt, Institute of Cancer Research, Royal Marsden Hospital, Surrey, UK (*E.coli* strains).

TK6 cells were obtained from European Collection of Authenticated Cell Cultures (ECACC). The ECACC states that its stocks of the TK6 cell line are obtained from the American Type Culture Collection (ATCC). TK6 cell stocks were checked for mycoplasma and confirmed to be mycoplasma free. The doubling time for TK6 cells is between 13 and 15 hours with a modal chromosome number of 47.

#### 2.4.2 Cell culture

Medium for all bacterial inoculum cultures was Nutrient Broth No.2 (Oxoid Limited) supplemented with ampicillin, as required. Bacterial test systems were plated onto minimal glucose agar medium (Vögel-Bonner agar, VWR Scientific Preparation Laboratory, Alderley Park, UK).

TK6 cells were passaged every 1 – 4 days using fresh RPMI 1640 medium containing 10% heat-inactivated horse serum (Gibco Life Technologies, UK), antibiotics and Pluronic F68 to ensure that the cell density never exceeded 1.0×10^06^ cells per mL. Following thawing cells were passaged for a minimum of 6 days prior to use. Incubation was at 37°C in a humidified atmosphere of 5% CO_2_ in air. Cells were used between passage 16 to 24.

#### 2.4.3 Reagents

Dimethyl sulphoxide (DMSO) was chosen as the vehicle for the *in vitro* studies due to its solubility properties with each of the test substances and its compatibility with the utilised test systems.

The positive control compounds for the bacterial reverse mutation tests on CGA335892 and SYN518580, 2-aminoanthracene, sodium azide, 9-aminoacridine.HCl (9-AA), 2-nitrofluorene, were obtained from Sigma and potassium dichromate was obtained from Honeywell.

For the *in vitro* micronucleus tests positive control compounds cyclophosphamide (CPA), mitomycin C (MMC), and colchicine (COL) were obtained from Sigma.

#### 2.4.4 Metabolic activation

S9 homogenate from the livers of Sprague Dawley rats treated with Aroclor 1254 was purchased from Molecular Toxicology Inc. (Boone, North Carolina, USA) via TRiNOVA Biochem GmbH, Germany (11-101). Each batch of S9 mix was tested with three pro-mutagens in the Ames test to confirm the efficacy of every batch (cyclophosphamide/TA1535, benzo(a)pyrene/TA98, 2-aminoanthracene/TA1535 and TA98). S9 homogenate was stored at −80°C and the S9 mix prepared immediately before use. The protein content of the homogenate was adjusted to 30 mg/mL with an appropriate vehicle prior to incorporation in the S9 mix.

For the bacterial reverse mutation test, the S9 mix comprised of 100 mmol/L phosphate buffer, 8 mmol/L magnesium chloride, 33 mmol/L potassium chloride, 4 mmol/L nicotinamide adenine dinucleotide phosphate, 5 mmol/L glucose-6-phosphate and 10% v/v rat liver homogenate (S9 fraction) and adjusted to a final pH of 7.4.

For the TK6 micronucleus test the S9 mix comprised of 0.25 mM nicotinamide adenine dinucleotide phosphate, 1.25 mM glucose-6-phosphate and 1% v/v Rat liver homogenate (S9 fraction).

#### 2.4.5 Bacterial reverse mutation test

The plate incorporation experiments were performed based on the methods of Maron and Ames, 1983, Venitt *et al*., 1984, Mortelmans and Zeiger, 2000 and Mortelmans and Riccio, 2000. Aliquots (100 μL) of bacterial culture from overnight cultures were mixed with solvent, positive control or test concentration (100 μL) and sodium phosphate buffer or S9 mix (500 μL) then molten 0.6% agar (2 mL) maintained at approximately 50°C and supplemented with biotin and histidine (0.05 mmol/L each) for *S. typhimurium* strains, or tryptophan (0.018 mmol/L) for the *E. coli* strains was added. The mixture was immediately poured onto minimal glucose agar plates and incubated for 3 days at 37°C. Pre-incubation experiments were performed as above except a 60-minute incubation step (37°C, 120 rpm orbital incubator) was included before the addition of the agar. For all experiments each concentration was conducted in triplicate. Revertant colonies were counted using Sorcerer® automated counter [Perceptive Instruments] adjusted appropriately to permit the optimal counting of mutant colonies or manually when necessary. Bacterial viability (≥1.8 × 10^8^ colony forming units per plate) and strain characteristics were confirmed. A minimum induction of three-fold (two-fold in *E. coli* strain WP2 pKM101) was required in the strain specific positive controls for a valid test. The test substance was considered mutagenic if there was a reproducible concentration related increase in the mean number of revertant colonies ≥2 times the concurrent solvent control.

#### 2.4.6 TK6 micronucleus test

TK6 cells were suspended at 1.5 × 10^05^ cells per mL (total 4.95 mL) in culture medium containing S9 mix where required (duplicate cultures). Aliquots (50 µL) of vehicle, test item or positive control were added to each culture and gently mixed. Short exposure; cultures incubated (3 h ±S9 mix, humidified atmosphere, 5% CO_2,_ 37°C). Cultures were then washed and medium replaced with fresh medium and further incubated until 1.5 to 2.0 population doublings were achieved in the concurrent solvent control. Continuous treatment (-S9); cultures were incubated (humidified atmosphere, 5% CO_2,_ 37°C) with the test item (or controls) until 1.5 to 2.0 population doublings were achieved in the concurrent solvent control.

Following treatment, cell densities were adjusted to 2.5×10^05^ cells per mL in fresh medium. Monolayers were prepared on slides fixed with methanol and stained with acridine orange (0.12 mg/mL in PBS, pH 7.4). Coded slides were scored blind using a fluorescence microscope (2000 mononuclear cells / test item concentration and control sample (1000 / culture where possible)). Mononucleated cells were selected if they were not overlapping and cytoplasm was clearly visible around the nuclei. MN were selected if they were <1/3 of the nucleus diameter and located within the intact cytoplasm, evenly stained, spherical with a well-defined outline and not connected to the main nucleus.

### 2.5 *In vivo* studies

All animal studies were conducted according to the Syngenta Animal Welfare Policy and are compliant with the relevant national guidelines (U.K. Animals (Scientific Procedures) Act, 1986 and associated guidelines, EU Directive 2010/63/EU for animal experiments, or the National Research Council’s Guide for the Care and Use of Laboratory Animals). The number of animals used in the experiments was chosen to ensure the minimum number of animals required under each guideline were available for genotoxicity evaluation at the end of the study period. The acceptability of each test was assessed against the criteria outlined in the corresponding OECD Guideline.

#### 2.5.1 Animals

##### Mice

Young adult male and female Crl:CD-1 mice were used in the preliminary test (3/sex/group) and *in vivo* micronucleus test (6 males/group) (Charles River (UK) Ltd., Margate, Kent, CT9 4LT, England). Mice were approximately 4-5 weeks on arrival, acclimatised for at least 11 days prior to dosing and housed in groups of three, by sex, in solid floor cages with bedding.

##### Rats

Comet assay, Crl:CD(SD) rats (Charles River UK Limited, Margate) acclimatised for a minimum of 5 days, 47-53 days old on day 1 of dosing. Similar age animals were used for the preliminary toxicity/toxicokinetic phase.

TGR range finder, male and female Fischer 344 (F344/NCrl or F344/NHsd) rats (Charles River Laboratories, Kingston, NY), 10-11 weeks of age at the initiation of dosing. This strain of rat was selected as the wild-type variant of the transgenic strain used on the main TGR study.

Main TGR study, pathogen free Fischer 344 Big Blue® transgenic male rats (Taconic nomenclature: F344-TgN(lambda/*lacI*, abbreviated as TgF344)) (BioReliance, Rockville, MD; bred by Taconic Biosciences, Inc., Germantown, NY), acclimatised for 7 days prior to first day of dosing and housed in groups of three in polycarbonate cages, 10 weeks old at the initiation of dosing.

##### Husbandry

Animals were maintained in a controlled environment (temperature between 20 to 24°C and humidity level of 30 to 70%, with a 12-hour light/dark cycle). Access to diet and water *ad libitum* throughout the study period. *Diet was* Teklad 2014C for the Crl:CD derived mice and rats and Teklad 2018C for the Fischer 344 wildtype and BigBlue® rats. Environmental enrichment was provided. Animals were randomised into treatment groups for each of the main genotoxicity studies; the weight variation did not exceed ±20% of the mean weight on Day 1 of the study.

#### 2.5.2 Micronucleus assays in the mouse with SYN518580 and CGA335892

SYN518580 and CGA335892 were formulated as suspensions in their respective vehicles (0.5 % w/v methylcellulose (1500 cps) or 0.5% w/v methylcellulose (400cp) with 0.1% v/v Tween 80).

In the preliminary and main micronucleus tests, mice were dosed twice, approximately 24 h apart (oral gavage, 10 mL/kg) with the vehicle or test substance. All animals were sacrificed approximately 48 h after the second dose.

A preliminary toxicity test was conducted in both male and female mice (3/sex/group) to establish the MTD of CGA335892 or SYN518580. Blood samples were collected via the lateral tail vein at 1, 4 and 24 h after the second dose for bioanalysis. The resultant plasma was stored frozen (≤ −70 °C), before analysis using a LC-MS-MS analytical method validated in line with the principles defined in SANTE/2020/12830 rev 1, 24 Feb 2021. The MTD for CGA335892 and SYN518580 were confirmed as 800 mg/kg/day and ≥2000 mg/kg/day respectively in both males and females.

In the main micronucleus tests, male mice (6 / group) were dosed with CGA335892 (0, 200, 400 or 800 mg/kg/day) or SYN518580 (0, 500, 1000 or 2000 mg/kg/day). 48 h post-treatment a terminal blood sample was taken for micronucleus scoring into tubes containing K_2_EDTA anticoagulant. Positive control bank samples (single oral gavage dose CPA (30 mg/kg)) were used as a positive control.

RET were scored for the presence of MN for each animal using a FACSVerse flow cytometer (min 4000 max approx. 20000) using the MicroFlow Plus Kit and as described in the instruction manual (Litron Laboratories). MN-RET data were considered acceptable if; the flow cytometry calibration standards were considered acceptable, there is no evidence of technical or human failure, the positive control samples induced a statistically significant increase in the MN-RET and the vehicle control was considered acceptable for inclusion in the HCD.

All MN-RET data was included in the analysis (no exclusions). MN-RET evaluation was conducted by flow cytometry, blinding of the study samples is not required.

### 2.6 CGA227731 *in vivo* studies

CGA227731 was formulated in 1% methylcellulose for all *in vivo* tests. Preparations of CGA227731 in this vehicle were shown to be accurate, homogenesis and stable for the duration of the use of the formulated test substance using a validated analytical method.

#### 2.6.1 Comet assay in the rat with CGA227731

Preliminary test; to establish the MTD male and female rats (2/sex/group) were administered two doses (0, 24 h) of CGA227731 by oral gavage (20 mL/kg) then sacrificed (48 h). For the toxicokinetic evaluation, blood samples from male animals dosed once (2000 mg/kg/day) were taken (0.5, 1, 2, 4, 8 and 24 h), Tmax was established to be approximately 1 h.

Main Comet test; male rats (6/group) received two doses of CGA227731 (0, 500, 1000 and 2000 mg/kg/day; oral gavage; 20 mL/kg) at 0 and 23 h then sacrificed (24 h). A positive control group of 3 male rats were administered a single dose of EMS (200 mg/kg (Sigma) in water (10 mL/kg)) 3 h prior to sacrifice. Liver and duodenum samples were collected, processed and analysed for Comets as outlined in ANSES 2023 (p438-9). Coded slides were used to evaluate all treated animals. The Comet data was considered acceptable if the concurrent vehicle control is considered acceptable for addition to the HCD and the positive control induced statistically significant increase in %TI and is comparable to positive control HCD. Data from all slides and animals were included in the analysis (no exclusions).

#### 2.6.2 TGR in the BigBlue® rat with CGA227731

Preliminary test: CGA227731 was administered at 0 (vehicle control), 500 or 1000 mg/kg/day to male and female F344 wildtype rats (5/sex/group) by oral gavage (10 mL/kg) for 14 consecutive days to establish the MTD in both sexes.

Transgenic rodent mutation study; CGA227731 was administered to male BigBlue® transgenic rats by oral gavage (10 mL/kg) at 0 (vehicle control), 250, 500 and 1000 mg/kg/day (6/group; 7 in top dose) once daily for 28 days. Rats were sacrificed 3 days following the final treatment. Samples of liver, duodenum and bone marrow were flash frozen in liquid nitrogen during necropsy then stored at −80°C until *cII* mutation analysis.

DNA isolated from previous positive control animals [F344 Big Blue® transgenic male rats exposed to ENU (20 mg/kg/day) by oral gavage, on 6 occasions between study days 1 and 28 and necropsied on Study Day 31], was used as a packaging positive control. The purpose of the packaging positive control is to confirm that the efficacy of the packaging reaction.

Initially, 5 animals/group were selected for evaluation of gene mutation. A supplementary assessment of mutant frequency was conducted using the sixth animal per dose group for liver.

##### 2.6.2.1 Extraction and packaging of genomic DNA

DNA was extracted from liver, bone marrow and duodenum samples following the Agilent RecoverEase™ Methods (Agilent 2018). Following dialysis, the DNA samples were stored at 2-8°C until analyzed for *cII* mutations.

Isolated DNA was processed using Packaging Reaction Mix (PRM), (New York University, New York, NY). Methods were based on Agilent 2015a, 2015b and Guttenplan *et al.,* 2004.

##### 2.6.2.2 Plating of packaging DNA

After inoculation of overnight *E. coli G1250* suspension cultures (strain developed at NIH, licensed to Agilent for use with Big Blue® *cII* selectable mutation assays), packaged phage were adsorbed onto the bacterial suspension cultures for at least 30 min, molten top agar was added and the cells were plated onto bottom agar plates. Packaged phage were incubated overnight (37 ± 2.0°C), then scored for plaque formation and titer determination; *cII* mutant selection plates were incubated for two days (24 ± 0.5°C), then scored for mutant plaques after incubation.

At least 125,000 phage were evaluated from at least 2 packagings. Titer and mutant frequency were calculated. Mutant frequency was calculated by dividing the total number of mutants per sample by the total number of phage evaluated per sample and normalizing to mutants per million phage evaluated (× 10^-6^). The assumption was that each plaque represents one phage that had infected one *E. coli* cell. Titer and mutant counts from individual packagings were totalled and the final mutant frequency calculated at the animal level. The data from one packaging (liver only, 1 animal at top dose) was excluded as the packaging positive control did not have an elevated MF and therefore the acceptance criteria were not met. As the minimum number of phage screened was exceeded in all animals this had no impact on the validity of the study data. The data from all other packagings met acceptance criteria and were included in the analysis. Data from all animals analysed for MF are included in the statistical analysis. For the liver, the initial data from the first 5 animals is also presented. Samples were labelled with animal number/tissue during packaging/scoring.

### 2.7 Statistical analysis

TK6 micronucleus test; the number of MN analysed from 2000 mononuclear cells for each selected test substance concentration was compared with that from the concurrent solvent control. Pair-wise statistical analysis employing a one-sided Fisher’s Exact test were used to evaluate statistical significance (p <0.05). A linear trend test was employed (Cochran Armitage) in order to confirm a dose-response (p <0.05).

The individual animal is the experimental unit in all *in vivo* genotoxicity tests reported. Calculations for treated groups versus the concurrent vehicle control and positive control versus vehicle control were conducted separately for each test.

For *in vivo* % RET data, the treated groups and positive control were compared with the vehicle control using a one-tailed Wilcoxon Rank Sum test. A Jonckheere trend test was conducted for dose related response.

MN-RET data was analysed using a Poisson model with a log link function using GENMOD in SAS. The effect was analysed using two statistical approaches; a linear trend test and pairwise comparison to the vehicle control (one sided Likelihood ratio test). All the tests were one-sided, a decreasing trend being not of interest. In case of decrease and calculated p value < 0.5, the final p value was estimated as 1-[p value]. Overdispersion of the data was estimated by the deviance of the model divided by the associated degrees of freedom. Where overdispersion was detected (dispersion parameter >1), a correction was applied on the model. Previous statistical evaluations have demonstrated that *in vivo* designs for MN based on OECD 474 (2016), (namely n=5, minimum of 4000 cells analysed /animal) provide power to detect 2 to 3 fold change with 80% power when MN-RET >0.1% (OECD 2017a).

Comet assay; the median percent tail intensity (%TI) was measured for three slides per animal, the mean of median animal %TI (experimental unit) was then derived followed by the group mean of median %TI.

Initially, Bartlett’s test for variance homogeneity (F1 test) was applied. If non-significant (p≥0.01), then parametric analysis was conducted using the Williams test (CGA227731 treated groups) or t-test (positive control group). If Bartlett’s test was significant (p>0.01), then logarithmic transformations were performed (if a parameter contained any zero values, 0.001 was added to all values for that parameter before applying log transformation). If Bartlett’s test was non-significant (p≥0.01) following logarithmic transformation, then the Williams test was performed on transformed data. The %TI based on the slide mean is also presented but no statistical analysis is conducted on this parameter as the mean of median tail intensity the most appropriate endpoint for genotoxicity evaluation (OECD 489, 2016). The Comet design was selected to follow the guideline and allow a 2.5-fold increase in %TI to be detected with 80% probability as described in Smith *et al*. (2008).

For the TGR, statistical analysis was performed using Minitab® 18.1. The individual log_10_ transformed MF data were evaluated using 1-Way ANOVA followed by Dunnet’s test. The ANOVA was considered appropriate as tests for normality (Ryan-Joiner) and equal variance (Levene’s test) passed (p≥0.05) in each case. The TGR is designed to detect a doubling of MF (OECD 488).

For all tests, probability values of less than 5 % (p<0.05) were regarded as statistically significant. The test performed for each variable is described in the footnote of the relevant summary table. Confidence intervals were calculated for the purposes of this publication.

## 3. Results

### 3.1 (Q)SAR

The results of the multi-(Q)SAR assessment of fludioxonil and the three dietary metabolites are summarised in Table 1. Briefly, no genotoxicity alerts were triggered for any of the structures in DEREK, for CAESAR only CGA227731 was within the applicability domain of the model and predicted to be mutagenic. Toxtree structural alerts for *in vivo* micronucleus formation were identified for all structures. In the OECD QSAR Toolbox non-specific mechanistic DNA binding alerts were triggered for all structures, CGA227731 and SYN518580 additionally triggered a non-specific mechanistic protein binding alert. Following this assessment, each of the metabolites was identified as the exemplar compound within its own group. Bacterial reverse mutation and *in vitro* micronucleus studies were therefore conducted on each metabolite as well as relevant higher tier *in vivo* follow-up studies where indicated.

### 3.2 CGA335892 *in vitro* studies

A bacterial reverse mutation test with CGA335892 (OECD 471, 1997) at up to cytotoxic concentrations showed no increases in revertants in any of the strains tested (*S. typhimurium* TA1535, TA1537, TA98 and TA100, and *E. coli* WP2 *uvrA*/pKM101 and pKM101) in either a plate incorporation or a pre-incubation test in both the presence and absence of metabolic activation (see Supplementary Tables 2 to 5 and Supplementary Table 12). The strain specific positive controls showed distinct increases in revertants demonstrating the sensitivity of the test system and the efficacy of the S9 mix. CGA335892 was therefore concluded to be negative in the bacterial reverse mutation test.

CGA335892 was tested up to cytotoxic concentrations (55%±5%) in an *in vitro* TK6 micronucleus test (OECD 487, 2016; 3 h ± S9 and 24 h -S9 exposures) resulting in statistically significant increases in micronuclei in all exposure conditions (see Figure 1). In the 3 h ± S9 exposures the top concentration evaluated (263 µg/mL) had MN frequencies outside the 95% confidence limits of the vehicle HCD; the trend test for the exposure with metabolic activation was statistically significant. In the continuous exposure, the MN frequencies of all CGA335892 concentrations was outside the 95% confidence limits and statistically significant (Fishers exact test p<0.001). The trend test was also statistically significant (Cochran Armitage trend test; p<0.0001). In this condition, large increases in micronuclei were observed even at concentrations with minimal cytotoxicity; a concentration with 15.45% cytotoxicity had a micronuclei frequency of 101/2000 cells (vehicle control of 21/2000 cells). CGA335892 was concluded to be clearly positive in the *in vitro* micronucleus test. In order to understand the biological relevance of this finding an *in vivo* micronucleus test in the mouse was conducted.

**Figure 1:**
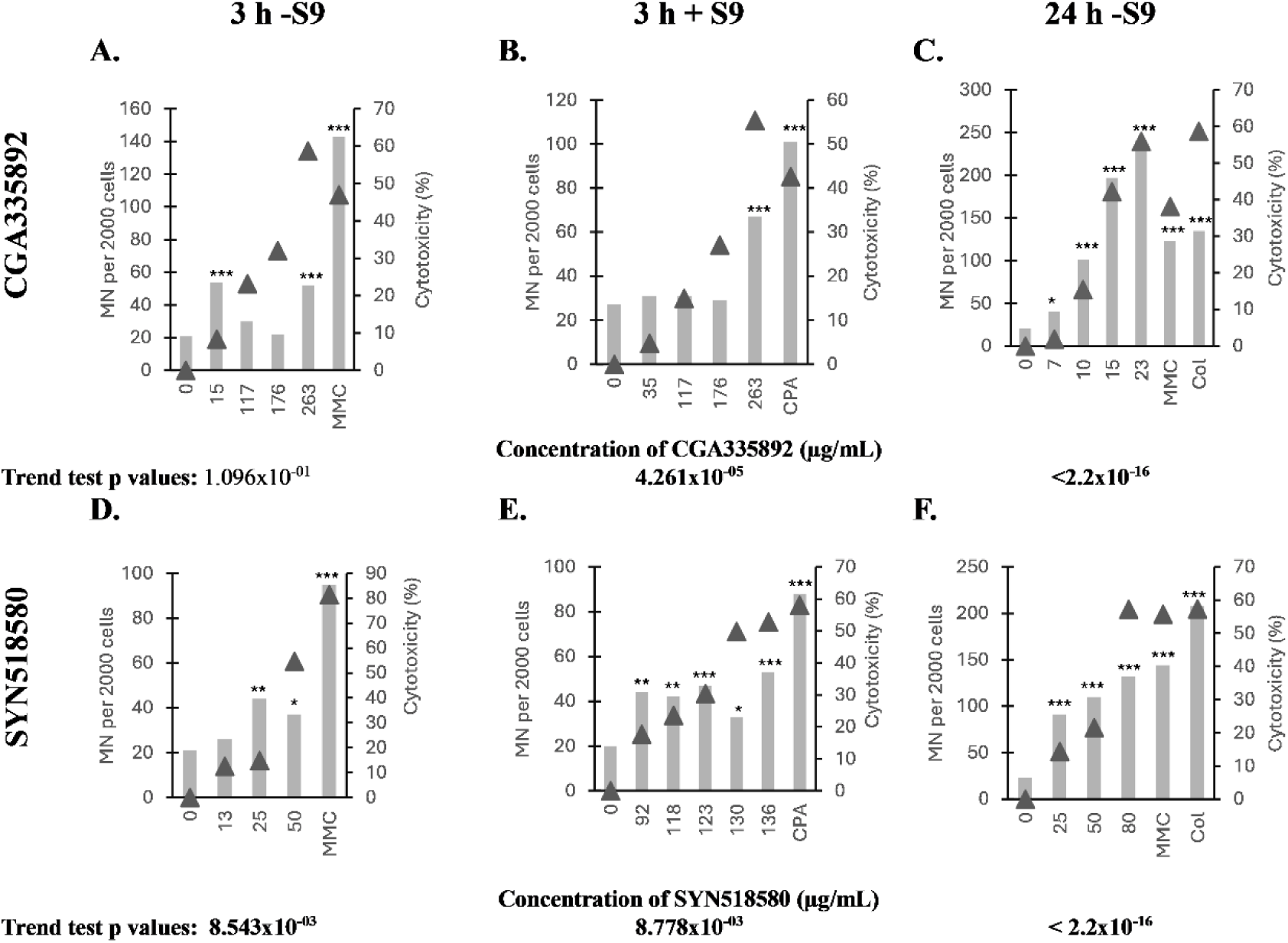
*In vitro* micronucleus results on (A-C) CGA335892 and (D-F) SYN518580 in TK6 cells. (3 h ±S9 and 24 h -S9). The number of micronucleated cells in 2000 mononucleated cells is presented (bar chart). Cytotoxicity (Δ) was calculated from relative population doubling. COL; colchicine at 7.5 ng/mL (24 h), MMC; mitomycin C at 50 ng/mL(3 h -S9) or 30 ng/mL (24 h), CPA: cyclophosphamide at 4 µg/mL (3h+S9). Fisher exact test *p<0.05; **p<0.01; ***p<0.001 versus concurrent vehicle control (1% DMSO; 0 µg/mL). Trend tests (Cochran-Armitage) using vehicle control and all test substance concentrations were conducted (values in bold p<0.05; significant). Concentrations are rounded to the nearest µg.

### 3.3 CGA335892 *in vivo* studies

*In vivo* MN test; in a preliminary toxicity test the maximum tolerated dose (MTD) was established (800 mg/kg/day) in male and female mice. A higher dose (1250 mg/kg/day) resulted in excessive toxicity and clear clinical observations consistent with systemic exposure. As there was no difference in the MTD between the sexes, the main study was conducted in males only at 0, 200, 400 and 800 mg/kg/day.

CGA335892 was quantified in all plasma samples collected on the preliminary test using a validated LC-MS/MS method (see Supplementary Table 13). The clinical signs observed in the preliminary test and the detection of CGA335892 in the plasma are lines of evidence that the bone marrow was exposed to CGA335892 in the micronucleus test (EFSA 2017).

In the mouse micronucleus test (OECD 474, 2016), the mean MN-RET frequency of all CGA335892 treated groups remained well within the 95% control limit of the vehicle HCD. At the intermediate dose of 400 mg/kg/day, a statistically significant increase in MN-RET frequency was observed (0.28% versus 0.21%; pairwise analysis using Poisson Model, p=0.0467; Table 2). As there was no statistically significant increase in MN frequency at the top dose, no statistically significant dose related trend (Poisson model trend analysis p=0.0957) and the mean MN-RET frequency of all CGA335892 treated groups remained well within the 95% control limits this statistical significance was not considered to be of biological relevance. It was concluded that CGA335892 was neither clastogenic nor aneugenic *in vivo* and was negative in this *in vivo* micronucleus test.

**Table 2:** Summary of *In vivo* Micronucleus Results with CGA335892 and SYN518580.

| Dose level (mg/kg bw/day) | N. | Mean Total RET evaluated per animal | %RET | Mean % MN-RET ±SD | Confidence interval MN-RET (95%) | P value | HCD 95% Control limit (± 2SD) | Trend test p value |
| --- | --- | --- | --- | --- | --- | --- | --- | --- |
| **CGA335892** |  |  |  |  |  |  |  |  |
| Vehicle# | 6 | 19626.33 | 1.39 | 0.21±0.05 | 0.17 - 0.25 | - | 0.09 - 0.34% | 0.0957 |
| 200 | 6 | 19508.17 | 1.67 | 0.20±0.05 | 0.16 - 0.24 | 0.6188 | | |
| 400/ | 6 | 19646.33 | 1.83 | **0.28±0.09** | 0.21 - 0.35 | **0.0467** |  |  |
| 800 | 6 | 20261.67 | 1.67 | 0.24±0.04 | 0.21 - 0.27 | 0.0943 | | |
| CPA | 6 | 18708.50 | **0.42\*** | **0.72±0.12** | 0.62 - 0.82 | **0.0000** | 0.40-1.01 | n/a |
| **SYN518580** |  |  |  |  |  |  |  |  |
| Vehicle# | 6 | 19432.33 | 2.26 | 0.16±0.03 | 0.14 - 0.18 | - | 0.09- 0.34% | **0.018** |
| 500 | 6 | 20216.17 | 2.45 | 0.14±0.05 | 0.10 - 0.18 | 0.8188 | | |
| 1000 | 5 | 20597.00 | 2.75 | 0.18±0.04 | 0.15 - 0.22 | 0.1944 | | |
| 2000 | 5 | 19618.80 | 3.32 | 0.22±0.1 | 0.13 - 0.31 | 0.0548 | | |
| CPA | 6 | 15042.50 | **0.65\*** | **0.79±0.22** | 0.61 - 0.97 | **<0.0001** | 0.40-1.01 | n/a |

### 3.4 SYN518580 *in vitro* studies

A bacterial reverse mutation test (OECD 471, 1997) with SYN518580 at up to cytotoxic concentrations showed no relevant reproducible increases in revertants in any of the strains tested (*S. typhimurium* TA1535, TA1537, TA98 and TA100, and *E. coli* WP2 *uvrA*/pKM101 and pKM101) in either a plate incorporation or a pre-incubation test in both the presence and absence of metabolic activation (Supplementary Tables 6 to 12).

In the plate incorporation test, there was an isolated marginal increase (2.1-fold over the concurrent control, TA1535, 500 µg/plate) in the absence of metabolic activation, higher concentrations could not be evaluated due to excessive cytotoxicity and the presence of pseudorevertants. This condition was repeated using a modified concentration spacing to test the reproducibility of this marginal increase. In the repeat test, there were no increases ≥2 fold at any concentration (maximum fold change 1.5; Supplementary Table 10). As the marginal increase observed in the first experiment was not reproduced in the second experiment the isolated increase in the first test was considered to be anomalous and not biologically significant.

In the pre-incubation test strain WP2 pKM101 with metabolic activation was repeated due to the solvent control being below the acceptable minimum range and not testing to a cytotoxic or regulatory maximum level (5000 µg/plate). The repeat experiment was conducted using an adjusted concentration level and showed there were no relevant increases in revertant colonies in this condition (Supplementary Table 11). There were no relevant reproducible increases in revertants in any condition, SYN518580 was concluded to be negative (i.e. non-mutagenic) in the bacterial reverse mutation test.

SYN518580 was tested up to cytotoxic concentrations (55%±5%) in an *in vitro* TK6 micronucleus test (OECD 487; 2016; 3 h ± S9 and 24 h -S9 exposures (see Figure 1)). Statistically significant increases in micronucleated mononucleated (MM) cells were observed in each of the three treated conditions at almost all concentrations of SYN518580 evaluated for micronuclei (Fishers exact test; p<0.05). The increases were outside the laboratory’s historical control 95% confidence limits at multiple concentrations and the concentration related trend was statistically significant in each condition (Cochran-Armitage trend test p<0.001). The *in vitro* micronucleus assay on SYN518580 was therefore concluded to be clearly positive in both the presence and absence of metabolic activation. SYN518580 is therefore considered to be clastogenic and/or aneugenic *in vitro*. In order to understand the biological relevance of this finding an *in vivo* micronucleus assay in the mouse was conducted on SYN518580.

### 3.5 SYN518580 *in vivo* studies

*In vivo* MN test; in a preliminary toxicity test, SYN518580 was tolerated up to the regulatory limit dose of 2000 mg/kg/day in male and female mice. There were no clinical observations in either sex. Since there was no difference in the tolerability between the sexes, the main micronucleus test was conducted in males only at 0, 500, 1000 and 2000 mg/kg/day.

SYN518580 was quantified in all plasma samples collected on the preliminary test using a validated analytical method (see Supplementary Table 15). Since the bone marrow is a highly perfused tissue this demonstrates that the bone marrow is exposed to SYN518580 following oral gavage dosing (lines of evidence EFSA 2017).

In the mouse micronucleus test (OECD 474; 2016), SYN518580 was administered (6 mice/ group, 0, 500, 1000 or 2000 mg/kg/day; 0 and 24 h) and animals were sacrificed 48 h following the last dose. At least five animals per group were analysed for MN-RET. Clinical observations (including 1000 mg/kg/day piloerection; 2000 mg/kg/day, piloerection, partially closed eyes and decreased activity) provide further lines of evidence of systemic and therefore bone marrow exposure occurred (EFSA 2017). They also indicate the assay was performed to the MTD.

There were no statistically significant increases in MN-RET and the frequency of MN-RET of all treated groups remained well within the laboratory’s 95% control limit for the vehicle control (0.09-0.34%) (see Table 2 and Supplementary Table 17). There was a statistically significant dose related trend (Poisson Model; p=0.0108). All animals had MN-RET frequencies within the observed (min-max) range of the laboratory’s historical control data and all, except one high dose animal (0.36%), were within the 95% control limit of the vehicle HCD.

Since no MN-RET values were statistically significant versus the concurrent vehicle control and all mean and individual animal MN-RET frequencies were within the vehicle HCD range (Supplementary Tables 16 and 17) the *in vivo* micronucleus test was concluded to be negative and that SYN518580 was neither clastogenic nor aneugenic *in vivo*.

### 3.6 CGA227731 *in vitro* studies

In an initial bacterial reverse mutation assay (OECD 471, 1997) with CGA227731 (95% purity) there were concentration related increases in the number of revertants above the threshold for a positive response in TA1537 in the presence and absence of metabolic activation (see Figure 2). There were no relevant increases in any of the other strains tested (TA98, TA100, TA1535 and WP2 uvrA (pKM101)). An additional test using ultra-pure (100%) material confirmed that the observed response in TA1537 was due to CGA227731 (data not shown). CGA227731 was concluded to be positive in the bacterial reverse mutation test.

**Figure 2:**
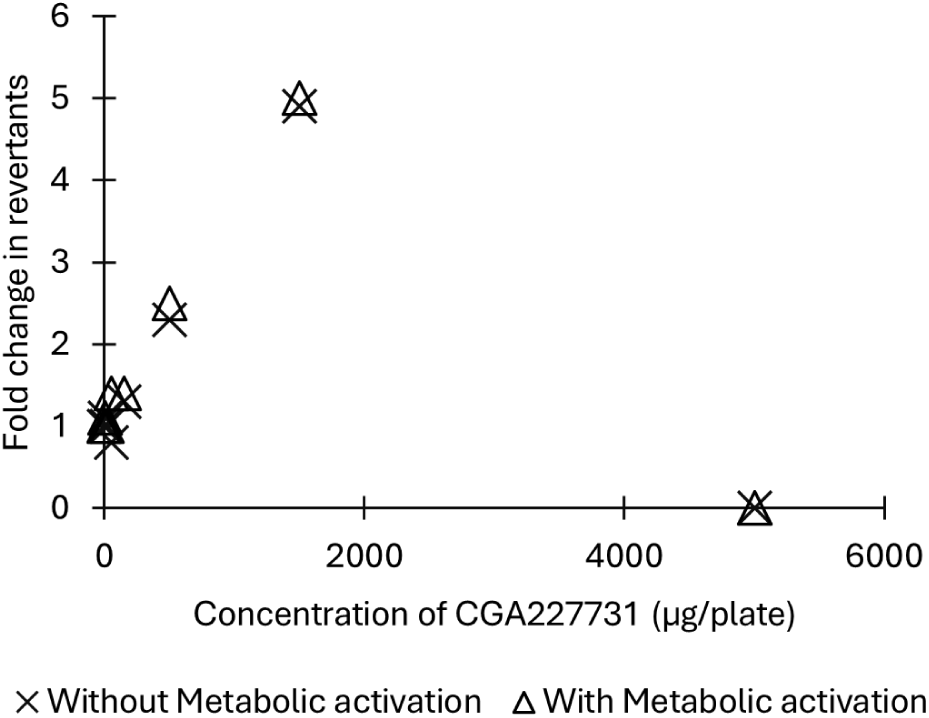
CGA227731 fold increase in revertants in TA1537. Positive control agents (9-AA without S9; 50 µg/plate and benzo[a]pyrene with S9; 5 µg/plate)) showed distinct increases (>6 fold) in the number of revertants confirming the test system sensitivity and the efficacy of the S9 mix.

In an *in vitro* micronucleus assay human lymphocytes were treated with CGA227731 (OECD 487, 2016; 3 h ±S9; 20 h - S9).

In all three treatment conditions, there were no statistically significant increases in MN-BN (Williams test; p>0.05), the frequency of MN-BN were within the 95% control limits of the vehicle HCD and there was no concentration related increase in MN-BN frequency (linear contrast; p>0.05) (see Figure 3). In the 3 h treatments in the presence and absence of S9-mix, precipitation was observed at the end of the treatment at the highest concentration scored for micronuclei (300 µg/mL). In the 20 h -S9 treatment, CGA227731 was tested up to a cytotoxic concentration of 90 µg/mL (57.5% cytotoxicity). Positive controls in each experiment showed statistically significant increases in MN-BN demonstrating the sensitivity of the test system to detect a known clastogen and aneugen. The efficacy of the S9-mix was demonstrated against a known clastogen (CPA). CGA227731 did not induce any increases in the frequency of MN-BN and was concluded to be clearly negative in the *in vitro* micronucleus assay. CGA227731 is therefore not clastogenic and not aneugenic *in vitro*.

**Figure 3:**
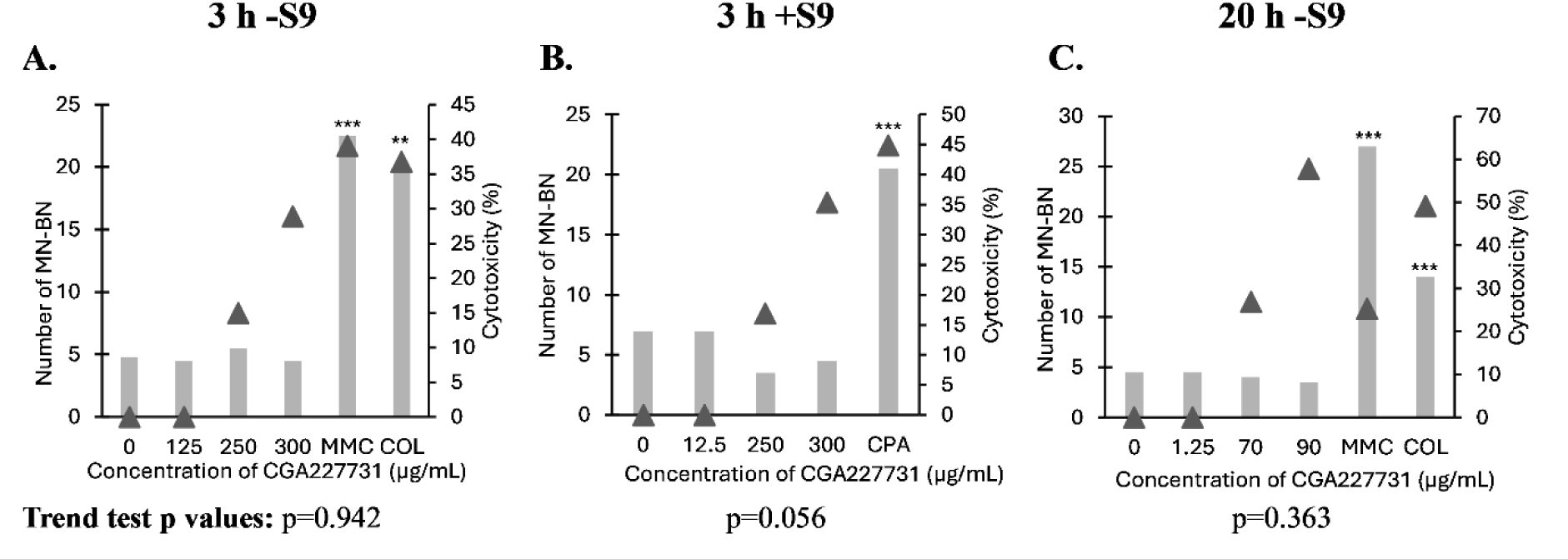
CGA227731 *in vitro* micronucleus assay in human lymphocytes: 3 h treatment in the A. absence and B. presence of metabolic activation. C. 20 h treatment in the absence of metabolic activation. The mean number of MN-BN in 1000 BN is presented (bar chart). Each concentration had replicate cultures (2000 BN scored / concentration). Cytotoxicity (Δ) was measured as cytostasis calculated from the CBPI. COL; colchicine at 0.07 µg/mL (3 h) or 0.015 µg/mL (20 h), MMC; mitomycin C at 0.3 µg/mL (3 h) or 0.1 µg/mL (20 h), CPA: cyclophosphamide at 10 µg/mL. Positive controls t-test; ** p<0.01; *** p<0.001. CGA227731 treated concentrations; Williams test p>0.05 (non-significant). Trend tests were conducted by linear contrast by group number using control and all test substance concentrations.

### 3.7 CGA227731 *in vivo* studies

To address the *in vivo* relevance of the positive bacterial reverse mutation test, an *in vivo* Comet assay (OECD 489, 2016) in the rat was conducted (EFSA 2016). The Comet assay detects DNA damage that may result in gene mutation or chromosomal damage if left unrepaired.

The response in TA1537 was observed in both the presence and absence of metabolic activation so the *in vivo* Comet assay was conducted in the liver and the duodenum (site of contact tissue). A toxicokinetic investigation conducted by oral gavage dosing prior to the start of the main Comet assay showed; the rat is systemically exposed to CGA227731, Tmax was approximately 1 h post dose, and identified the appropriate sampling time for the Comet assay.

In accordance with the recommendations by Dertinger *et.al.* (2023) the comparison to the HCD data is made on a like for like basis (i.e. individual animal data on the study are compared against the HCD generated from individual animals (iHCD) and the group mean data are compared against the HCD generated using the study means (mHCD))

In the duodenum, the were no statistically significant increases in median percent tail intensity (%TI) following administration of CGA227731 at up to the limit dose of 2000 mg/kg/day (Williams test; p>0.05). The group median %TI values for all treatment groups were within the vehicle mHCD 95% confidence limits (Table 3 and Supplementary Table 19).

**Table 3:** Summary of *In vivo* Comet Results with CGA227731.

| CGA227731<br>(mg/kg/day) | Group mean<br>tail intensity (%<br>±SD) | Group mean of<br>median tail<br>intensity (%<br>±SD) | Confidence<br>interval group<br>mean of median<br>tail intensity<br>(95%) | Median %TI<br>95% confidence<br>limit mHCD | Group mean<br>number of<br>hedgehog<br>cells (±SD) |
| --- | --- | --- | --- | --- | --- |
| **Duodenum** |  |  |  |  |  |
| Vehicle | 4.83 ±0.9 | 1.88 ±0.9 | 1.16 - 2.60 | 0.0 – 2.4 | 4.5 ±3.0 |
| 500 | 4.38 ±0.8 | 1.91 ± 0.9 | 1.19 - 2.63 |  | 3.5 ±2.3 |
| 1000 | 4.99 ±0.9 | 1.62 ± 0.9 | 0.90 - 2.34 |  | 5.5 ±2.3 |
| 2000 | 4.91 ±0.6 | 2.05 ± 0.7 | 1.49 - 2.61 |  | 7.7 ±3.8 |
| 200 mg/kg EMS | 29.54 ±2.4 | **26.18±2.9\*\*\*<sup>+</sup>** | 23.86 - 28.50 | 29.9 – 65.1 | 0.0 ±0.0 |
| **Liver##** |  |  |  |  |  |
| Vehicle | 1.76 ±0.4 | 0.17±0.1<sup>L</sup> | 0.09 - 0.25 | 0.1 – 1.0 | 0.0 ±0.0 |
| 500 | 1.50 ±0.4 | 0.18±0.1<sup>L</sup> | 0.10 - 0.26 |  | 0.2 ±0.4 |
| 1000 | 1.78 ±0.3 | 0.24± 0.2<sup>L</sup> | 0.08 - 0.40 |  | 2.8 ±1.6 |
| 2000 | 2.56±0.8 | **1.03 ± 0.6 \*\*\*<sup>L</sup>** | 0.55 - 1.51 |  | 4.8 ±4.1 |
| 200 mg/kg EMS | 20.54 ±0.7 | **18.22 ±1.3 \*\*\*<sup>L+</sup>** | 17.18 - 19.26 | 22.7 – 61.9 | 0.0 ±0.0 |

Vehicle = 1% methylcellulose; EMS = ethyl methanesulphonate (positive control), dosed once approximately 3 h prior to termination. Statistical analysis performed on median tail intensity values only. Results of statistical analysis using the appropriate nonparametric method of analysis based on permutation (one-sided probabilities): L – Analysis performed upon logarithmically transformed data. p values for comparisons with control using Williams' test, unless indicated otherwise (+ t-test). ## Result in the liver are the combined values following additional analysis.
\*\*\* p<0.001 (bold; significant); otherwise p>0.05 (non-significant)

In the liver, there was a statistically significant increase in the median %TI in rats administered CGA227731 at the top dose of 2000 mg/kg/day versus the concurrent vehicle control (Williams test; p<0.001). The group median %TI for the top dose (1.02%) was fractionally outside the 95% confidence limits of the mHCD (1.0%). However, the individual animal and group median %TI were all within the absolute range of the respective HCD. An additional 150 cells were analysed to resolve the observed increase at 2000 mg/kg/day. The result of the combined analysis was concordant with the initial assessment (1.03% TI at 2000 mg/kg/day; see Table 3 and Supplementary Tables 18 and 19). A dose related increase in the number of hedgehog cells was also observed in the liver and following histopathological examination an increase in the number of hepatocellular mitotic figures, characteristic of target tissue toxicity, was observed in two animals at 2000 mg/kg/day (data not shown).

Given the minimal increase in %TI (<1% absolute increase in %TI, within HCD), the evidence of target tissue toxicity and hedgehogs the *in vivo* Comet assay was concluded to be negative in the liver and duodenum.

From the available *in vitro* and *in vivo* data the authors were of the opinion that CGA227731 was not genotoxic. However, The Pesticide Peer Review Experts Meeting (5 April 2019) formed a different opinion and concluded that based on the bacterial reverse mutation assay response in TA1537 and the increase in %TI observed in the Comet assay, despite evidence of tissue toxicity and hedgehogs, that CGA227731 was genotoxic *in vivo*.

To address the concerns raised by The Pesticide Peer Review Experts Meeting (5 April 2019) regarding the potential *in vivo* mutagenicity of CGA227731 a second higher tier *in vivo* study addressing the specific apical endpoint of gene mutation was conducted. An *in vivo* transgenic rodent gene mutation assay on CGA227731 was performed in BigBlue® rats (OECD 488, 2020). A preliminary test in wildtype rats of the same background strain showed that CGA227731 was tolerated up to the limit dose of 1000 mg/kg/day over 14 days in both sexes. The main study was therefore conducted in males only.

Male BigBlue® rats were administered CGA227731 (0, 250, 500 or 1000 mg/kg/day) once daily for 28 days then sacrificed three days after the last dose. The liver, duodenum and bone marrow were selected for *cII* mutant frequency analysis. In all tissues evaluated there were no relevant increases in *cII* mutation frequency following treatment with CGA227731 at up to the limit dose of 1000 mg/kg/day (Table 4).

**Table 4:** In vivo Mutation at cII locus in BigBlue® Rats.

| CGA227731<br>(mg/kg/day) | Mutant Frequency ± SD ($\times 10^{-6}$)<br>[Confidence Interval- 95%] | | | |
| --- | --- | --- | --- | --- |
|  | Liver<br>(n=5) | Liver<br>(n=6*) | Duodenum<br>(n=5) | Bone Marrow<br>(n=5) |
| Vehicle | 64.0±20.2<br>[46.29 - 81.71] | 61.8±18.9<br>[46.68 - 76.92] | 50.7±14.5<br>[37.99 - 63.41] | 32.6±15.1<br>[19.36 - 45.84] |
| 250 | 59.5±23.7<br>[38.73 - 80.27] | 57.3±21.0<br>[40.50 - 74.10] | 34.2±12.8<br>[22.98 - 45.42] | 40.2±23.0<br>[20.04 - 60.36] |
| 500 | 53.3±14.5<br>[40.59 - 66.01] | 51.6±13.6<br>[40.72 - 62.48] | 30.5±4.4<br>[26.64 - 34.36] | 47.0±22.1<br>[27.63 - 66.37] |
| 1000 | 59.9±15.0<br>[46.75 - 73.05] | 65.9±19.9<br>[49.98 - 81.82] | 35.4±15.3<br>[21.99 - 48.81] | 39.3±8.9<br>[31.50 - 47.10] |
| ENU# | 303.5±104.0<br>[212.34 - 394.66] | **289.9±85.8\*\*\***<br>[214.69 - 365.11] | **810.7±116.4\*\***<br>[708.67 - 912.73] | **343.1±108.7\*\***<br>[247.82 - 438.38] |
| **Vehicle mHCD** |  |  |  |  |
| N | 40 |  | 9 | 23 |
| Mean ± SD | 41.5 ± 11.7 | | 41.0 ± 11.5 | 29.4 ± 5.7 |
| 95% vehicle control limit | 18.1 - 64.9 |  | 18.0 - 64.0 | 18.0 - 40.8 |
| Range | 25.2 - 76.2 |  | 27.4 - 61.9 | 19.3 - 38.7 |

\* = Values reported for N=6 in Groups 1-4. \*\* = Statistically significant (1-Way ANOVA, $p < 0.001$ ). # Packaging positive controls from animals treated with *N*-ethyl-*N*-nitrosourea (ENU) (20 mg/kg/day). *N*-ethyl-*N*-nitrosourea (ENU) by oral gavage, at 20 mg/kg/day once daily on Study Days 1, 2, 3, 10, 17 and 24 and necropsied on Study Day 31 (liver) or once daily on Study Days 1, 2, 3, 12, 19 and 26 and necropsied on Study Day 31, (duodenum and bone marrow). <sup>a</sup> A sixth animal was not packaged for the positive control group. DNA from the first five animals was used for this group when packaging the sixth animal in Groups 1-4 resulting in one group mean after the initial packaging and a new group mean after additional packaging as part of analysing the sixth animal per Groups 1 to 4. Statistical analysis for liver (n=5) are not presented

In the liver, initially DNA from 5 animals was packaged and analysed for mutation frequency, in this initial analysis there were no statistically significant increases in mutation frequency at any dose level and all group mean mutation frequencies were within the 95% control limits of the mHCD. Due to some variability in the individual animal mutation frequency in both vehicle and treated groups, an additional animal DNA was packaged and analysed to increase the statistical power of the study in this tissue. This combined analysis confirmed there were no statistically significant increases in mutation frequency. The MF for all individual animals treated with vehicle or CGA227731 were within the observed range of the iHCD; one vehicle control animal and two animals receiving 1000 mg/kg/day had MF outside the 95% control limit (see Supplementary Tables 20 and 21). The mean MF for the top dose was also marginally outside the 95% control limit (65.9 vs. 64.9 × 10^-6^) following the inclusion of the sixth animal. As there was no statistically significant difference between any treatment group and the vehicle and the test substance data was comparable to the HCD, CGA227731 was concluded to be negative in this tissue.

In the duodenum, there were no statistically significant increases in MF versus the concurrent vehicle control as the MF was below the concurrent vehicle control at all treated doses. All mean and individual MF were within 95% control limit of the respective HCD confirming a negative result in this tissue.

In the bone marrow, the mean MF of the intermediate group of 500 mg/kg/day was outside the 95% control limit and range of the mHCD however, the MF was not statistically different from the vehicle control. There were no statistically significant increases at any dose level and the mean MF values for the other two CGA227731 dose groups were within the 95% control mHCD limits. The MF for all animals were within the range of the iHCD; one animal in both the vehicle control and low dose group and three animals in the intermediate dose group had MF above the 95% control limits iHCD. These data confirmed a negative result in this tissue.

Tissues from animals that were treated with ENU on a separate occasion were included in all packagings to confirm the efficacy of the packaging. In each tissue, a statistically significant increase in MF was observed in ENU treated tissues compared to the vehicle control confirming the efficacy of the packaging and the validity of the study. Since there were no relevant increases in any tissue evaluated CGA227731 was concluded to be non-mutagenic *in vivo*.

## 4. Discussion

The evaluation of the genotoxic potential of dietary metabolites for an active substance is a tiered process. The first step is to compile a list of all the dietary metabolites and gather information (structure, SMILES, genotoxicity data) on the dietary metabolites and any structurally related compounds (e.g. parent, ground water metabolites, other active substances or metabolites from the same chemical class). The next phase of the assessment is to run the (Q)SAR including at least one statistical based model and one knowledge-based model to afford a consensus output which should be accompanied by expert review. In the read-across process compounds are placed into groups based on their structural similarity and the alerts present, then an “exemplar” chemical structure is identified for each group. The exemplar must cover all the alerts for each structure in the group and have sufficient structural similarity to the other compounds in the grouping. Depending on the data richness in some cases, a ‘2 to 1’ read-across may be possible to cover all the alerts for a particular compound to reduce the need for further unnecessary testing. For any groups where not all the required genotoxicity endpoints are covered; then the appropriate *in vitro* tests should be initiated. Appropriate endpoint specific *in vivo* higher tier testing should be conducted in response to equivocal / positive *in vitro* results. At the present time other non-animal methods to follow up an *in vitro* positive genotoxicity results are not accepted within the Regulation (EC)1107/2009 framework.

For the assessment of the three fludioxonil dietary metabolites used in this case study, additional (Q)SAR alert(s) to parent and other groups within the dietary metabolite assessment were identified. Initial attempts to mitigate these alerts through grouping, read-across and expert review, including consideration of the predictivity of the additional alerts for SYN518580 and CGA335892, were made to the regulatory agencies. For example, in the case of SYN518580, where the structural alert “SA_34” (H-acceptor-path3-H-acceptor) for *in vivo* micronucleus formation was triggered in ToxTree and the OECD QSAR Toolbox, the promiscuity and low predictivity of the alert was communicated to the EU regulators and a read-across to a ground water metabolite with a negative genotoxicity data package was proposed. For CGA335892, a proposal was made to read across to available parent (fludioxonil) data. The authors did not have a genotoxicity concern for SYN518580 and CGA335892 based on expert review of the *in silico* data. Ultimately these proposals were considered by the EU regulators not sufficient mitigation to address the additional (Q)SAR alerts, therefore new *in vitro* genotoxicity studies on these two metabolites (CGA335892 and SYN518580) and CGA227731 were conducted (bacterial reverse mutation test, *in vitro* micronucleus).

An overview of the studies conducted on fludioxonil and the metabolites presented in this case study are presented in Table 5. The genotoxicity evaluation for CGA335892 and SYN518580 showed high similarity; the bacterial reverse mutation tests were negative, the MNvit assays positive and followed up with a negative *in vivo* micronucleus assay in each case.

**Table 5:** Overview of genotoxicity studies conducted on fludioxonil and its metabolites CGA335892, SYN518580 and CGA227731.

| Compound | *In vitro* |  |  |  | *In vivo* |  |  |  |  |
| --- | --- | --- | --- | --- | --- | --- | --- | --- | --- |
|  | Bacterial reverse mutation | MNvit | IVC | HPRT | MN # | Comet ## | TGR### | UDS | Rodent dominant lethal |
| Fludioxonil*** | -ve |  | +ve | -ve | -ve (BM); -ve (hepatocytes*) |  |  | -ve | -ve |
| CGA335892 | -ve | +ve |  |  | -ve (PB) |  |  |  |  |
| SYN518580 | -ve | +ve |  |  | -ve (PB) |  |  |  |  |
| CGA227731 | +ve (TA1537)*** | -ve*** |  |  |  | -ve *** (liver** and duodenum) | -ve (liver, duodenum, bone marrow) |  |  |

In the bacterial reverse mutation tests on CGA335892 and SYN518580, extreme cytotoxicity and /or pseudorevertants were observed which limited the concentration level tested. Pseudorevertants may be caused when the test item is so cytotoxic that most of the bacterial cells are killed. The few surviving cells then have access to sufficient histidine/tryptophan to allow them to grow to macroscopic size and appear to be revertants. They are easily distinguishable from true revertants as there is no background lawn present and the phenomenon was described in one of the original papers from the Ames laboratory though they did not use the term pseudorevertants (Ames *et al.,* 1975).

Since the pseudorevertants are caused by excessive cytotoxicity, not mutagenicity, they have no relevance on the mutagenic evaluation of the test substance other than to limit the concentration that can be evaluated. Both compounds were tested up to a cytotoxic concentration in each of the experimental conditions with at least 4 non-cytotoxic and 5 analysable concentrations scored for each strain in each condition. Therefore, the tests fulfilled the requirements of the OECD 471 (2020) guideline. As there were no relevant reproducible increases in revertants in any condition, SYN518580 are CGA335892 were concluded to be negative (i.e. non-mutagenic) in the bacterial reverse mutation test.

The MNvit for CGA335892 and SYN518580 were both clearly positive requiring an *in vivo* micronucleus assay be conducted for each compound. The *in vivo* erythrocyte micronucleus assay is the most appropriate direct follow up to the *in vitro* micronucleus assay as both assays cover the same genetic toxicity endpoints (clastogenicity and aneugenicity). The *in vivo* MN studies on CGA335892 and SYN518580 were negative. Each compound was quantified in the plasma of exposed mice in their respective range finder assays at all timepoints evaluated (1, 4, 24 h post second dose) using validated analytical methods. The quantification of the test substance in the plasma represents a line of evidence that the bone marrow was exposed to the test substance in the main micronucleus test (EFSA 2017). The clinical signs observed on the SYN518580 main MN study; (piloerection, decreased activity, partially closed eyes) and CGA335892 range finder (prostration, low posture, partially closed eyes, intermittent laboured breathing) are further lines of evidence of systemic exposure. Bone marrow is a highly perfused tissue and quantification of the test substance in the plasma also demonstrates the exposure of the bone marrow. SYN518580 was tested up to the maximum regulatory dose of 2000 mg/kg/day (OECD 474, 2016). CGA335892 was tested up to the MTD of 800 mg/kg/day. The dose level scale used to establish the MTD was derived from Mackay and Elliot (1992); 1250 mg/kg/day was not tolerated in the range finder and the next dose level down from this was 800 mg/kg/day. Whilst there were no clinical observations at the top dose level tested in the main study, for animal welfare reasons and to align with guideline study requirements it is not possible or necessary to test to a higher level.

EFSA (2011) guidance on genotoxicity testing indicates that a compound that has tested positive in an *in vitro* genotoxicity test but negative in an appropriate *in vivo* follow-up test, addressing the relevant endpoint(s), is concluded to have no genotoxic potential as long as there is significant evidence of target tissue exposure. For CGA335892 and SYN518580, as the bacterial reverse mutation assays and *in vivo* micronucleus assays, addressing between them all genotoxicity endpoints, were negative and significant evidence of bone marrow exposure is available, these dietary metabolites are concluded to be of no genotoxic concern.

The *in vitro* genotoxicity data for CGA227731 showed differential results compared to CGA335892, SYN518580 and all other fludioxonil dietary metabolites. The bacterial reverse mutation test was positive in strain TA1537 with reproducible concentration-related increases in both the presence and absence of metabolic activation. This result was concordant with the CAESAR model (Q)SAR prediction for mutagenicity for this compound. In contrast, the *in vitro* micronucleus assay was clearly negative confirming that CGA227731 is neither clastogenic nor aneugenic *in vitro*.

At the time of the positive bacterial reverse mutation test on CGA227731, two options were proposed for the follow up a positive *in vitro* gene mutation assay; an *in vivo* alkaline Comet assay (OECD 489) or TGR (OECD 488) (EFSA 2016). With the recent adoption of the Mammalian Erythrocyte Pig-a Gene Mutation Assay guideline (OECD 470 2022, 2025) another *in vivo* test is now potentially available, though the use of this test may be limited as it only allows assessment for mutations in the bone marrow and not at a site of contact or liver.

The *in vivo* alkaline Comet assay in the duodenum and liver was initially conducted on CGA227731 to follow up the positive response in the bacterial reverse mutation assay. The *in vivo* alkaline Comet assay (Comet) (OECD 489) is a readily accessible tool to investigate DNA damage *in vivo* and has been proposed as a higher tier assay to follow-up positive *in vitro* findings, (EFSA 2011, EU 2013, EFSA 2016). Comparisons have also been made as to the suitability of the Comet or TGR assays as *in vivo* follow-up assays (Kirkland *et al.,* 2019; Robison *et al.,* 2021; Zeller *et al.,* 2019, Zeller *et al.,* 2020).

The Comet assay on CGA227731 in the duodenum was clearly negative however, a small statistically significant increase in %TI was observed at the top dose in the liver. The %TI remained within the observed range of the mHCD and at the top dose there was less than a 1% absolute change in tail intensity. There was also a dose dependent increase in the number of hedgehogs. Hedgehog cells are cells that have a small or non-existent head and a large diffuse tail and are scored independently of the genotoxicity endpoint (%TI) for the Comet assay (OECD 489, 2016). An increase in the mean number of hedgehog cells was observed in the livers of animals receiving 1000 mg/kg/ day versus the concurrent control (2.8/150 cells versus 0 in concurrent control) however, there were no increases in median %TI at this dose level. The increase in hedgehogs in the absence of an increase in %TI at this dose level is suggestive of cytotoxicity in the liver rather than evidence of genotoxicity. This is supported by the increase in mitotic figures seen in the liver of two animals at 2000 mg/kg/day. It was therefore concluded that the minimal increase that remained within the historical control range was unlikely to be of genotoxic concern. CGA227731 was concluded to be negative in the liver and duodenum in the rat Comet assay. It should also be noted that the mutagenicity in the bacterial reverse mutation assay was observed in both the presence and absence of metabolic activation, if the increase in %TI in the liver of the Comet was genuinely reflective of the mutagenic potential of CGA227731, then a response in the duodenum would also be expected.

The *in vivo* Comet assay on CGA227731 has been reviewed by both EFSA and the Joint FAO/WHO Meeting on Pesticide Residues (JMPR). The JMPR evaluation concluded that the *in vivo* Comet assay was negative but that the Comet assay was not a suitable assay for this compound. CGA227731 is essentially a planar molecule and they proposed that the response in TA1537 was most likely due to frameshift mutations caused by a planar DNA intercalating agent and that this type of mutation is out of the applicability domain of the *in vivo* Comet assay (JMPR 2022). It may be noted that intercalation has been shown to induce a response in the (*in vitro*) Comet assay previously (Henderson *et al.,* 1998). EFSA Peer Review Meeting concluded that CGA227731 was genotoxic both *in vitro* and *in vivo* based on the response in TA1537 and the Comet assay (EFSA, 2019).

Whilst the Comet assay is versatile it does not however address the same apical endpoint of concern for the follow-up of an *in vitro* mutagenicity finding, namely gene mutation, but rather primary DNA damage and is noted as an “indicator assay” (EFSA 2011). As such the Comet assay is vulnerable to a number of confounding factors. For example, there is currently no consensus opinion on the cause or relevance of “hedgehogs” (Lorenzo *et al.,* 2013; Guerard *et al.,* 2014) or the critical aspects for the conduct of the Comet assay to ensure data reliability (Vasquez and Dewhurst, 2024). In the situation where a Comet assay has afforded a result open to differential interpretation (as in the case of CGA227731), then it is often appropriate to conduct a further *in vivo* assay such as the TGR addressing the apical mutagenicity endpoint. Given the differential interpretation at the EFSA Peer review meeting, a TGR assay in the BigBlue® rat was initiated to investigate the *in vivo* mutagenic potential of CGA227731. The BigBlue® rat was selected to ensure consistency in the species between the TGR and Comet assay. The same dosing route and vehicle was also used. Liver, bone marrow and duodenum were selected for the analysis of mutant frequency to ensure a slowly proliferating tissue (liver), and a rapidly dividing tissue (bone marrow/duodenum) was exposed to CGA227731.

CGA227731 did not induce cII mutations in any of the tissues evaluated and is therefore concluded to be not mutagenic *in vivo*. This study has been reviewed by the FAO/WHO and they agreed with this conclusion (JMPR 2022).

The data presented on the three fludioxonil dietary metabolites in this case study illustrate the detailed and comprehensive data generation undertaken to address the EU regulatory data requirements. During this process a work-flow encompassing *in silico*, *in vitro* and *in vivo* data generation is progressed to arrive at a robust assessment of genotoxicity. Following this process and based on an extensive battery of *in vitro* and *in vivo* genotoxicity test data the dietary metabolites of fludioxonil, CGA335892, SYN518580 and CGA227731, are concluded to not be of genotoxic concern under the Regulation (EC)1107/2009 framework. Since all other identified dietary metabolites of fludioxonil have been concluded to be non-genotoxic during the AIR3 renewal (ANSES 2023) based on this tiered assessment process, it is further concluded that all identified dietary metabolites of fludioxonil are non-genotoxic.

## Abbreviations

CBPI: cytokinesis block proliferation index
EC: European Commission
EFSA: European Food Safety Authority
HCD: historical control data
MN: micronuclei
MNvit: *in vitro* mammalian cell micronucleus test
MN-BN: micronucleated binucleate cells
MN-RET: micronucleated reticulocytes
MTD: maximum tolerated dose
OECD: Organisation for Economic Co-operation and Development
(Q)SAR: (Quantitative) Structure Activity Relationships
RET: reticulocytes
TGR: transgenic rodent gene mutation assay

## Declaration of generative AI and AI-assisted technologies in the writing process

During the preparation of this work the author(s) used SynGPT [version 3.5] (2024), in order to help edit parts of the method section. After using this tool, the authors reviewed and further edited the content as needed and take full responsibility for the content of the published article.

## ARRIVE statement

All animal studies are reported according to the ARRIVE guidelines.

## Funding

The studies were conducted at 3^rd^ party contract research organisations and funded by Syngenta Ltd.

## Declaration of competing interests

The authors are employees of Syngenta Ltd. The authors have no other competing interests.

## CRediT authorship contribution statement

**Zofi McKenzie:** conceptualization, funding acquisition, project administration, supervision, visualization, writing-original draft, writing – review and editing. **Anne-Sophie Parant:** project administration, writing – original draft, writing – review and editing. **Katy L. Bridgwood:** writing - original draft, writing – review and editing. **Ewan D. Booth:** conceptualization, writing - original draft, writing – review and editing

## Supplementary Information

**Supplementary Table 1:**
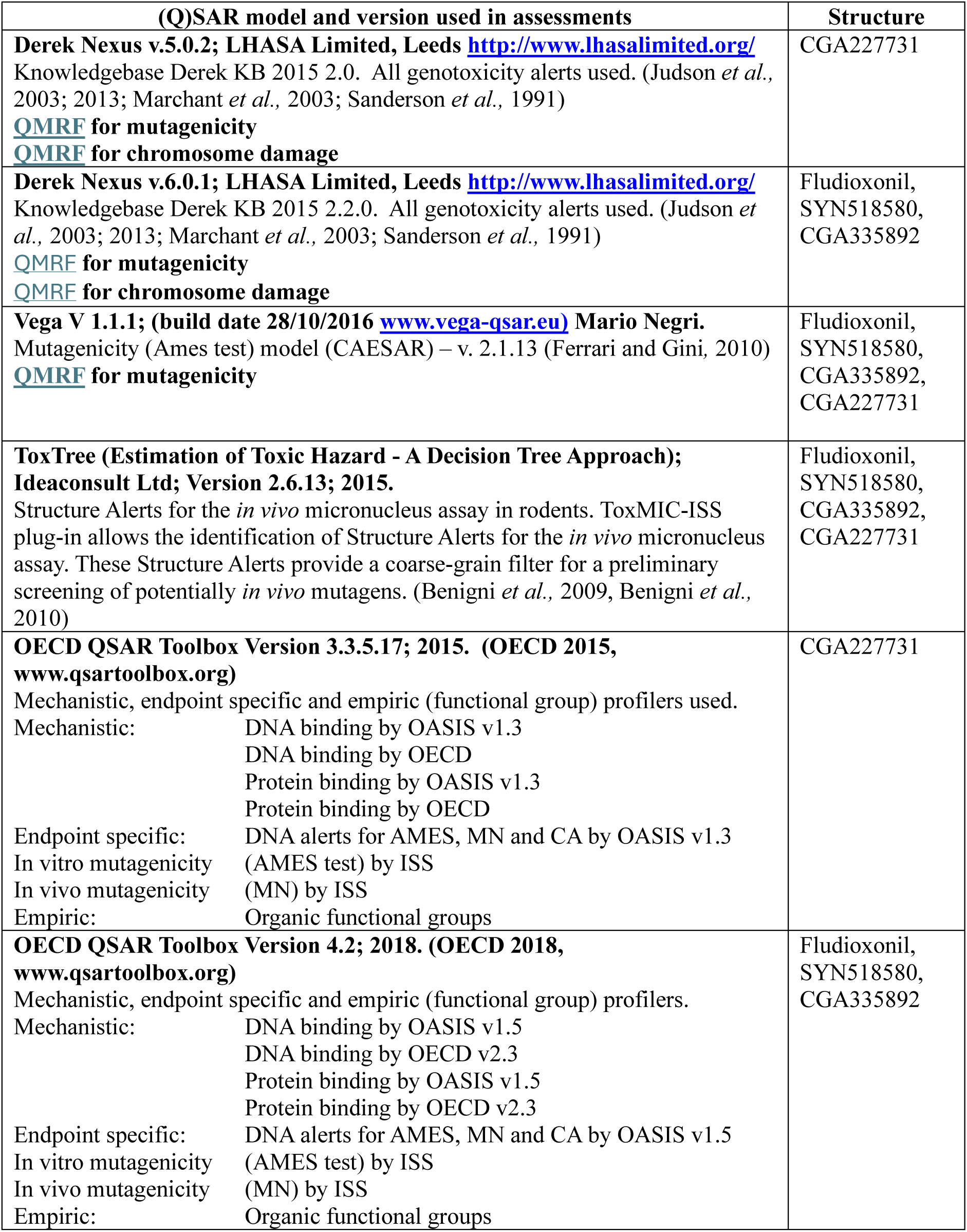
(Q)SAR Software.

| (Q)SAR model and version used in assessments | Structure |
| --- | --- |
| **Derek Nexus v.5.0.2; LHASA Limited, Leeds** http://www.lhasalimited.org/<br>Knowledgebase Derek KB 2015 2.0. All genotoxicity alerts used. (Judson *et al.*, 2003; 2013; Marchant *et al.*, 2003; Sanderson *et al.*, 1991)<br>QMRF for mutagenicity<br>QMRF for chromosome damage | CGA227731 |
| **Derek Nexus v.6.0.1; LHASA Limited, Leeds** http://www.lhasalimited.org/<br>Knowledgebase Derek KB 2015 2.2.0. All genotoxicity alerts used. (Judson *et al.*, 2003; 2013; Marchant *et al.*, 2003; Sanderson *et al.*, 1991)<br>QMRF for mutagenicity<br>QMRF for chromosome damage | Fludioxonil,<br>SYN518580,<br>CGA335892 |
| **Vega V 1.1.1; (build date 28/10/2016** www.vega-qsar.eu) **Mario Negri.**<br>Mutagenicity (Ames test) model (CAESAR) – v. 2.1.13 (Ferrari and Gini, 2010)<br>QMRF for mutagenicity | Fludioxonil,<br>SYN518580,<br>CGA335892,<br>CGA227731 |
| **ToxTree (Estimation of Toxic Hazard - A Decision Tree Approach);**<br>**Ideaconsult Ltd; Version 2.6.13; 2015.**<br>Structure Alerts for the *in vivo* micronucleus assay in rodents. ToxMIC-ISS plug-in allows the identification of Structure Alerts for the *in vivo* micronucleus assay. These Structure Alerts provide a coarse-grain filter for a preliminary screening of potentially *in vivo* mutagens. (Benigni *et al.*, 2009, Benigni *et al.*, 2010) | Fludioxonil,<br>SYN518580,<br>CGA335892,<br>CGA227731 |
| **OECD QSAR Toolbox Version 3.3.5.17; 2015. (OECD 2015, www.qsartoolbox.org)**<br>Mechanistic, endpoint specific and empiric (functional group) profilers used.<br>Mechanistic: DNA binding by OASIS v1.3<br>DNA binding by OECD<br>Protein binding by OASIS v1.3<br>Protein binding by OECD<br>Endpoint specific: DNA alerts for AMES, MN and CA by OASIS v1.3<br>In vitro mutagenicity (AMES test) by ISS<br>In vivo mutagenicity (MN) by ISS<br>Empiric: Organic functional groups | CGA227731 |
| **OECD QSAR Toolbox Version 4.2; 2018. (OECD 2018, www.qsartoolbox.org)**<br>Mechanistic, endpoint specific and empiric (functional group) profilers.<br>Mechanistic: DNA binding by OASIS v1.5<br>DNA binding by OECD v2.3<br>Protein binding by OASIS v1.5<br>Protein binding by OECD v2.3<br>Endpoint specific: DNA alerts for AMES, MN and CA by OASIS v1.5<br>In vitro mutagenicity (AMES test) by ISS<br>In vivo mutagenicity (MN) by ISS<br>Empiric: Organic functional groups | Fludioxonil,<br>SYN518580,<br>CGA335892 |

**Supplementary Table 2:** Plate Incorporation Experiment Results with CGA335892 in the Presence of Metabolic Activation.

| Concentration<br>per plate |  | Number of revertant colonies per plate |  |  |  |  |  |  |  |  |  |  |  |
| --- | --- | --- | --- | --- | --- | --- | --- | --- | --- | --- | --- | --- | --- |
|  |  | *S. typhimurium* LT2 |  |  |  |  |  |  |  | *E. coli* WP2 |  |  |  |
| μg | μmol | TA1535 |  | TA1537 |  | TA98 |  | TA100 |  | *uvrA* /pKM101 |  | pKM101 |  |
| Solvent control |  | 11 | 10 | 13 | 12 | 33 | 35 | 139 | 143 | 201 | 202 | 100 | 104 |
|  |  | 10 | 0.6 | 14 | 2.1 | 36 | 1.5 | 150 | 5.9 | 198 | 5.1 | 112 | 7.2 |
|  |  | 10 |  | 10 |  | 35 |  | 141 |  | 208 |  | 99 |  |
| 1.6 | 0.006 | 10 | 10 | 14 | 11 | 27 | 31 | 134 | 120 | 211 | 217 | 116 | 105 |
|  |  | 10 | 0.0 | 9 | 2.5 | 38 | 5.9 | 112 | 12.4 | 221 | 5.3 | 97 | 9.8 |
|  |  | 10 | 1.0 | 11 | 0.9 | 29 | 0.9 | 113 | 0.8 | 219 | 1.1 | 102 | 1.0 |
| 5 | 0.019 | 7 | 10 | 12 | 11 | 36 | 32 | 109 | 110 | 202 | 210 | 105 | 106 |
|  |  | 10 | 2.5 | 10 | 1.0 | 29 | 3.6 | 121 | 10.5 | 213 | 7.4 | 98 | 8.5 |
|  |  | 12 | 1.0 | 11 | 0.9 | 31 | 0.9 | 100 | 0.8 | 216 | 1.0 | 115 | 1.0 |
| 16 | 0.061 | 11 | 9 | 9 | 11 | 33 | 33 | 129 | 130 | 220 | 200 | 116 | 106 |
|  |  | 8 | 1.5 | 11 | 1.5 | 32 | 1.5 | 122 | 9.1 | 196 | 18.3 | 94 | 11.1 |
|  |  | 9 | 0.9 | 12 | 0.9 | 35 | 0.9 | 140 | 0.9 | 184 | 1.0 | 107 | 1.0 |
| 50 | 0.189 | 12 | 12 | 13 | 11 | 29 | 33 | 136 | 131 | 209 | 207 | 100 | 98 |
|  |  | 14 | 2.0 | 10 | 2.1 | 40 | 5.9 | 128 | 4.4 | 207 | 2.5 | 103 | 6.2 |
|  |  | 10 | 1.2 | 9 | 0.9 | 31 | 0.9 | 129 | 0.9 | 204 | 1.0 | 91 | 0.9 |
| 160 | 0.606 | 10 | 10 | 12 | 11 | 37 | 37 | 121 | 110 | 203 | 200 | 110 | 108 |
|  |  | 8 | 2.0 | 10 | 1.2 | 41 | 4.0 | 104 | 9.3 | 201 | 3.1 | 103 | 4.7 |
|  |  | 12 | 1.0 | 10 | 0.9 | 33 | 1.1 | 106 | 0.8 | 197 | 1.0 | 112 | 1.0 |
| 500 | 1.89 | 0 f | 0 | 0 f | 0 | 0 f | 0 | 0 f | 0 | 133 v | 115 | 17 v | 25 |
|  |  | 0 f | 0.0 | 0 f | 0.0 | 0 f | 0.0 | 0 f | 0.0 | 97 v | 18.0 | 30 v | 7.2 |
|  |  | 0 f | 0.0 | 0 f | 0.0 | 0 f | 0.0 | 0 f | 0.0 | 116 v | 0.6 | 29 v | 0.2 |
| 1600 | 6.06 | 0 n | 0 | 0 n | 0 | 0 n | 0 | 0 n | 0 | 0 f | 0 | 0 f | 0 |
|  |  | 0 n | 0.0 | 0 n | 0.0 | 0 n | 0.0 | 0 n | 0.0 | 0 f | 0.0 | 0 f | 0.0 |
|  |  | 0 n | 0.0 | 0 n | 0.0 | 0 n | 0.0 | 0 n | 0.0 | 0 f | 0.0 | 0 f | 0.0 |
| 5000 | 18.9 | 0 np | 0 | 0 np | 0 | 0 np | 0 | 0 np | 0 | 0 np | 0 | 0 np | 0 |
|  |  | 0 np | 0.0 | 0 np | 0.0 | 0 np | 0.0 | 0 np | 0.0 | 0 np | 0.0 | 0 np | 0.0 |
|  |  | 0 np | 0.0 | 0 np | 0.0 | 0 np | 0.0 | 0 np | 0.0 | 0 np | 0.0 | 0 np | 0.0 |
| Positive control |  | 271 | 242 | 195 | 213 | 1785 | 1792 | 2252 | 2012 | 2374 | 2487 | 282 | 268 |
|  |  | 259 | 40.3 | 217 | 16.4 | 1795 | 6.4 | 1869 | 209.3 | 2657 | 149.9 | 241 | 23.1 |
|  |  | 196 | 24.2 | 227 | 17.8 | 1797 | 51.2 | 1914 | 14.1 | 2430 | 12.3 | 280 | 2.6 |

Positive controls
TA1535 2-aminoanthracene 2 μg/plate TA1537 2-aminoanthracene 2 μg/plate TA98 2-aminoanthracene 2 μg/plate TA100 2-aminoanthracene 2 μg/plate *uvrA*/pKM101 2-aminoanthracene 20 μg/plate pKM101 2-aminoanthracene 20 μg/plate
Comments
v: very reduced lawn f: pseudorevertants n: no survival p: precipitate
= mean *italics* = Standard Deviation **bold** = fold increase over solvent control
TA1535 and TA1537 solvent control and test item plates were scored manually, all other plates were scored using Sorcerer image analysis system.

**Supplementary Table 3:** Plate Incorporation Experiment Results with CGA335892 in the Absence of Metabolic Activation.

| Concentration<br>per plate |  | Number of revertant colonies per plate |  |  |  |  |  |  |  |  |  |  |  |
| --- | --- | --- | --- | --- | --- | --- | --- | --- | --- | --- | --- | --- | --- |
|  |  | *S. typhimurium* LT2 |  |  |  |  |  |  |  | *E. coli* WP2 |  |  |  |
| μg | μmol | TA1535 |  | TA1537 |  | TA98 |  | TA100 |  | *uvrA* /pKM101 |  | pKM101 |  |
| Solvent control |  | 13 | 13 | 11 | 11 | 26 | 27 | 127 | 121 | 151 | 150 | 72 | 71 |
|  |  | 12 | 1.0 | 10 | 1.0 | 27 | 0.6 | 116 | 5.7 | 142 | 7.5 | 79 | 8.0 |
|  |  | 14 |  | 12 |  | 27 |  | 119 |  | 157 |  | 63 |  |
| 1.6 | 0.006 | 12 | 12 | 9 | 10 | 33 | 32 | 117 | 119 | 166 | 153 | 64 | 67 |
|  |  | 11 | 1.0 | 12 | 1.5 | 29 | 3.1 | 119 | 2.0 | 152 | 13.0 | 73 | 5.5 |
|  |  | 13 | 0.9 | 10 | 0.9 | 35 | 1.2 | 121 | 1.0 | 140 | 1.0 | 63 | 0.9 |
| 5 | 0.019 | 12 | 12 | 9 | 12 | 31 | 27 | 116 | 117 | 154 | 148 | 70 | 73 |
|  |  | 13 | 1.5 | 12 | 3.0 | 26 | 3.2 | 113 | 5.1 | 148 | 6.5 | 83 | 8.9 |
|  |  | 10 | 0.9 | 15 | 1.1 | 25 | 1.0 | 123 | 1.0 | 141 | 1.0 | 66 | 1.0 |
| 16 | 0.061 | 10 | 11 | 10 | 10 | 24 | 26 | 109 | 111 | 140 | 139 | 59 | 73 |
|  |  | 10 | 2.3 | 9 | 0.6 | 23 | 4.9 | 125 | 13.6 | 141 | 2.1 | 80 | 11.8 |
|  |  | 14 | 0.8 | 10 | 0.9 | 32 | 1.0 | 98 | 0.9 | 137 | 0.9 | 79 | 1.0 |
| 50 | 0.189 | 14 | 11 | 11 | 9 | 16 | 19 | 97 | 103 | 146 | 148 | 53 | 65 |
|  |  | 10 | 2.6 | 8 | 1.7 | 24 | 4.2 | 116 | 11.3 | 147 | 3.2 | 80 | 13.7 |
|  |  | 9 | 0.8 | 8 | 0.8 | 18 | 0.7 | 96 | 0.9 | 152 | 1.0 | 62 | 0.9 |
| 160 | 0.606 | 8 | 7 | 8 | 7 | 29 | 28 | 91 | 102 | 140 | 138 | 64 | 67 |
|  |  | 6 | 1.0 | 7 | 0.6 | 21 | 7.0 | 109 | 9.6 | 137 | 1.5 | 67 | 2.5 |
|  |  | 7 | 0.5 | 7 | 0.6 | 35 | 1.0 | 106 | 0.8 | 138 | 0.9 | 69 | 0.9 |
| 500 | 1.89 | 0 f | 0 | 0 f | 0 | 0 f | 0 | 0 f | 0 | 121 v | 95 | 31 v | 30 |
|  |  | 0 f | 0.0 | 0 f | 0.0 | 0 f | 0.0 | 0 f | 0.0 | 104 v | 31.5 | 35 v | 5.0 |
|  |  | 0 f | 0.0 | 0 f | 0.0 | 0 f | 0.0 | 0 f | 0.0 | 60 v | 0.6 | 25 v | 0.4 |
| 1600 | 6.06 | 0 n | 0 | 0 n | 0 | 0 n | 0 | 0 n | 0 | 0 f | 0 | 0 f | 0 |
|  |  | 0 n | 0.0 | 0 n | 0.0 | 0 n | 0.0 | 0 n | 0.0 | 0 f | 0.0 | 0 f | 0.0 |
|  |  | 0 n | 0.0 | 0 n | 0.0 | 0 n | 0.0 | 0 n | 0.0 | 0 f | 0.0 | 0 f | 0.0 |
| 5000 | 18.9 | 0 np | 0 | 0 np | 0 | 0 np | 0 | 0 np | 0 | 0 np | 0 | 0 np | 0 |
|  |  | 0 np | 0.0 | 0 np | 0.0 | 0 np | 0.0 | 0 np | 0.0 | 0 np | 0.0 | 0 np | 0.0 |
|  |  | 0 np | 0.0 | 0 np | 0.0 | 0 np | 0.0 | 0 np | 0.0 | 0 np | 0.0 | 0 np | 0.0 |
| Positive control |  | 524 | 467 | 296 | 257 | 307 | 353 | 584 | 528 | 1077 | 1069 | 706 | 712 |
|  |  | 410 | 57.0 | 209 | 44.2 | 366 | 41.1 | 499 | 48.2 | 1058 | 9.8 | 642 | 73.2 |
|  |  | 468 | 35.9 | 266 | 23.4 | 386 | 13.1 | 502 | 4.4 | 1072 | 7.1 | 788 | 10.0 |

Positive controls
TA1535 sodium azide 0.5 μg/plate TA1537 9-aminoacridine.HCl 50 μg/plate TA98 2-nitrofluorene 1 μg/plate TA100 sodium azide 0.5 μg/plate *uvrA*/pKM101 potassium dichromate 25 μg/plate pKM101 potassium dichromate 25 μg/plate
Comments
v: very reduced lawn f: pseudorevertants n: no survival p: precipitate
= mean *italics* = Standard Deviation **bold** = fold increase over solvent control
TA1535 and TA1537 solvent control and test item plates were scored manually, all other plates were scored using Sorcerer image analysis system.

**Supplementary Table 4:** CGA335892: Liquid Pre-Incubation Experiment Results with CGA335892 in the Presence of Metabolic Activation.

| Concentration<br>per plate |  | Number of revertant colonies per plate |  |  |  |  |  |  |  |  |  |  |  |
| --- | --- | --- | --- | --- | --- | --- | --- | --- | --- | --- | --- | --- | --- |
|  |  | *S. typhimurium* LT2 |  |  |  |  |  |  |  | *E. coli* WP2 |  |  |  |
|  |  | TA1535 |  | TA1537 |  | TA98 |  | TA100 |  | *uvrA* /pKM101 |  | pKM101 |  |
| μg | μmol |  |  |  |  |  |  |  |  |  |  |  |  |
| Solvent control |  | 13 | 12 | 11 | 11 | 36 | 35 | 137 | 130 | 167 | 176 | 91 | 95 |
|  |  | 11 | 1.0 | 12 | 1.0 | 36 | 2.3 | 130 | 6.5 | 163 | 19.7 | 103 | 7.2 |
|  |  | 12 |  | 10 |  | 32 |  | 124 |  | 199 |  | 90 |  |
| 0.5 | 0.00189 | 12 | 12 | 11 | 11 | 40 | 39 | 142 | 142 | 159 | 177 | 96 | 92 |
|  |  | 13 | 1.5 | 12 | 1.0 | 38 | 1.2 | 128 | 13.5 | 176 | 18.5 | 96 | 6.4 |
|  |  | 10 | 1.0 | 10 | 1.0 | 38 | 1.1 | 155 | 1.1 | 196 | 1.0 | 85 | 1.0 |
| 1.6 | 0.00606 | 10 | 10 | 10 | 10 | 29 | 30 | 149 | 151 | 181 | 169 | 93 | 87 |
|  |  | 9 | 0.6 | 9 | 1.5 | 31 | 1.2 | 150 | 3.2 | 159 | 11.2 | 73 | 11.8 |
|  |  | 10 | 0.8 | 12 | 0.9 | 29 | 0.9 | 155 | 1.2 | 166 | 1.0 | 94 | 0.9 |
| 5 | 0.0189 | 12 | 12 | 11 | 11 | 27 | 28 | 118 | 126 | 159 | 163 | 77 | 80 |
|  |  | 13 | 1.0 | 10 | 1.0 | 29 | 1.2 | 127 | 7.5 | 173 | 8.7 | 78 | 3.8 |
|  |  | 11 | 1.0 | 12 | 1.0 | 29 | 0.8 | 133 | 1.0 | 157 | 0.9 | 84 | 0.8 |
| 16 | 0.0606 | 10 | 11 | 10 | 11 | 35 | 32 | 124 | 126 | 168 | 178 | 92 | 87 |
|  |  | 11 | 1.0 | 11 | 1.0 | 30 | 2.5 | 122 | 5.9 | 178 | 10.0 | 93 | 9.5 |
|  |  | 12 | 0.9 | 12 | 1.0 | 32 | 0.9 | 133 | 1.0 | 188 | 1.0 | 76 | 0.9 |
| 50 | 0.189 | 10 | 11 | 12 | 12 | 40 | 36 | 122 | 138 | 176 | 183 | 86 | 90 |
|  |  | 10 | 1.2 | 12 | 0.0 | 36 | 4.5 | 142 | 14.8 | 195 | 10.2 | 91 | 3.2 |
|  |  | 12 | 0.9 | 12 | 1.1 | 31 | 1.0 | 151 | 1.1 | 179 | 1.0 | 92 | 0.9 |
| 160 | 0.606 | 4 r | 3 | 4 r | 5 | 27 r | 21 | 78 r | 107 | 109 s | 134 | 60 s | 86 |
|  |  | 3 r | 1.0 | 5 r | 1.0 | 19 r | 5.3 | 116 r | 26.1 | 153 s | 22.5 | 98 s | 22.2 |
|  |  | 2 r | 0.3 | 6 r | 0.5 | 17 r | 0.6 | 128 r | 0.8 | 139 s | 0.8 | 99 s | 0.9 |
| 500 | 1.89 | 0 f | 0 | 0 f | 0 | 0 f | 0 | 0 f | 0 | 56 v | 44 | 34 v | 41 |
|  |  | 0 f | 0.0 | 0 f | 0.0 | 0 f | 0.0 | 0 f | 0.0 | 11 v | 28.9 | 47 v | 6.6 |
|  |  | 0 f | 0.0 | 0 f | 0.0 | 0 f | 0.0 | 0 f | 0.0 | 65 v | 0.3 | 42 v | 0.4 |
| 1600 | 6.06 | 0 np | 0 | 0 np | 0 | 0 np | 0 | 0 np | 0 | 0 np | 0 | 0 np | 0 |
|  |  | 0 np | 0.0 | 0 np | 0.0 | 0 np | 0.0 | 0 np | 0.0 | 0 np | 0.0 | 0 np | 0.0 |
|  |  | 0 np | 0.0 | 0 np | 0.0 | 0 np | 0.0 | 0 np | 0.0 | 0 np | 0.0 | 0 np | 0.0 |
| Positive control |  | 137 | 155 | 193 | 174 | 2080 | 2030 | 2303 | 2283 | 2049 | 1794 | 288 | 274 |
|  |  | 149 | 21.1 | 194 | 34.4 | 1998 | 43.7 | 2295 | 28.6 | 1581 | 236.8 | 320 | 54.4 |
|  |  | 178 | 12.9 | 134 | 15.8 | 2013 | 58.0 | 2250 | 17.6 | 1752 | 10.2 | 214 | 2.9 |

Positive controls
TA1535 2-aminoanthracene 2 μg/plate TA1537 2-aminoanthracene 2 μg/plate TA98 2-aminoanthracene 2 μg/plate TA100 2-aminoanthracene 2 μg/plate *uvrA*/pKM101 2-aminoanthracene 20 μg/plate pKM101 2-aminoanthracene 20 μg/plate
Comments
s: slightly reduced lawn r: reduced lawn v: very reduced lawn f: pseudorevertants n: no survival p: precipitate
= mean *italics* = Standard Deviation **bold** = fold increase over solvent control
TA1535 and TA1537 solvent control and test item plates were scored manually, all other plates were scored using Sorcerer image analysis system.

**Supplementary Table 5:**
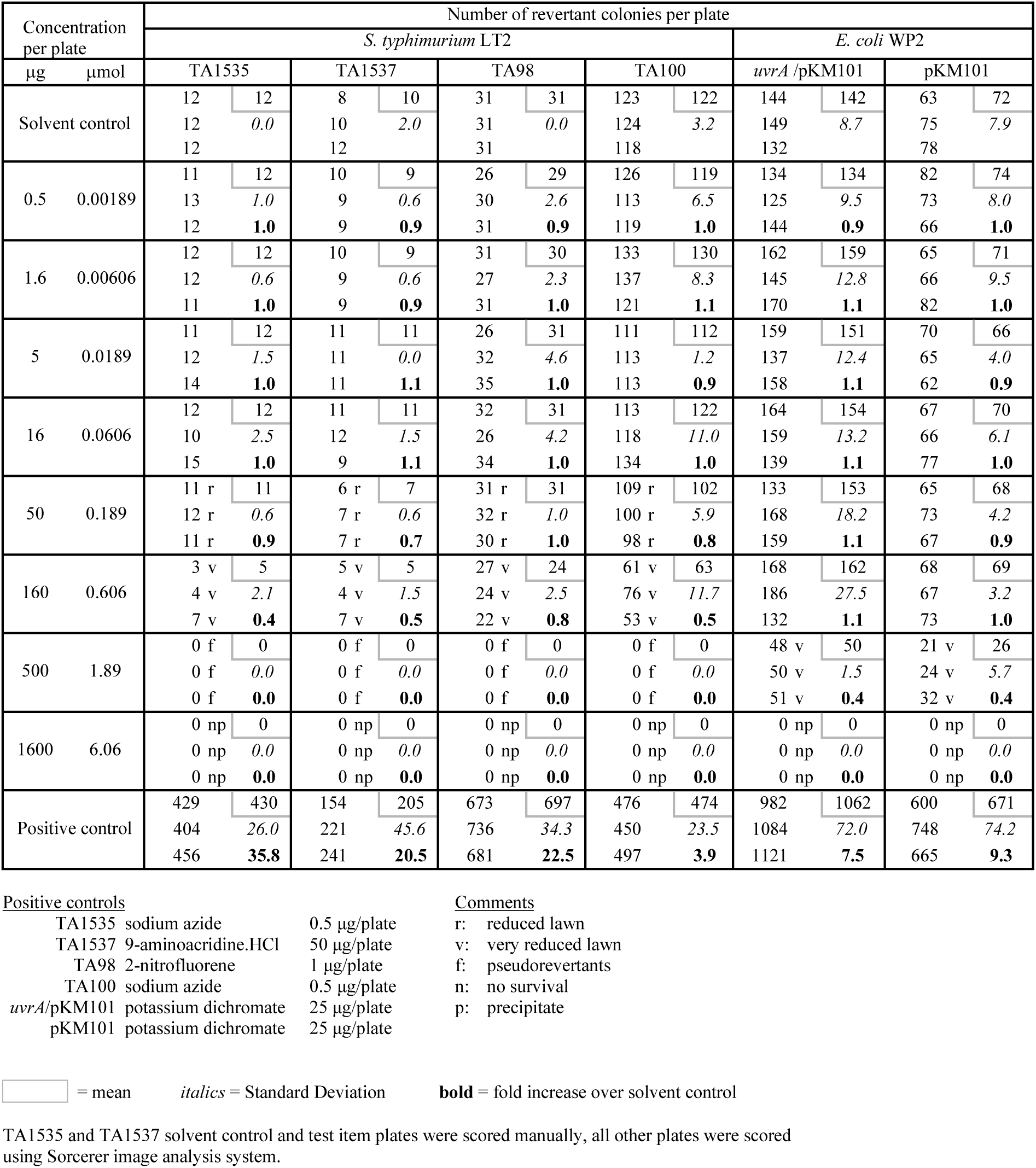
Liquid Pre-Incubation Experiment Results with CGA335892 in the Absence Metabolic Activation.

**Supplementary Table 6:**
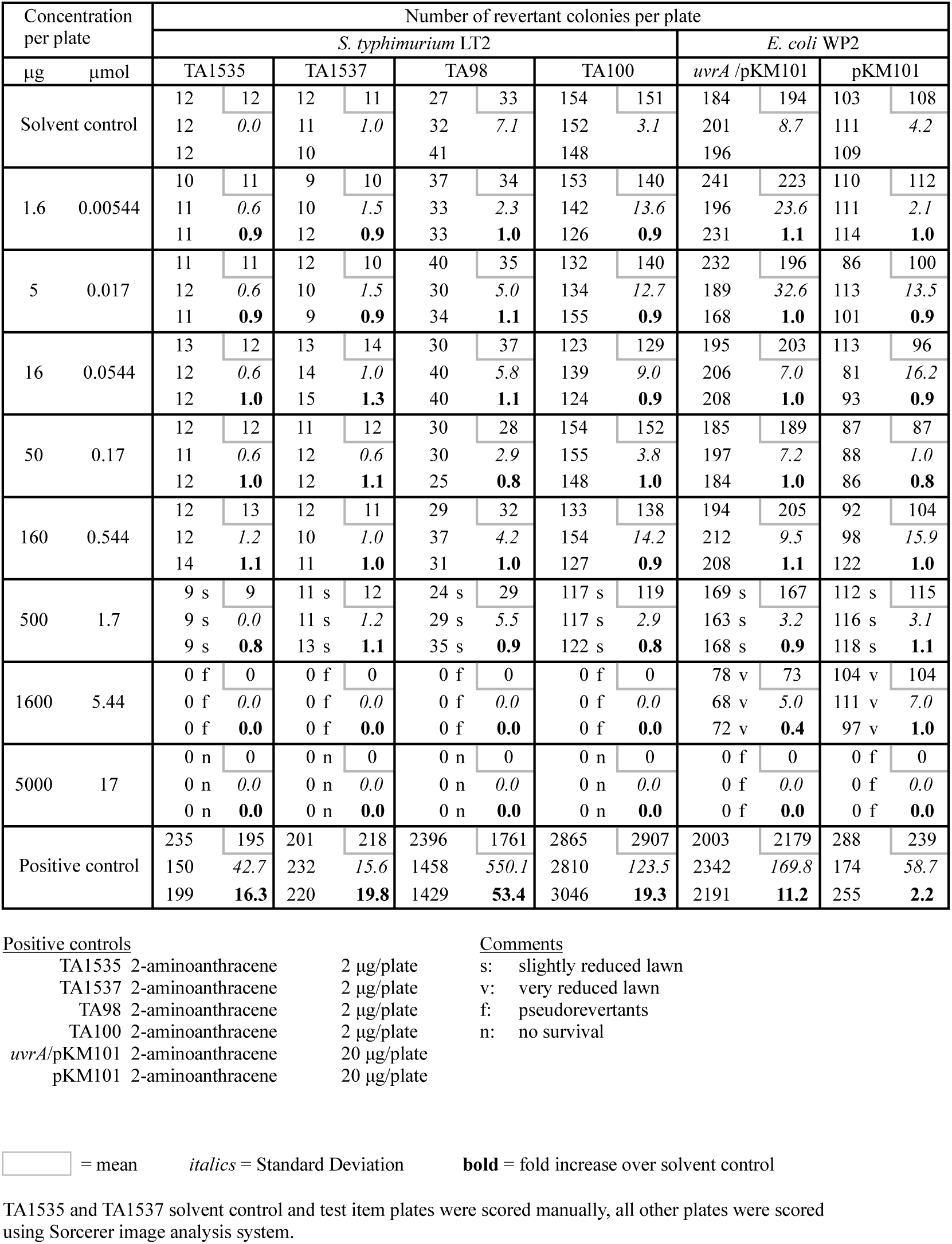
Plate Incorporation Experiment Results with SYN518580 in the Presence of Metabolic Activation.

**Supplementary Table 7:** Plate Incorporation Experiment with SYN518580 in the Absence of Metabolic Activation.

| Concentration<br>per plate | Number of revertant colonies per plate |  |  |  |  |  |  |  |  |  |  |  |
| --- | --- | --- | --- | --- | --- | --- | --- | --- | --- | --- | --- | --- |
|  | *S. typhimurium* LT2 |  |  |  |  |  |  |  | *E. coli* WP2 |  |  |  |
| µg µmol | TA1535 |  | TA1537 |  | TA98 |  | TA100 |  | *uvrA* /pKM101 |  | pKM101 |  |
| Solvent control | 12 | 12 | 12 | 12 | 29 | 29 | 133 | 132 | 163 | 159 | 72 | 74 |
|  | 12 | *0.6* | 13 | *0.6* | 29 | *0.6* | 132 | *1.5* | 159 | *4.5* | 81 | *5.9* |
|  | 13 |  | 12 |  | 28 |  | 130 |  | 154 |  | 70 |  |
| 1.6 0.00544 | 14 | 12 | 11 | 11 | 30 | 32 | 140 | 117 | 165 | 150 | 67 | 67 |
|  | 11 | *1.5* | 10 | *0.6* | 31 | *2.1* | 103 | *20.1* | 132 | *16.7* | 62 | *5.0* |
|  | 12 | **1.0** | 11 | **0.9** | 34 | **1.1** | 108 | **0.9** | 153 | **0.9** | 72 | **0.9** |
| 5 0.017 | 12 | 12 | 10 | 11 | 25 | 28 | 102 | 108 | 131 | 139 | 75 | 72 |
|  | 12 | *0.6* | 11 | *1.0* | 31 | *3.1* | 106 | *7.8* | 155 | *13.6* | 72 | *3.5* |
|  | 13 | **1.0** | 12 | **0.9** | 27 | **1.0** | 117 | **0.8** | 132 | **0.9** | 68 | **1.0** |
| 16 0.0544 | 12 | 13 | 14 | 13 | 31 | 30 | 112 | 111 | 150 | 155 | 71 | 73 |
|  | 13 | *0.6* | 14 | *1.2* | 29 | *1.2* | 108 | *2.3* | 162 | *6.4* | 73 | *1.5* |
|  | 13 | **1.1** | 12 | **1.1** | 29 | **1.0** | 112 | **0.8** | 152 | **1.0** | 74 | **1.0** |
| 50 0.17 | 12 | 12 | 10 | 10 | 31 | 30 | 128 | 122 | 130 | 133 | 60 | 67 |
|  | 13 | *0.6* | 9 | *0.6* | 27 | *2.3* | 126 | *8.1* | 139 | *5.5* | 67 | *6.5* |
|  | 12 | **1.0** | 10 | **0.8** | 31 | **1.0** | 113 | **0.9** | 129 | **0.8** | 73 | **0.9** |
| 160 0.544 | 18 | 17 | 11 | 12 | 29 | 29 | 114 | 118 | 155 | 154 | 75 | 74 |
|  | 18 | *1.7* | 12 | *1.0* | 29 | *0.6* | 114 | *6.9* | 140 | *13.5* | 77 | *3.6* |
|  | 15 | **1.4** | 13 | **1.0** | 30 | **1.0** | 126 | **0.9** | 167 | **1.0** | 70 | **1.0** |
| 500 1.7 | 26 r | 25 | 12 r | 12 | 34 r | 31 | 138 r | 128 | 76 r | 83 | 75 r | 83 |
|  | 27 r | *2.1* | 12 r | *0.6* | 27 r | *3.6* | 111 r | *14.8* | 97 r | *12.4* | 82 r | *8.5* |
|  | 23 r | **2.1** | 11 r | **1.0** | 32 r | **1.1** | 135 r | **1.0** | 75 r | **0.5** | 92 r | **1.1** |
| 1600 5.44 | 0 f | 0 | 0 f | 0 | 0 f | 0 | 0 f | 0 | 0 f | 0 | 0 f | 0 |
|  | 0 f | *0.0* | 0 f | *0.0* | 0 f | *0.0* | 0 f | *0.0* | 0 f | *0.0* | 0 f | *0.0* |
|  | 0 f | **0.0** | 0 f | **0.0** | 0 f | **0.0** | 0 f | **0.0** | 0 f | **0.0** | 0 f | **0.0** |
| 5000 17 | 0 n | 0 | 0 n | 0 | 0 n | 0 | 0 n | 0 | 0 n | 0 | 0 n | 0 |
|  | 0 n | *0.0* | 0 n | *0.0* | 0 n | *0.0* | 0 n | *0.0* | 0 n | *0.0* | 0 n | *0.0* |
|  | 0 n | **0.0** | 0 n | **0.0** | 0 n | **0.0** | 0 n | **0.0** | 0 n | **0.0** | 0 n | **0.0** |
| Positive control | 440 | 437 | 134 | 191 | 200 | 193 | 543 | 546 | 1066 | 1041 | 820 | 765 |
|  | 420 | *16.2* | 295 | *90.5* | 193 | *7.0* | 552 | *5.5* | 1048 | *29.1* | 700 | *60.6* |
|  | 452 | **36.4** | 143 | **15.9** | 186 | **6.7** | 542 | **4.1** | 1009 | **6.5** | 775 | **10.3** |

Positive controls
TA1535 sodium azide 0.5 µg/plate TA1537 9-aminoacridine.HCl 50 µg/plate TA98 2-nitrofluorene 1 µg/plate TA100 sodium azide 0.5 µg/plate *uvrA*/pKM101 potassium dichromate 25 µg/plate pKM101 potassium dichromate 25 µg/plate
Comments
r: reduced lawn f: pseudorevertants n: no survival
= mean *italics* = Standard Deviation **bold** = fold increase over solvent control
TA1535 and TA1537 solvent control and test item plates were scored manually, all other plates were scored using Sorcerer image analysis system.

**Supplementary Table 8:**
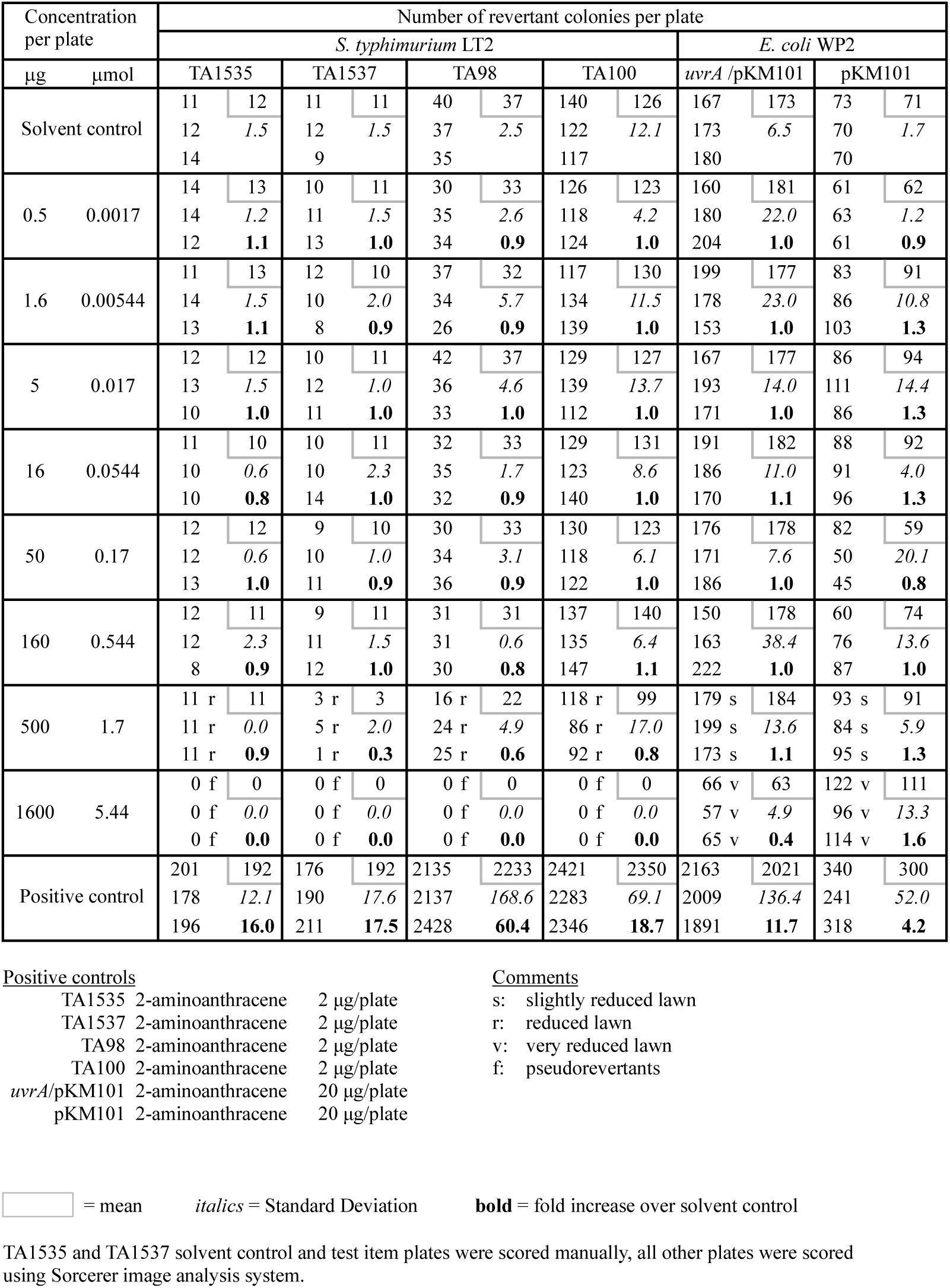
Liquid Pre-Incubation Experiment Results with SYN518580 in the Presence of Metabolic Activation.

**Supplementary Table 9:**
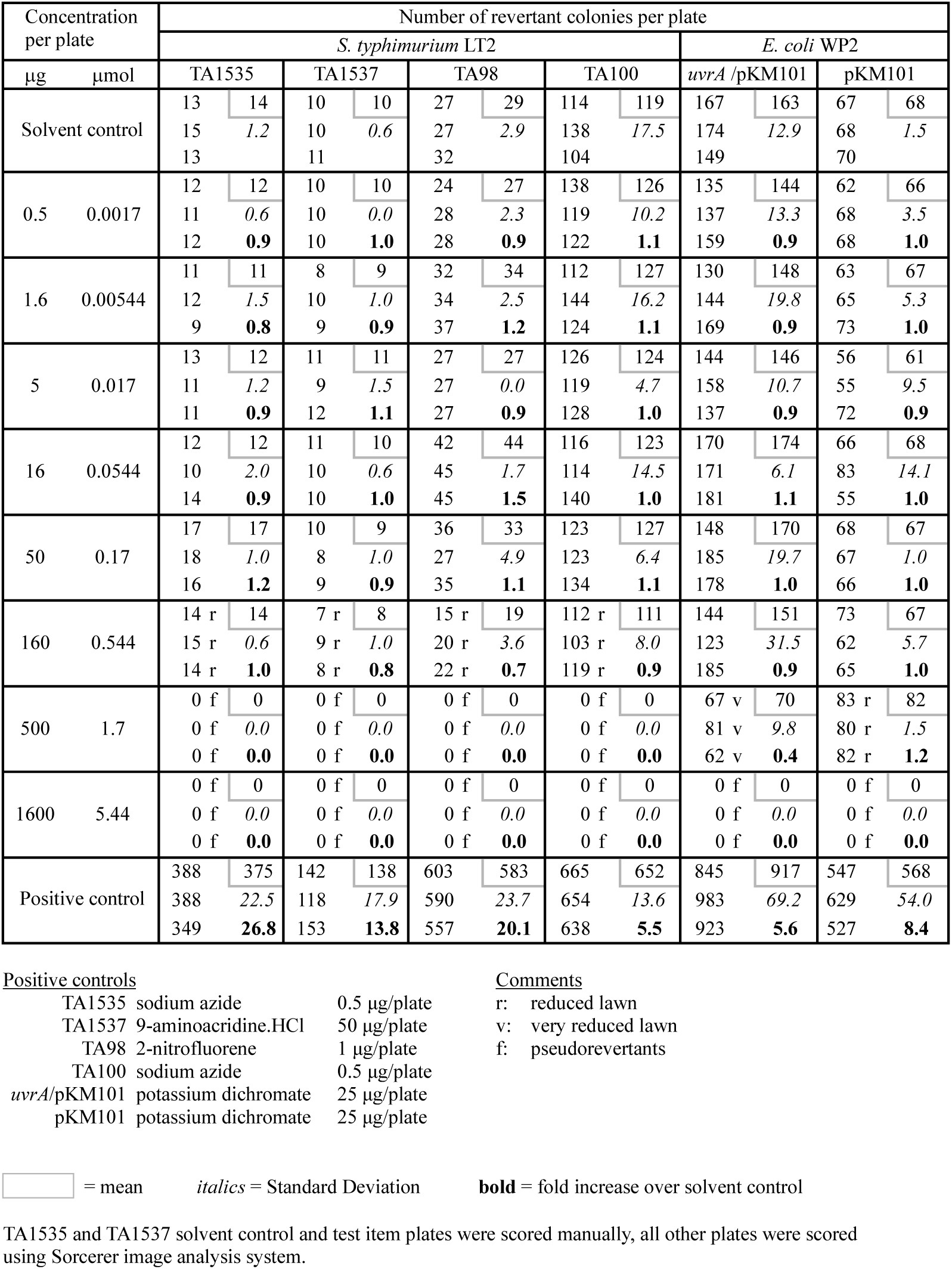
Liquid Pre-Incubation Experiment Results with SYN518580 in the Absence of Metabolic Activation.

**Supplementary Table 10:** Repeat Plate Incorporation Experiment Result (TA1535) with SYN518580 in the Absence of Metabolic Activation.

| Concentration per plate |  | Number of revertant colonies per plate |  |
| --- | --- | --- | --- |
|  |  | *S. typhimurium* LT2 |  |
| µg | µmol | TA1535 |  |
| Solvent control |  | 12 | 13 |
|  |  | 13 | *1.5* |
|  |  | 15 |  |
| 5 | 0.017 | 12 | 13 |
|  |  | 14 | *1.2* |
|  |  | 12 | **1.0** |
| 16 | 0.0544 | 11 | 11 |
|  |  | 12 | *0.6* |
|  |  | 11 | **0.8** |
| 50 | 0.17 | 12 | 12 |
|  |  | 11 | *0.6* |
|  |  | 12 | **0.9** |
| 160 | 0.544 | 12 | 13 |
|  |  | 13 | *1.5* |
|  |  | 15 | **1.0** |
| 250 | 0.85 | 11 s | 14 |
|  |  | 18 s | *3.6* |
|  |  | 13 s | **1.1** |
| 500 | 1.7 | 20 r | 20 |
|  |  | 20 r | *0.6* |
|  |  | 21 r | **1.5** |
| 1000 | 3.4 | 21 v | 19 |
|  |  | 15 v | *3.8* |
|  |  | 22 v | **1.5** |
| 1250 | 4.25 | 0 f | 0 |
|  |  | 0 f | *0.0* |
|  |  | 0 f | **0.0** |
| Positive control |  | 468 | 489 |
|  |  | 526 | *32.4* |
|  |  | 472 | **37.6** |

Positive controls
TA1535 sodium azide 0.5 µg/plate
Comments
s: slightly reduced lawn r: reduced lawn v: very reduced lawn f: pseudorevertants
= mean *italics* = Standard Deviation **bold** = fold increase over solvent control
TA1535 solvent control and test item plates were scored manually.

**Supplementary Table 11:**
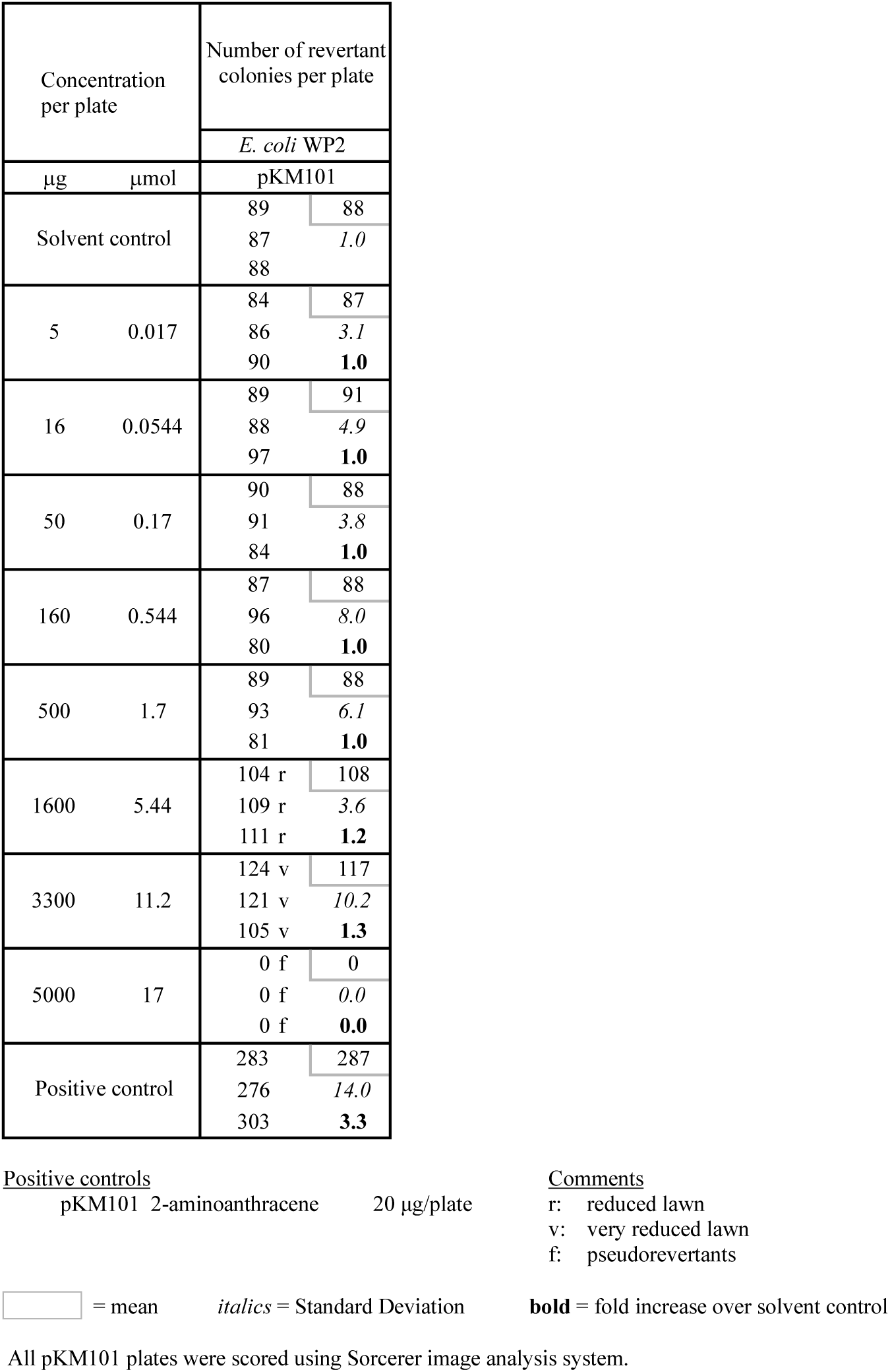
Repeat Liquid Pre-incubation Experiment Result (WP2 pKM101) with SYN518580 in the Presence of Metabolic Activation.

**Supplementary Table 12:**
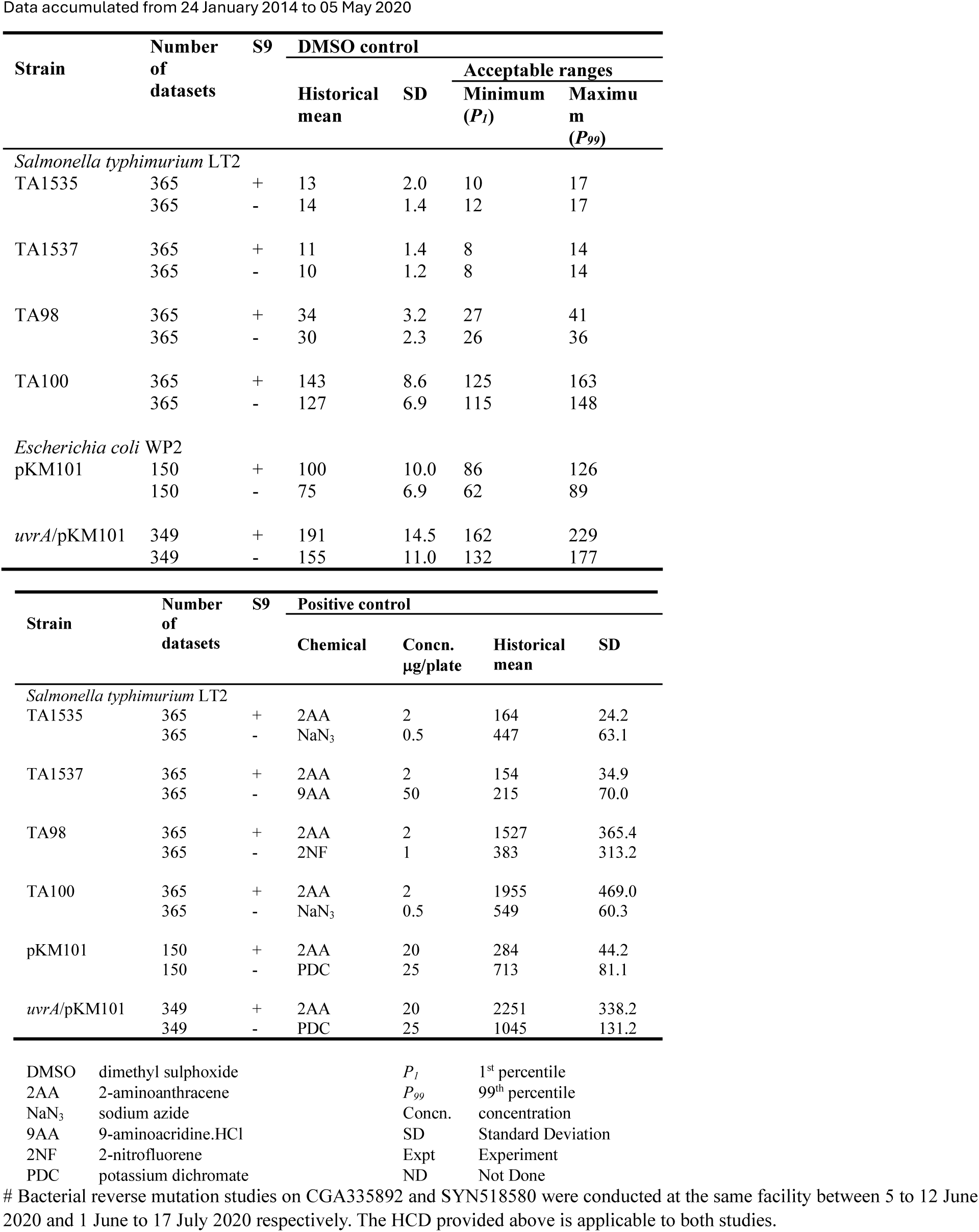
Historical Control Data for Bacterial Reverse Mutation Test on CGA335892 and SYN518580#.

Data accumulated from 24 January 2014 to 05 May 2020
| Strain | Number of datasets | S9 | DMSO control |  |  |  |
| --- | --- | --- | --- | --- | --- | --- |
|  |  |  | Historical mean | SD | Acceptable ranges |  |
| | | | | | Minimum ($P_1$) | Maximum ($P_{99}$) |
| *Salmonella typhimurium* LT2 |  |  |  |  |  |  |
| TA1535 | 365 | + | 13 | 2.0 | 10 | 17 |
|  | 365 | - | 14 | 1.4 | 12 | 17 |
| TA1537 | 365 | + | 11 | 1.4 | 8 | 14 |
|  | 365 | - | 10 | 1.2 | 8 | 14 |
| TA98 | 365 | + | 34 | 3.2 | 27 | 41 |
|  | 365 | - | 30 | 2.3 | 26 | 36 |
| TA100 | 365 | + | 143 | 8.6 | 125 | 163 |
|  | 365 | - | 127 | 6.9 | 115 | 148 |
| *Escherichia coli* WP2 |  |  |  |  |  |  |
| pKM101 | 150 | + | 100 | 10.0 | 86 | 126 |
|  | 150 | - | 75 | 6.9 | 62 | 89 |
| *uvrA*/pKM101 | 349 | + | 191 | 14.5 | 162 | 229 |
|  | 349 | - | 155 | 11.0 | 132 | 177 |

| Strain | Number of datasets | S9 | Positive control |  |  |  |
| --- | --- | --- | --- | --- | --- | --- |
| | | | Chemical | Concn. μg/plate | Historical mean | SD |
| *Salmonella typhimurium* LT2 |  |  |  |  |  |  |
| TA1535 | 365 | + | 2AA | 2 | 164 | 24.2 |
|  | 365 | - | NaN<sub>3</sub> | 0.5 | 447 | 63.1 |
| TA1537 | 365 | + | 2AA | 2 | 154 | 34.9 |
|  | 365 | - | 9AA | 50 | 215 | 70.0 |
| TA98 | 365 | + | 2AA | 2 | 1527 | 365.4 |
|  | 365 | - | 2NF | 1 | 383 | 313.2 |
| TA100 | 365 | + | 2AA | 2 | 1955 | 469.0 |
|  | 365 | - | NaN<sub>3</sub> | 0.5 | 549 | 60.3 |
| pKM101 | 150 | + | 2AA | 20 | 284 | 44.2 |
|  | 150 | - | PDC | 25 | 713 | 81.1 |
| *uvrA*/pKM101 | 349 | + | 2AA | 20 | 2251 | 338.2 |
|  | 349 | - | PDC | 25 | 1045 | 131.2 |

|  |  |  |  |
| --- | --- | --- | --- |
| DMSO | dimethyl sulphoxide | $P_1$ | 1<sup>st</sup> percentile |
| 2AA | 2-aminoanthracene | $P_{99}$ | 99<sup>th</sup> percentile |
| NaN<sub>3</sub> | sodium azide | Concn. | concentration |
| 9AA | 9-aminoacridine.HCl | SD | Standard Deviation |
| 2NF | 2-nitrofluorene | Expt | Experiment |
| PDC | potassium dichromate | ND | Not Done |

**Supplementary Table 13:** CGA335892 Plasma Concentration in the Mouse Following Two Doses by Oral Gavage.

| **Group** | **Animal** | **Timepoint (hours)** | **Dose level (mg/kg/day)** | **Result (ng/mL)** |
| --- | --- | --- | --- | --- |
| RF1M | 71 | 1 | 800 | 3470 |
| RF1M | 71 | 4 | 800 | 1860 |
| RF1M | 71 | 24 | 800 | 567 |
| RF1M | 72 | 1 | 800 | 2710 |
| RF1M | 72 | 4 | 800 | 1850 |
| RF1M | 72 | 24 | 800 | 722 |
| RF1M | 73 | 1 | 800 | 2660 |
| RF1M | 73 | 4 | 800 | 1610 |
| RF1M | 73 | 24 | 800 | 676 |
| RF1F | 74 | 1 | 800 | 1580 |
| RF1F | 74 | 4 | 800 | 882 |
| RF1F | 74 | 24 | 800 | 277 |
| RF1F | 75 | 1 | 800 | 4570 |
| RF1F | 75 | 4 | 800 | 3450 |
| RF1F | 75 | 24 | 800 | 495 |
| RF1F | 76 | 1 | 800 | 5030 |
| RF1F | 76 | 4 | 800 | 1500 |
| RF1F | 76 | 24 | 800 | 213 |

RF1 = Range-finder 1, M = male, F = female

**Supplementary Table 14:** Individual Animal Data from the in vivo Micronucleus Assay with CGA335892.

| Animal Number | Group | Dose CGA335892 (mg/kg/day) | RET | NCE | MN-RET | MN-NCE | % RET | MN-RET frequency |
| --- | --- | --- | --- | --- | --- | --- | --- | --- |
| 1 | 1 | 0 | 20014 | 1181513 | 35 | 1937 | 1.67 | 0.17 |
| 2 | 1 | 0 | 19867 | 1104311 | 45 | 1907 | 1.77 | 0.23 |
| 3 | 1 | 0 | 17096 | 1451784 | 30 | 2132 | 1.16 | 0.18 |
| 4 | 1 | 0 | 19591 | 1202012 | 57 | 2580 | 1.60 | 0.29 |
| 5 | 1 | 0 | 21204 | 2214549 | 36 | 3033 | 0.95 | 0.17 |
| 6 | 1 | 0 | 19738 | 1639215 | 45 | 3501 | 1.19 | 0.23 |
| 7 | 2 | 200 | 19879 | 1194431 | 53 | 2602 | 1.64 | 0.27 |
| 8 | 2 | 200 | 19856 | 1043737 | 36 | 1391 | 1.87 | 0.18 |
| 9 | 2 | 200 | 18343 | 1258451 | 30 | 1494 | 1.44 | 0.16 |
| 10 | 2 | 200 | 18525 | 1552948 | 37 | 2526 | 1.18 | 0.20 |
| 11 | 2 | 200 | 20359 | 1080161 | 51 | 1968 | 1.85 | 0.25 |
| 12 | 2 | 200 | 19850 | 964368 | 30 | 1283 | 2.02 | 0.15 |
| 13 | 3 | 400 | 20172 | 1295283 | 40 | 2237 | 1.53 | 0.20 |
| 14 | 3 | 400 | 19694 | 1349595 | 37 | 2039 | 1.44 | 0.19 |
| 15 | 3 | 400 | 20192 | 1045834 | 55 | 1845 | 1.90 | 0.27 |
| 16 | 3 | 400 | 19853 | 1041344 | 73 | 2830 | 1.87 | 0.37 |
| 17 | 3 | 400 | 18397 | 704196 | 75 | 1787 | 2.55 | 0.41 |
| 18 | 3 | 400 | 19244 | 1108298 | 46 | 2392 | 1.71 | 0.24 |
| 19 | 4 | 800 | 20629 | 1104741 | 39 | 1850 | 1.83 | 0.19 |
| 20 | 4 | 800 | 20118 | 1133159 | 52 | 1837 | 1.75 | 0.26 |
| 21 | 4 | 800 | 19908 | 953875 | 57 | 2077 | 2.05 | 0.29 |
| 22 | 4 | 800 | 20401 | 1341585 | 43 | 2583 | 1.50 | 0.21 |
| 23 | 4 | 800 | 20193 | 1345226 | 54 | 3357 | 1.48 | 0.27 |
| 24 | 4 | 800 | 20026 | 1383611 | 50 | 2700 | 1.43 | 0.25 |
| 101:4B | PC | CPA 30 mg/kg | 20564 | 9116174 | 177 | 12492 | 0.23 | 0.85 |
| 103:4B | PC | CPA 30 mg/kg | 20387 | 5477810 | 176 | 9011 | 0.37 | 0.86 |
| 105:4B | PC | CPA 30 mg/kg | 19579 | 4407322 | 143 | 8230 | 0.44 | 0.73 |
| 106:4B | PC | CPA 30 mg/kg | 19787 | 3908414 | 117 | 6017 | 0.51 | 0.59 |
| 107:4B | PC | CPA 30 mg/kg | 15979 | 3874115 | 98 | 7118 | 0.41 | 0.61 |
| 110:4A | PC | CPA 30 mg/kg | 15143 | 2785214 | 101 | 4395 | 0.54 | 0.66 |

CPA: cyclophosphamide, PC: positive Control bank

**Supplementary Table 15:**
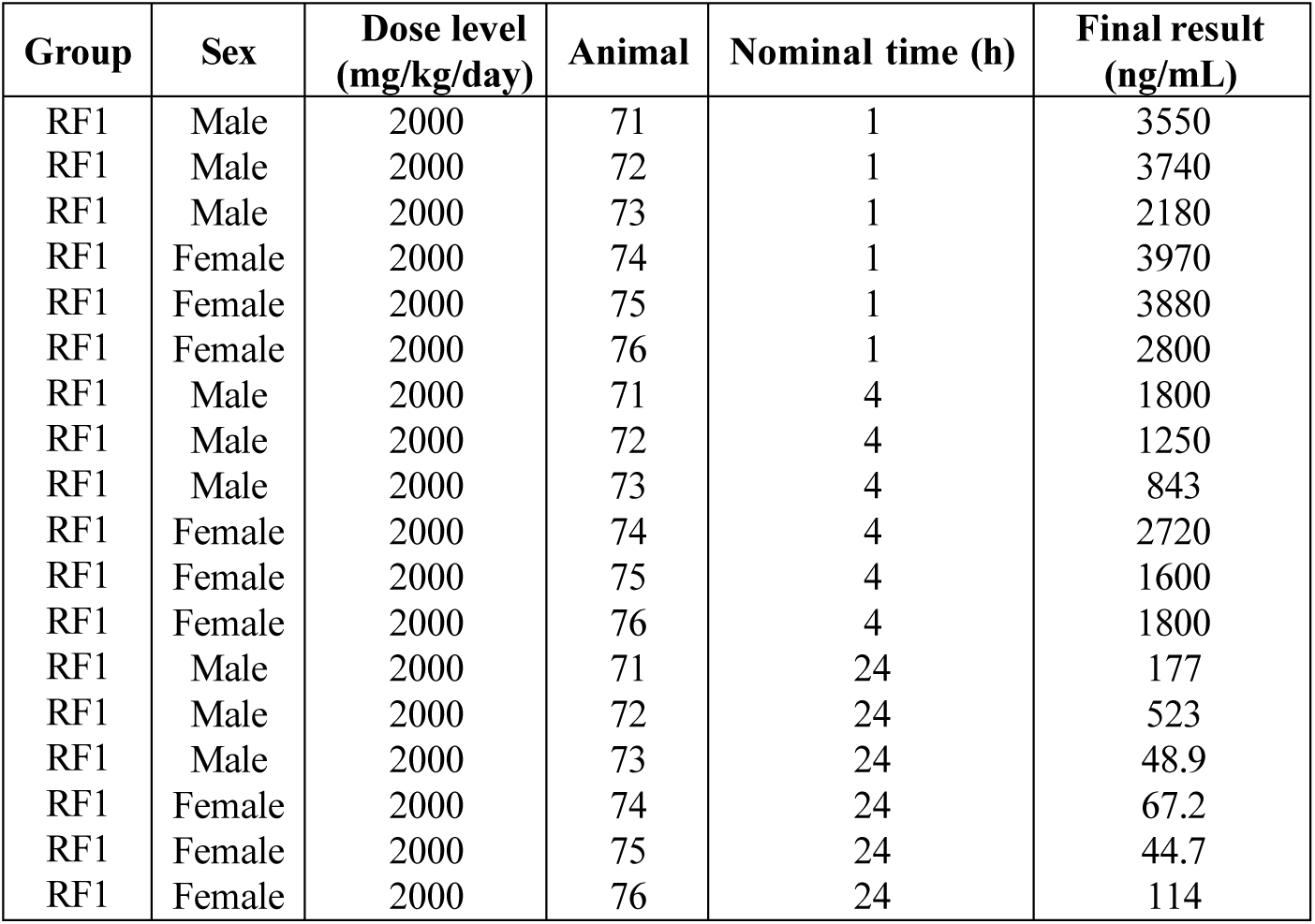
SYN518580 Plasma Concentration in the Mouse Following Two Doses by Oral Gavage.

**Supplementary Table 16:**
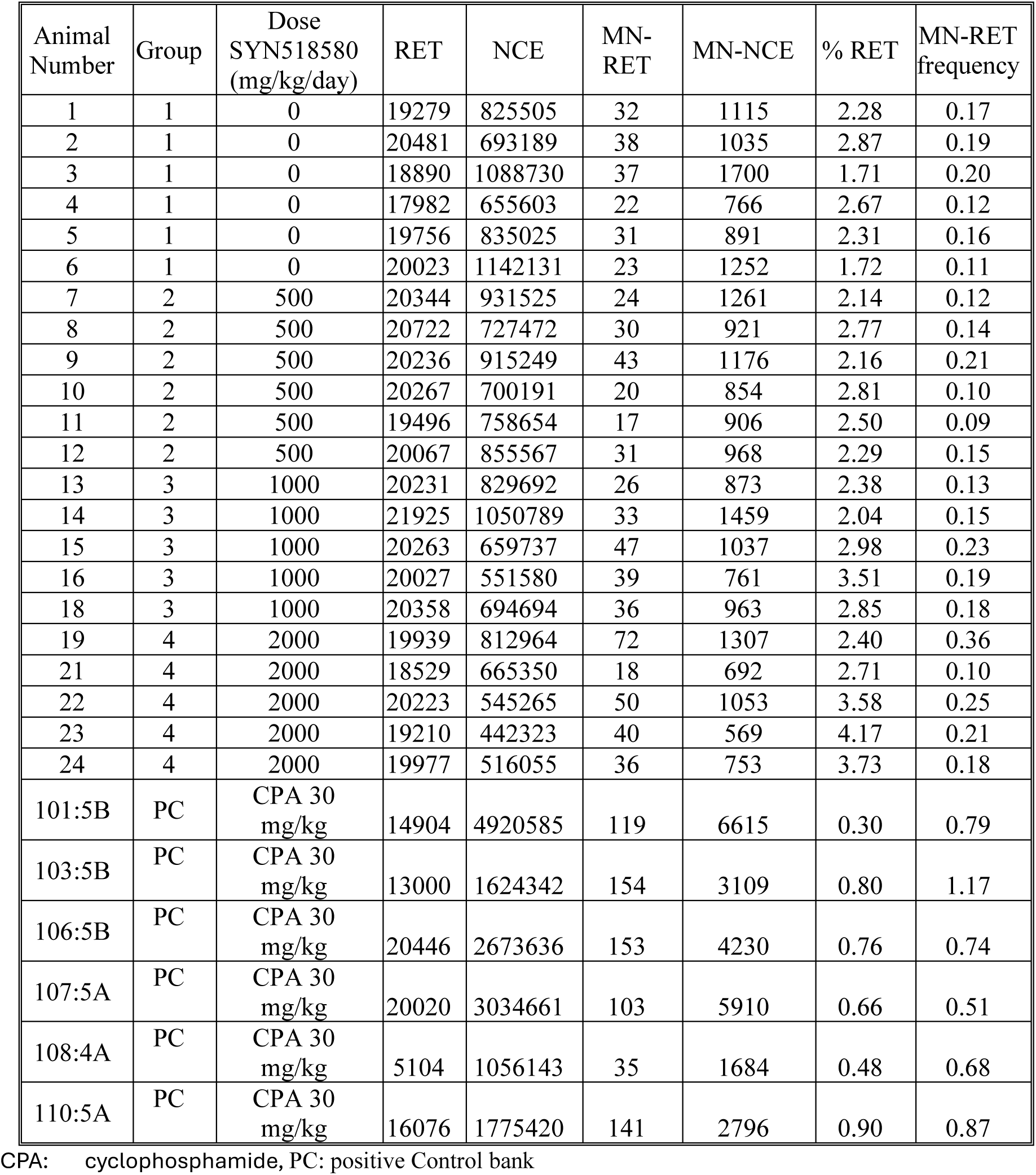
Individual Animal Data from the In Vivo Micronucleus Assay with SYN518580.

| Animal Number | Group | Dose SYN518580 (mg/kg/day) | RET | NCE | MN-RET | MN-NCE | % RET | MN-RET frequency |
| --- | --- | --- | --- | --- | --- | --- | --- | --- |
| 1 | 1 | 0 | 19279 | 825505 | 32 | 1115 | 2.28 | 0.17 |
| 2 | 1 | 0 | 20481 | 693189 | 38 | 1035 | 2.87 | 0.19 |
| 3 | 1 | 0 | 18890 | 1088730 | 37 | 1700 | 1.71 | 0.20 |
| 4 | 1 | 0 | 17982 | 655603 | 22 | 766 | 2.67 | 0.12 |
| 5 | 1 | 0 | 19756 | 835025 | 31 | 891 | 2.31 | 0.16 |
| 6 | 1 | 0 | 20023 | 1142131 | 23 | 1252 | 1.72 | 0.11 |
| 7 | 2 | 500 | 20344 | 931525 | 24 | 1261 | 2.14 | 0.12 |
| 8 | 2 | 500 | 20722 | 727472 | 30 | 921 | 2.77 | 0.14 |
| 9 | 2 | 500 | 20236 | 915249 | 43 | 1176 | 2.16 | 0.21 |
| 10 | 2 | 500 | 20267 | 700191 | 20 | 854 | 2.81 | 0.10 |
| 11 | 2 | 500 | 19496 | 758654 | 17 | 906 | 2.50 | 0.09 |
| 12 | 2 | 500 | 20067 | 855567 | 31 | 968 | 2.29 | 0.15 |
| 13 | 3 | 1000 | 20231 | 829692 | 26 | 873 | 2.38 | 0.13 |
| 14 | 3 | 1000 | 21925 | 1050789 | 33 | 1459 | 2.04 | 0.15 |
| 15 | 3 | 1000 | 20263 | 659737 | 47 | 1037 | 2.98 | 0.23 |
| 16 | 3 | 1000 | 20027 | 551580 | 39 | 761 | 3.51 | 0.19 |
| 18 | 3 | 1000 | 20358 | 694694 | 36 | 963 | 2.85 | 0.18 |
| 19 | 4 | 2000 | 19939 | 812964 | 72 | 1307 | 2.40 | 0.36 |
| 21 | 4 | 2000 | 18529 | 665350 | 18 | 692 | 2.71 | 0.10 |
| 22 | 4 | 2000 | 20223 | 545265 | 50 | 1053 | 3.58 | 0.25 |
| 23 | 4 | 2000 | 19210 | 442323 | 40 | 569 | 4.17 | 0.21 |
| 24 | 4 | 2000 | 19977 | 516055 | 36 | 753 | 3.73 | 0.18 |
| 101:5B | PC | CPA 30 mg/kg | 14904 | 4920585 | 119 | 6615 | 0.30 | 0.79 |
| 103:5B | PC | CPA 30 mg/kg | 13000 | 1624342 | 154 | 3109 | 0.80 | 1.17 |
| 106:5B | PC | CPA 30 mg/kg | 20446 | 2673636 | 153 | 4230 | 0.76 | 0.74 |
| 107:5A | PC | CPA 30 mg/kg | 20020 | 3034661 | 103 | 5910 | 0.66 | 0.51 |
| 108:4A | PC | CPA 30 mg/kg | 5104 | 1056143 | 35 | 1684 | 0.48 | 0.68 |
| 110:5A | PC | CPA 30 mg/kg | 16076 | 1775420 | 141 | 2796 | 0.90 | 0.87 |

CPA: cyclophosphamide, PC: positive Control bank

**Supplementary Table 17:** Summary of Mouse Micronucleus Negative and Positive Historical Control Data 2017 – 2020#.

| **Males Negative Control** |  |  |  |  |  |  |  |
| --- | --- | --- | --- | --- | --- | --- | --- |
| | N | Mean | SD | 95 % Control limit<br>(mean ± 2SD) | | Range<br>(min / max) | |
| MN-RET Frequency (%)<br>(MN-RET/RET) | 132 | 0.21 | 0.06 | 0.09 | 0.34 | 0.09 | 0.39 |
| % RET | 132 | 2.15 | 2.03 | -1.91 | 6.21 | 0.69 | 18.26 |
| **Males Positive Control<sup>1</sup>** |  |  |  |  |  |  |  |
| | N | Mean | SD | 95 % Control limit<br>(mean ± 2SD) | | Range<br>(min / max) | |
| MN-RET Frequency (%)<br>(MN-RET/RET) | 31 | 0.70 | 0.15 | 0.40 | 1.01 | 0.44 | 0.98 |
| % RET | 31 | 0.45 | 0.15 | 0.16 | 0.75 | 0.21 | 0.73 |

N: number of animals
Note: any discrepancy in this table is due to rounding differences
1: positive Control used was CPA 30 mg/kg administered by oral gavage.
Negative Control data was generated from individual animals from 20 experiments.

**Supplementary Table 18:**
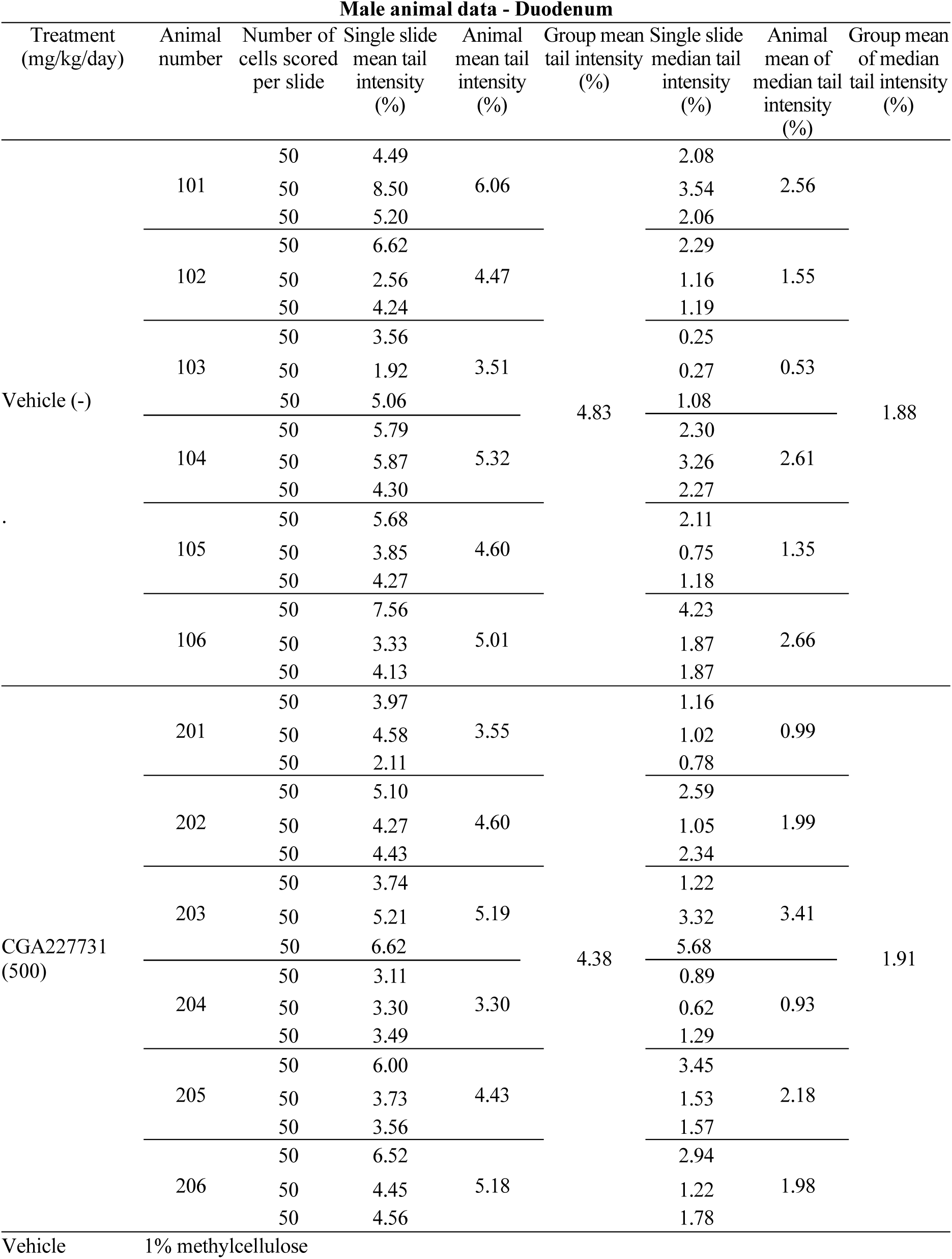
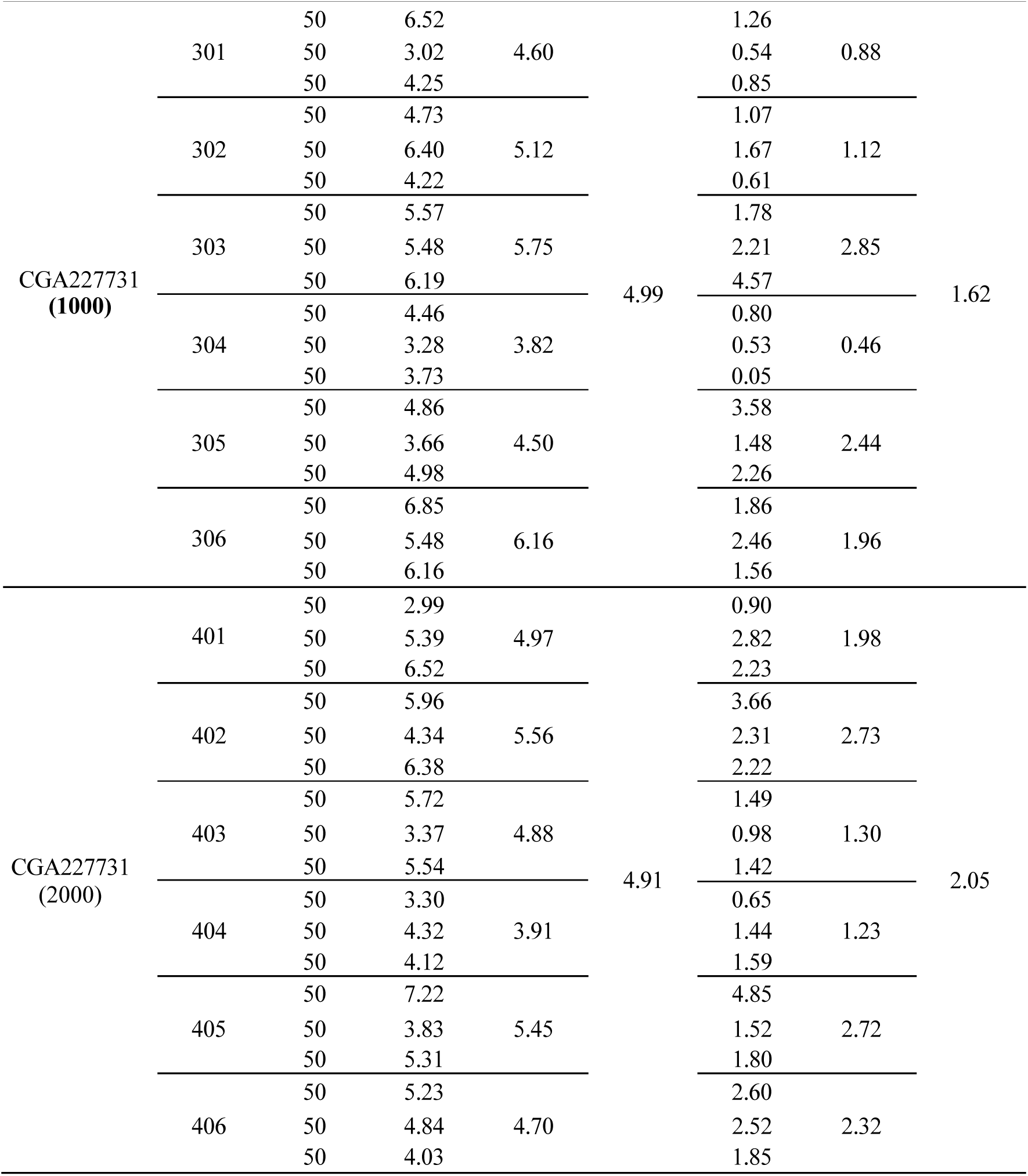
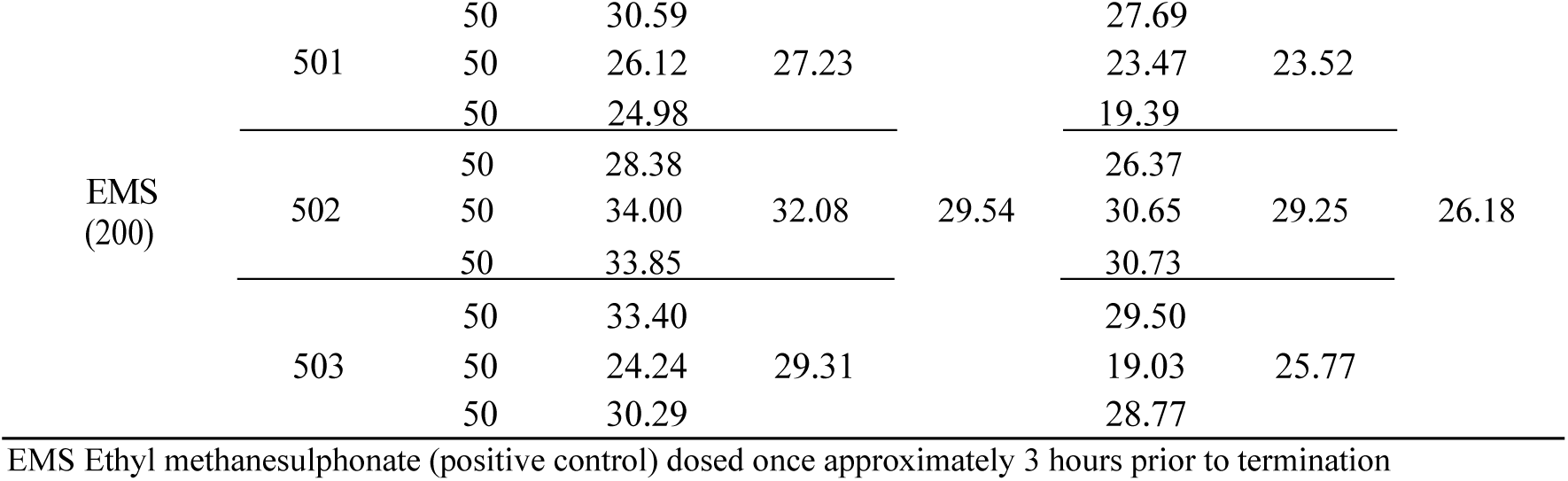
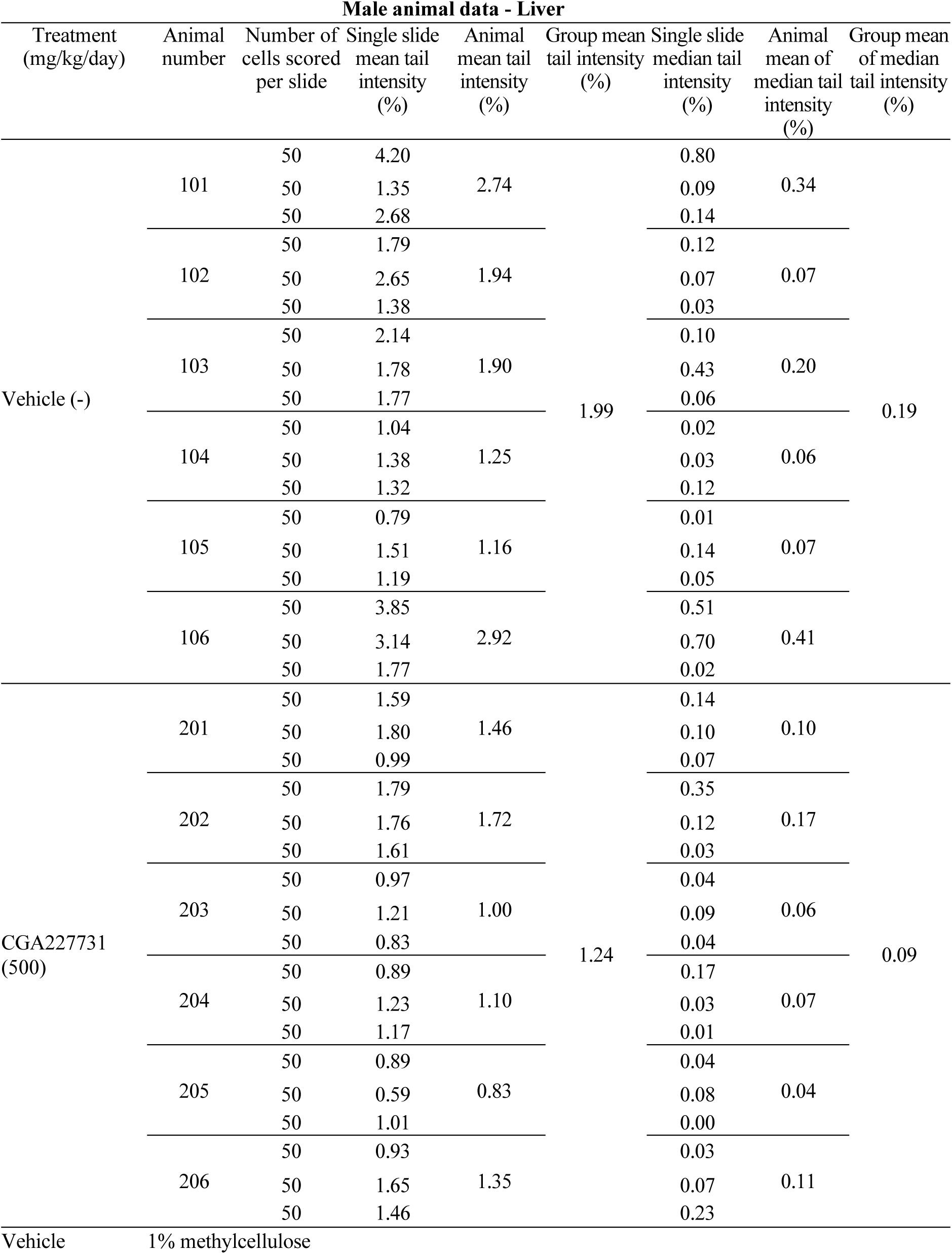
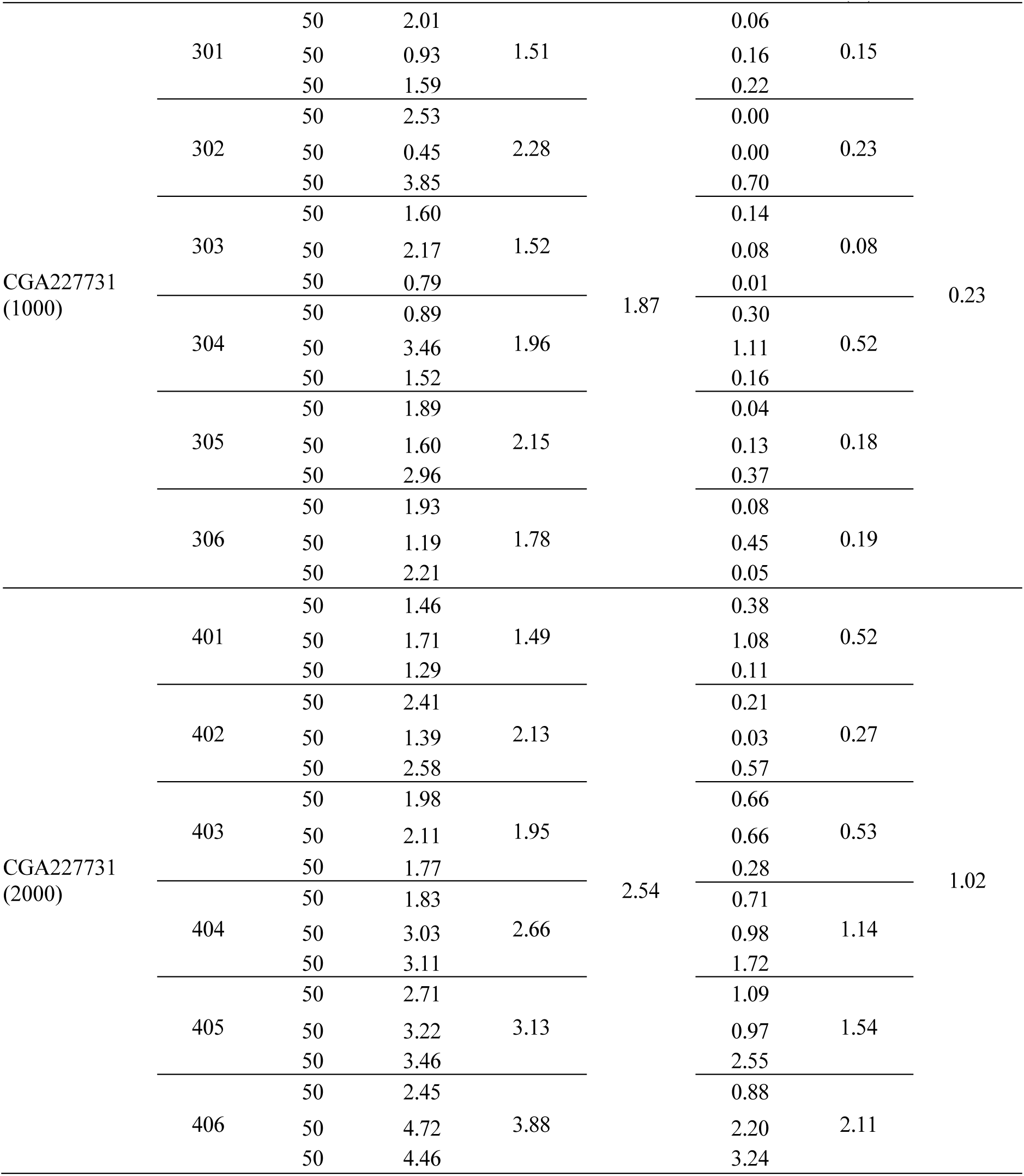
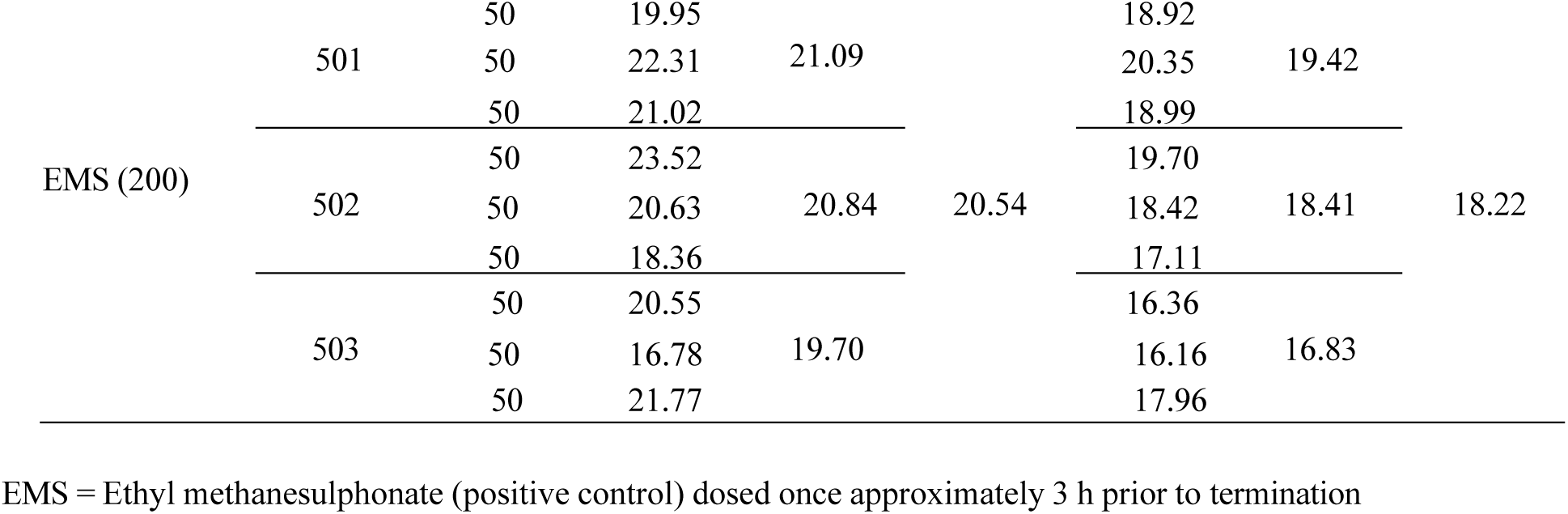
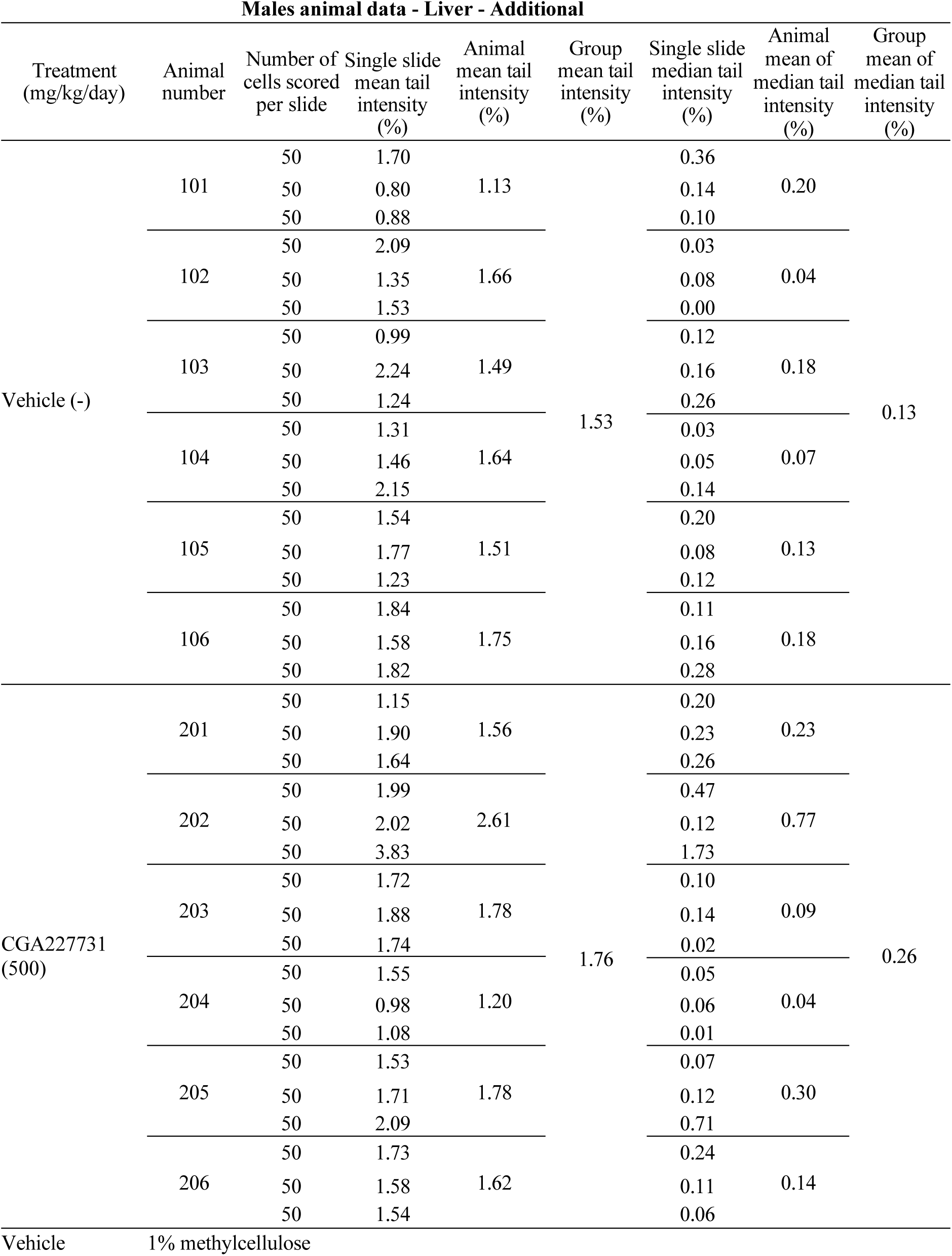
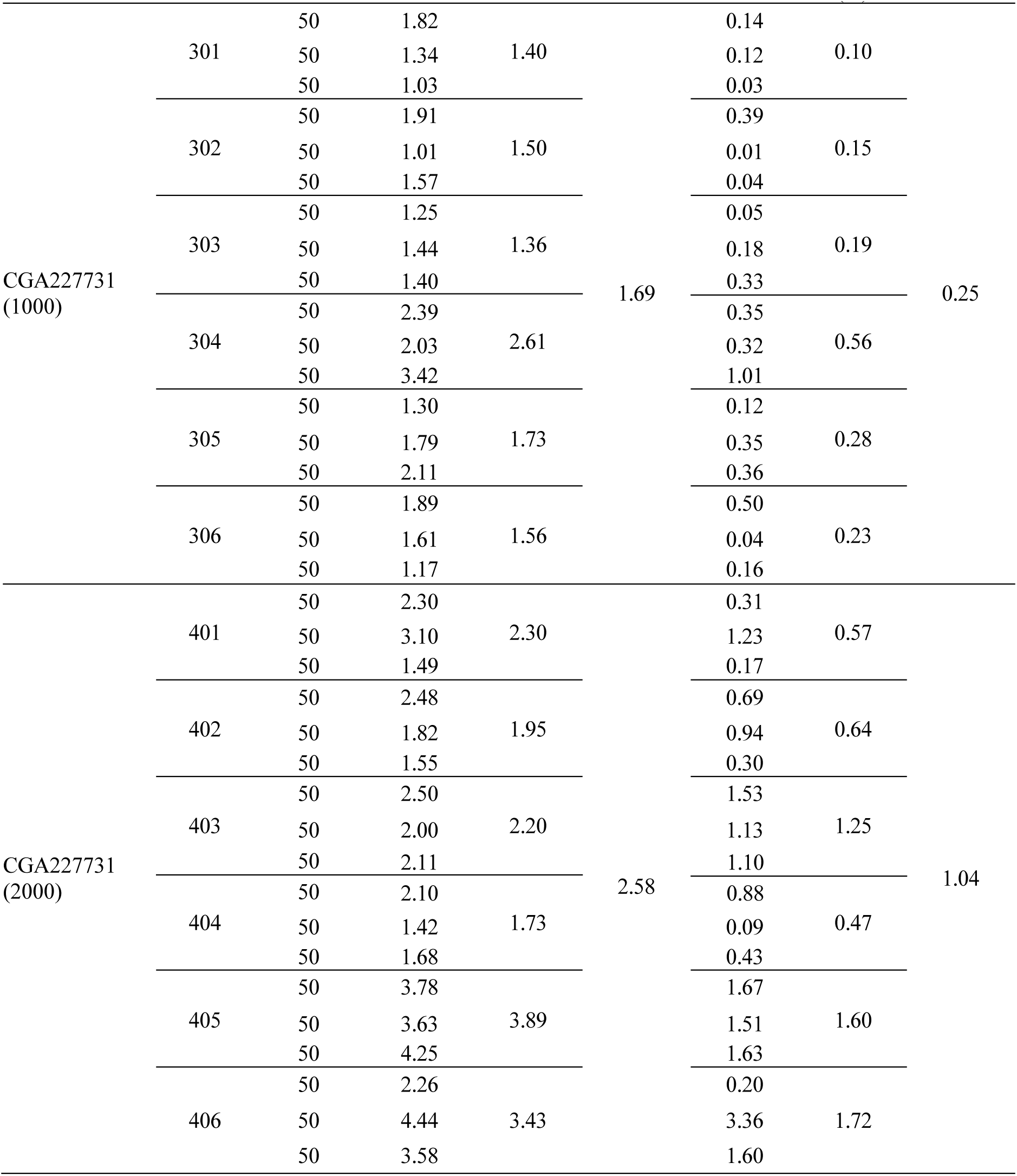
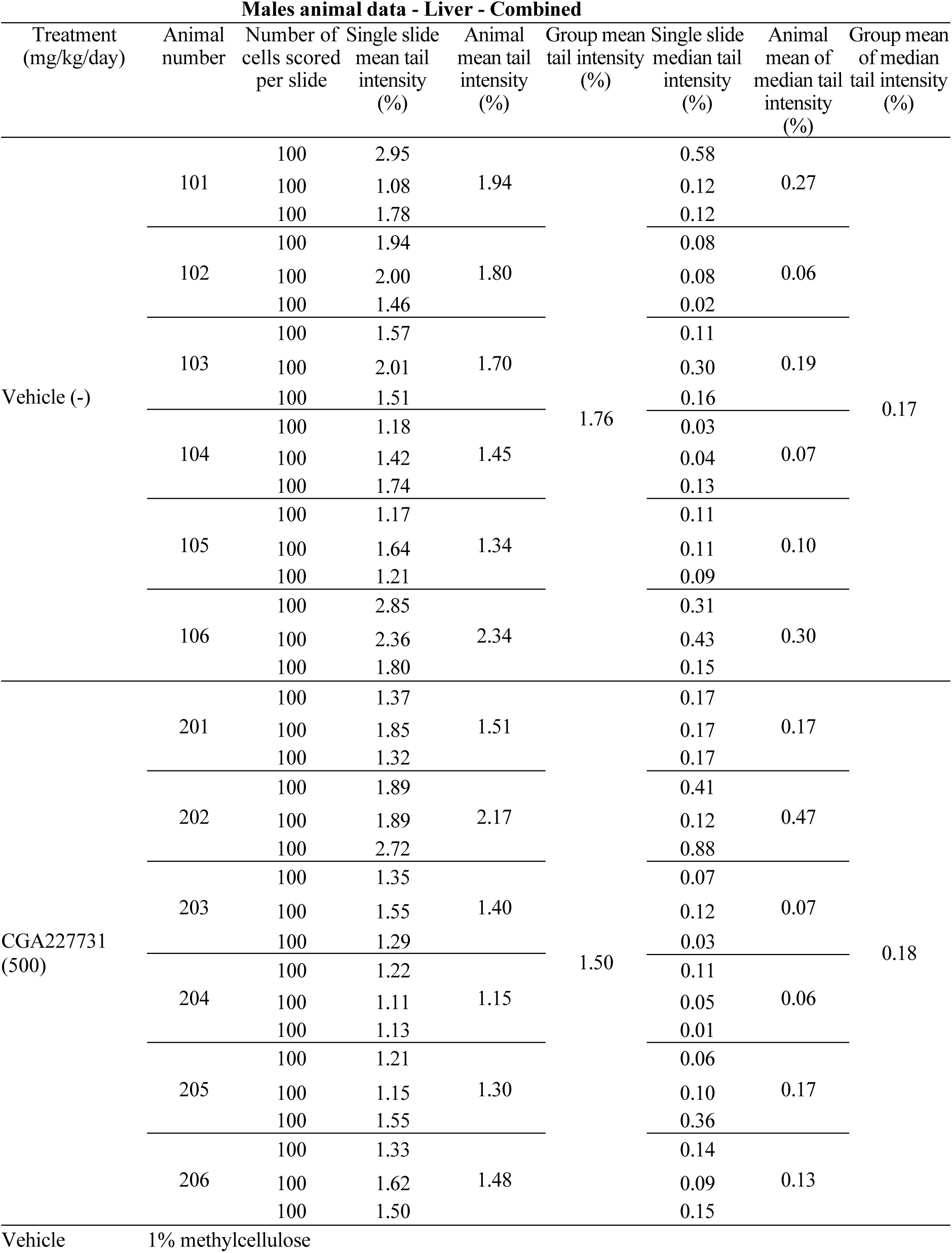
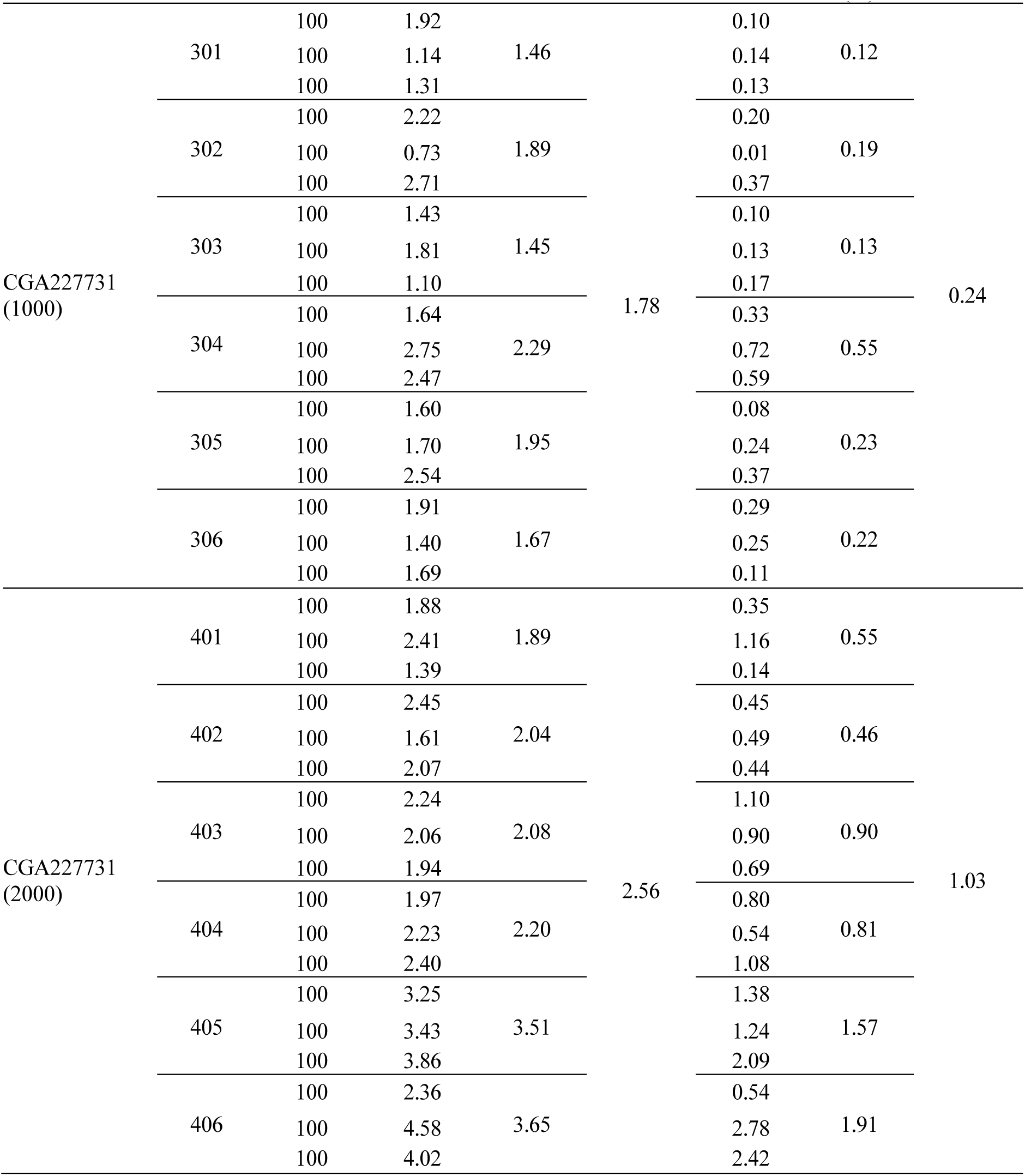
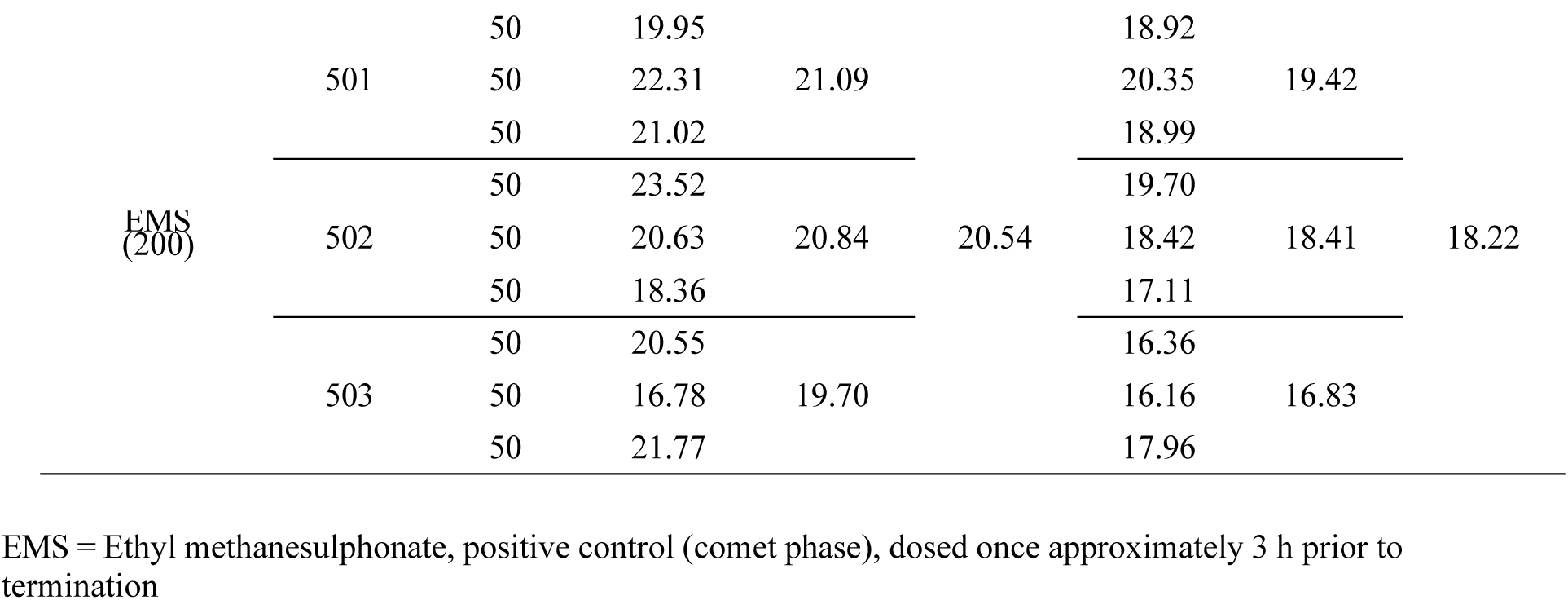
Individual Animal Data for the Rat Comet assay on CGA227731.

**Supplementary Table 19:**
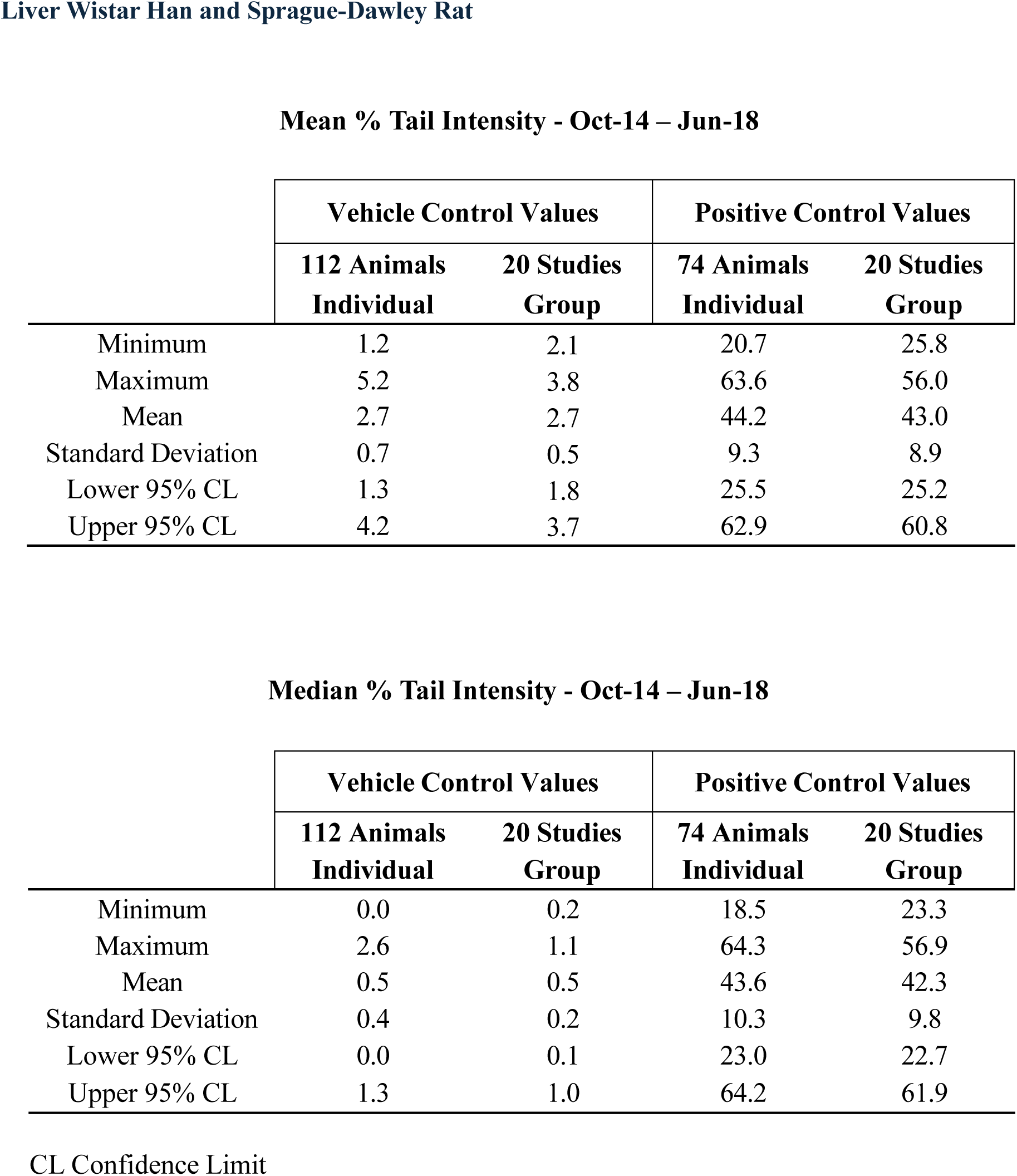
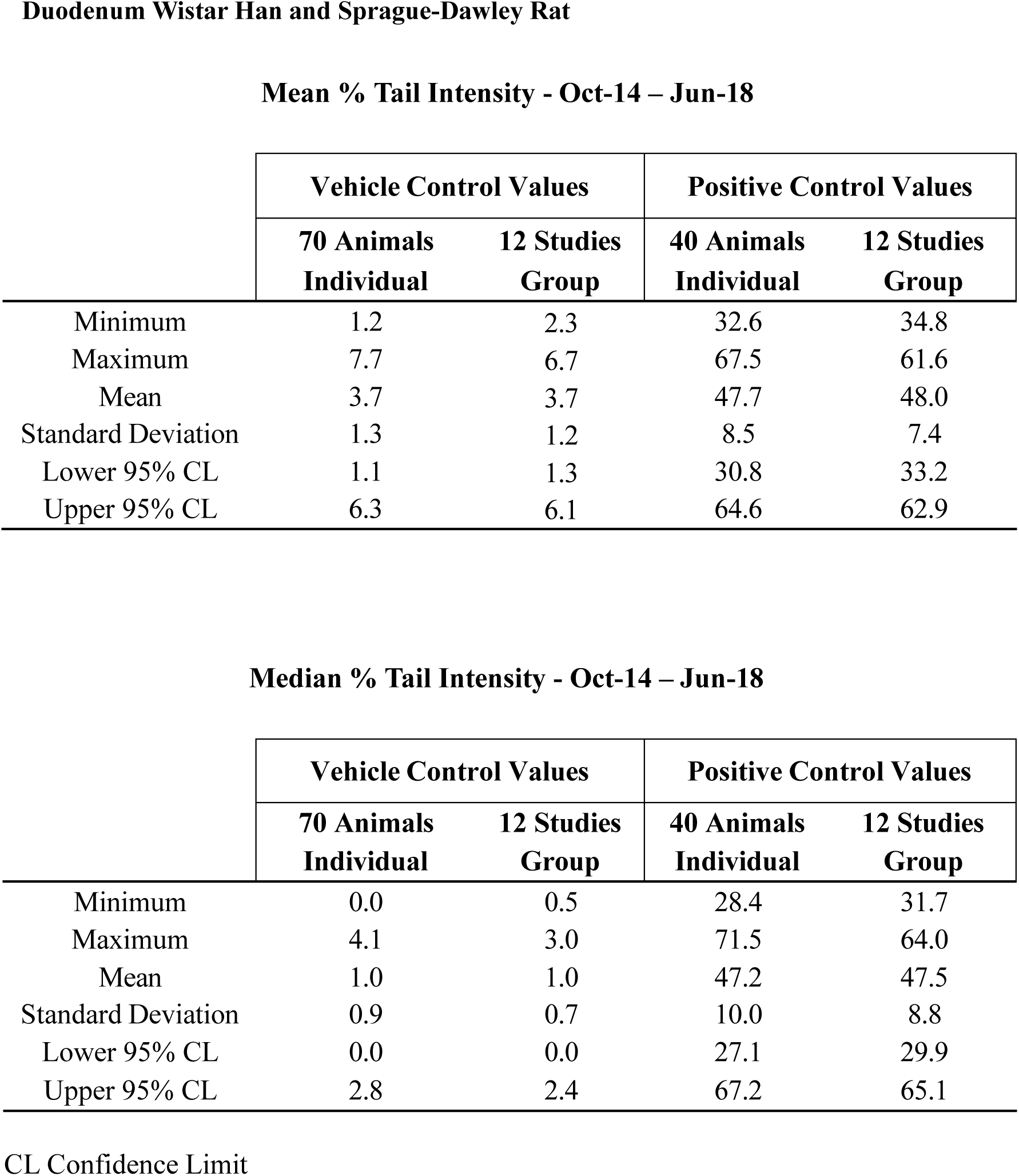
Comet Historical Control Data – Liver Wistar Han and Sprague-Dawley Rat.

**Mean % Tail Intensity - Oct-14 – Jun-18**
|  | Vehicle Control Values |  | Positive Control Values |  |
| --- | --- | --- | --- | --- |
|  | 112 Animals<br>Individual | 20 Studies<br>Group | 74 Animals<br>Individual | 20 Studies<br>Group |
| Minimum | 1.2 | 2.1 | 20.7 | 25.8 |
| Maximum | 5.2 | 3.8 | 63.6 | 56.0 |
| Mean | 2.7 | 2.7 | 44.2 | 43.0 |
| Standard Deviation | 0.7 | 0.5 | 9.3 | 8.9 |
| Lower 95% CL | 1.3 | 1.8 | 25.5 | 25.2 |
| Upper 95% CL | 4.2 | 3.7 | 62.9 | 60.8 |

**Median % Tail Intensity - Oct-14 – Jun-18**
|  | Vehicle Control Values |  | Positive Control Values |  |
| --- | --- | --- | --- | --- |
|  | 112 Animals<br>Individual | 20 Studies<br>Group | 74 Animals<br>Individual | 20 Studies<br>Group |
| Minimum | 0.0 | 0.2 | 18.5 | 23.3 |
| Maximum | 2.6 | 1.1 | 64.3 | 56.9 |
| Mean | 0.5 | 0.5 | 43.6 | 42.3 |
| Standard Deviation | 0.4 | 0.2 | 10.3 | 9.8 |
| Lower 95% CL | 0.0 | 0.1 | 23.0 | 22.7 |
| Upper 95% CL | 1.3 | 1.0 | 64.2 | 61.9 |

**Mean % Tail Intensity - Oct-14 – Jun-18**
|  | Vehicle Control Values |  | Positive Control Values |  |
| --- | --- | --- | --- | --- |
|  | 70 Animals<br>Individual | 12 Studies<br>Group | 40 Animals<br>Individual | 12 Studies<br>Group |
| Minimum | 1.2 | 2.3 | 32.6 | 34.8 |
| Maximum | 7.7 | 6.7 | 67.5 | 61.6 |
| Mean | 3.7 | 3.7 | 47.7 | 48.0 |
| Standard Deviation | 1.3 | 1.2 | 8.5 | 7.4 |
| Lower 95% CL | 1.1 | 1.3 | 30.8 | 33.2 |
| Upper 95% CL | 6.3 | 6.1 | 64.6 | 62.9 |

**Median % Tail Intensity - Oct-14 – Jun-18**
|  | Vehicle Control Values |  | Positive Control Values |  |
| --- | --- | --- | --- | --- |
|  | 70 Animals<br>Individual | 12 Studies<br>Group | 40 Animals<br>Individual | 12 Studies<br>Group |
| Minimum | 0.0 | 0.5 | 28.4 | 31.7 |
| Maximum | 4.1 | 3.0 | 71.5 | 64.0 |
| Mean | 1.0 | 1.0 | 47.2 | 47.5 |
| Standard Deviation | 0.9 | 0.7 | 10.0 | 8.8 |
| Lower 95% CL | 0.0 | 0.0 | 27.1 | 29.9 |
| Upper 95% CL | 2.8 | 2.4 | 67.2 | 65.1 |

CL Confidence Limit

**Supplementary Table 20:**
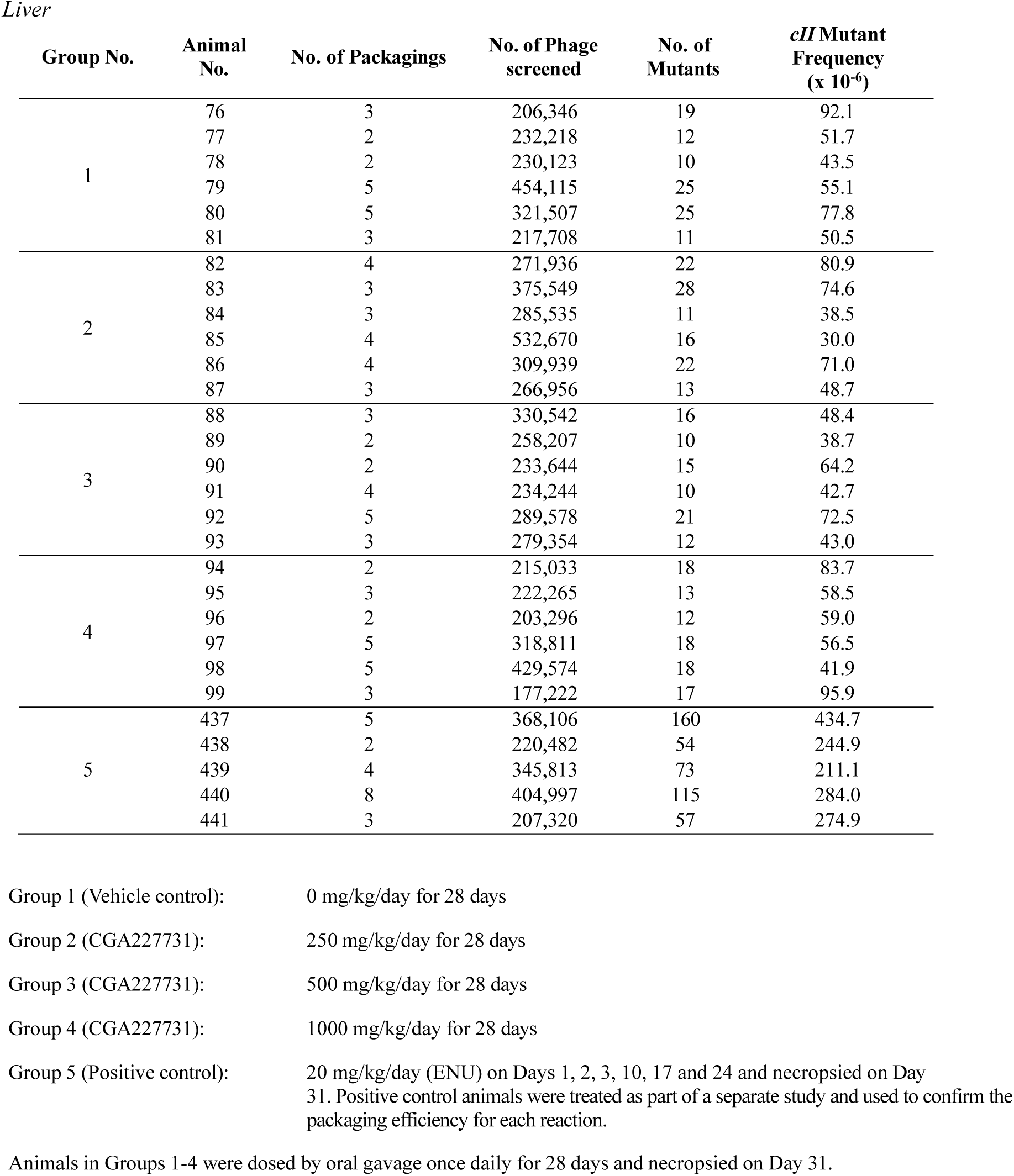
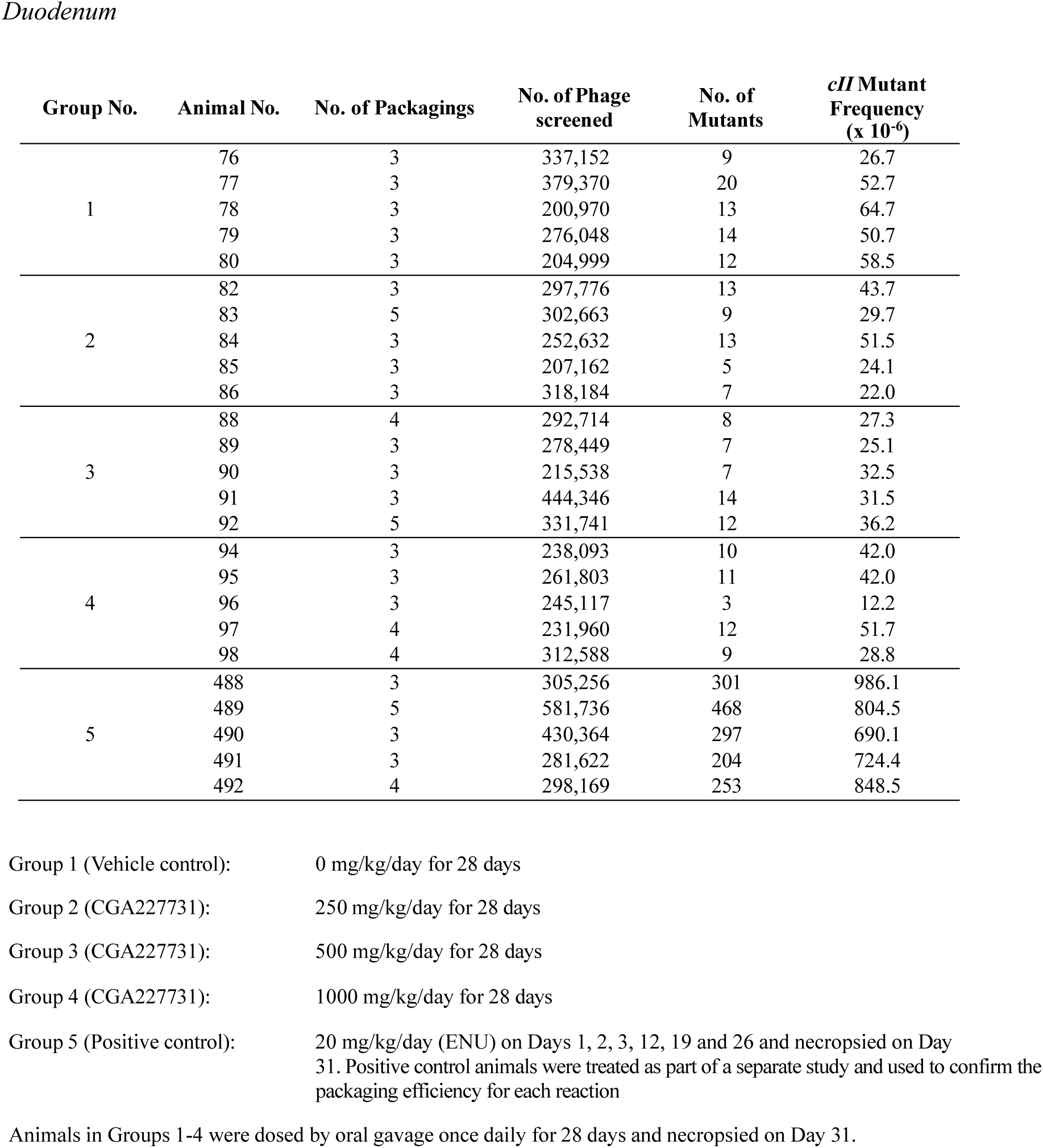
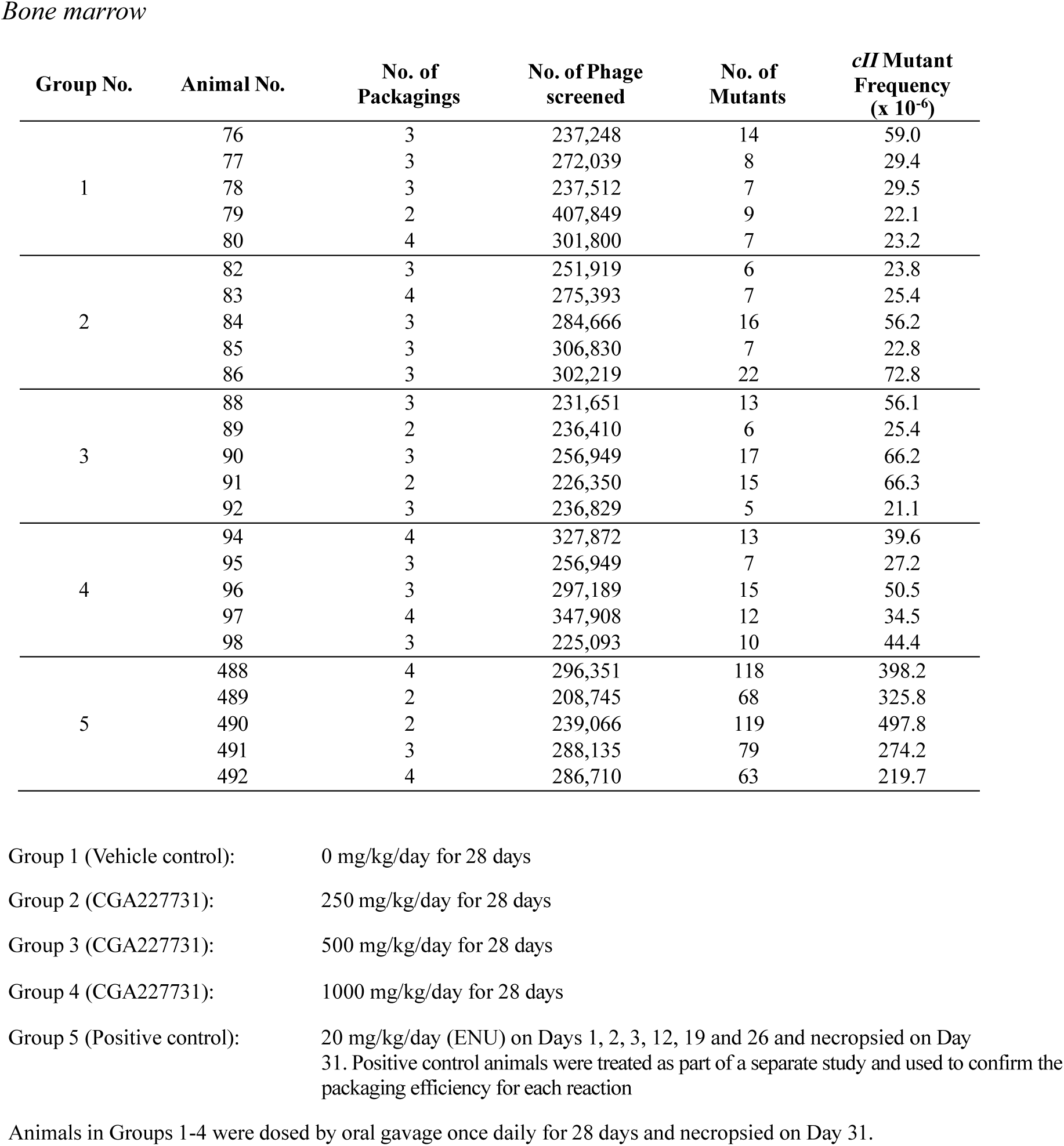
Individual animal data for cII mutations following treatment with CGA227731 in BigBlue® rats.

| Group No. | Animal No. | No. of Packagings | No. of Phage screened | No. of Mutants | *cII* Mutant Frequency (x 10<sup>-6</sup>) |
| --- | --- | --- | --- | --- | --- |
| 1 | 76 | 3 | 206,346 | 19 | 92.1 |
|  | 77 | 2 | 232,218 | 12 | 51.7 |
|  | 78 | 2 | 230,123 | 10 | 43.5 |
|  | 79 | 5 | 454,115 | 25 | 55.1 |
|  | 80 | 5 | 321,507 | 25 | 77.8 |
|  | 81 | 3 | 217,708 | 11 | 50.5 |
| 2 | 82 | 4 | 271,936 | 22 | 80.9 |
|  | 83 | 3 | 375,549 | 28 | 74.6 |
|  | 84 | 3 | 285,535 | 11 | 38.5 |
|  | 85 | 4 | 532,670 | 16 | 30.0 |
|  | 86 | 4 | 309,939 | 22 | 71.0 |
|  | 87 | 3 | 266,956 | 13 | 48.7 |
| 3 | 88 | 3 | 330,542 | 16 | 48.4 |
|  | 89 | 2 | 258,207 | 10 | 38.7 |
|  | 90 | 2 | 233,644 | 15 | 64.2 |
|  | 91 | 4 | 234,244 | 10 | 42.7 |
|  | 92 | 5 | 289,578 | 21 | 72.5 |
|  | 93 | 3 | 279,354 | 12 | 43.0 |
| 4 | 94 | 2 | 215,033 | 18 | 83.7 |
|  | 95 | 3 | 222,265 | 13 | 58.5 |
|  | 96 | 2 | 203,296 | 12 | 59.0 |
|  | 97 | 5 | 318,811 | 18 | 56.5 |
|  | 98 | 5 | 429,574 | 18 | 41.9 |
|  | 99 | 3 | 177,222 | 17 | 95.9 |
| 5 | 437 | 5 | 368,106 | 160 | 434.7 |
|  | 438 | 2 | 220,482 | 54 | 244.9 |
|  | 439 | 4 | 345,813 | 73 | 211.1 |
|  | 440 | 8 | 404,997 | 115 | 284.0 |
|  | 441 | 3 | 207,320 | 57 | 274.9 |
| 1 | 76 | 3 | 337,152 | 9 | 26.7 |
|  | 77 | 3 | 379,370 | 20 | 52.7 |
|  | 78 | 3 | 200,970 | 13 | 64.7 |
|  | 79 | 3 | 276,048 | 14 | 50.7 |
|  | 80 | 3 | 204,999 | 12 | 58.5 |
| 2 | 82 | 3 | 297,776 | 13 | 43.7 |
|  | 83 | 5 | 302,663 | 9 | 29.7 |
|  | 84 | 3 | 252,632 | 13 | 51.5 |
|  | 85 | 3 | 207,162 | 5 | 24.1 |
|  | 86 | 3 | 318,184 | 7 | 22.0 |
| 3 | 88 | 4 | 292,714 | 8 | 27.3 |
|  | 89 | 3 | 278,449 | 7 | 25.1 |
|  | 90 | 3 | 215,538 | 7 | 32.5 |
|  | 91 | 3 | 444,346 | 14 | 31.5 |
|  | 92 | 5 | 331,741 | 12 | 36.2 |
| 4 | 94 | 3 | 238,093 | 10 | 42.0 |
|  | 95 | 3 | 261,803 | 11 | 42.0 |
|  | 96 | 3 | 245,117 | 3 | 12.2 |
|  | 97 | 4 | 231,960 | 12 | 51.7 |
|  | 98 | 4 | 312,588 | 9 | 28.8 |
| 5 | 488 | 3 | 305,256 | 301 | 986.1 |
|  | 489 | 5 | 581,736 | 468 | 804.5 |
|  | 490 | 3 | 430,364 | 297 | 690.1 |
|  | 491 | 3 | 281,622 | 204 | 724.4 |
|  | 492 | 4 | 298,169 | 253 | 848.5 |
| 1 | 76 | 3 | 237,248 | 14 | 59.0 |
|  | 77 | 3 | 272,039 | 8 | 29.4 |
|  | 78 | 3 | 237,512 | 7 | 29.5 |
|  | 79 | 2 | 407,849 | 9 | 22.1 |
|  | 80 | 4 | 301,800 | 7 | 23.2 |
| 2 | 82 | 3 | 251,919 | 6 | 23.8 |
|  | 83 | 4 | 275,393 | 7 | 25.4 |
|  | 84 | 3 | 284,666 | 16 | 56.2 |
|  | 85 | 3 | 306,830 | 7 | 22.8 |
|  | 86 | 3 | 302,219 | 22 | 72.8 |
| 3 | 88 | 3 | 231,651 | 13 | 56.1 |
|  | 89 | 2 | 236,410 | 6 | 25.4 |
|  | 90 | 3 | 256,949 | 17 | 66.2 |
|  | 91 | 2 | 226,350 | 15 | 66.3 |
|  | 92 | 3 | 236,829 | 5 | 21.1 |
| 4 | 94 | 4 | 327,872 | 13 | 39.6 |
|  | 95 | 3 | 256,949 | 7 | 27.2 |
|  | 96 | 3 | 297,189 | 15 | 50.5 |
|  | 97 | 4 | 347,908 | 12 | 34.5 |
|  | 98 | 3 | 225,093 | 10 | 44.4 |
| 5 | 488 | 4 | 296,351 | 118 | 398.2 |
|  | 489 | 2 | 208,745 | 68 | 325.8 |
|  | 490 | 2 | 239,066 | 119 | 497.8 |
|  | 491 | 3 | 288,135 | 79 | 274.2 |
|  | 492 | 4 | 286,710 | 63 | 219.7 |

Group 1 (Vehicle control): 0 mg/kg/day for 28 days
Group 2 (CGA227731): 250 mg/kg/day for 28 days
Group 3 (CGA227731): 500 mg/kg/day for 28 days
Group 4 (CGA227731): 1000 mg/kg/day for 28 days
Group 5 (Positive control): 20 mg/kg/day (ENU) on Days 1, 2, 3, 12, 19 and 26 and necropsied on Day 31. Positive control animals were treated as part of a separate study and used to confirm the packaging efficiency for each reaction
Animals in Groups 1-4 were dosed by oral gavage once daily for 28 days and necropsied on Day 31.

**Supplementary Table 21:** Historical control data for cII mutations in BigBlue® Rat (2014-2021)

|  | Mutant frequency (x10<sup>-6</sup>) |  |  |  |  |  |
| --- | --- | --- | --- | --- | --- | --- |
|  | Liver |  | Duodenum |  | Bone Marrow |  |
|  | Individual | Studies | Individual | Studies | Individual | Studies |
| **Vehicle** |  |  |  |  |  |  |
| Mean | 41.9 | 41.5 | 40.3 | 41.0 | 29.4 | 29.4 |
| SD | 19.7 | 11.7 | 17.7 | 11.5 | 13.2 | 5.7 |
| 95% control limit | 2.5 - 81.3 | 18.1 - 64.9 | 4.9 - 75.7 | 18.0 - 64.0 | 3.0 - 55.8 | 18.0 - 40.8 |
| Range | 5.0 - 120.2 | 25.2 - 76.2 | 9.6 - 105.5 | 27.4 - 61.9 | 6.8 - 71.7 | 19.3 - 38.7 |
| N | 208 | 40 | 48 | 9 | 115 | 23 |
| **Positive control** |  |  |  |  |  |  |
| Mean | 210.6 | 209.4 | 747.8 | 747.6 | 388.2 | 391.9 |
| SD | 90.0 | 65.3 | 134.3 | 109.2 | 127.7 | 104.2 |
| 95% control limit | 30.6 - 390.6 | 78.8 - 340.0 | 479.2 – 1016.4 | 529.2 – 966.0 | 132.8 - 643.6 | 183.5 - 600.3 |
| Range | 60.8 - 614.1 | 116.4 - 389.9 | 439.3 – 967.6 | 593.7 – 840.9 | 143.3 - 1030.6 | 201.5 - 650.3 |
| N | 169 | 33 | 21 | 4 | 100 | 20 |

Vehicle control data pooled; dosed for 28 days and necropsied on Day 31. Duodenum HCD generated from vehicle controls dosed by oral gavage. For other tissues, other dosing routes are pooled. Bone marrow HCD also includes data from animals dosed by oral gavage for 28 days and necropsied on Day 50/56.
Positive control data; pooled positive controls; dosed with ENU; 20 mg/kg/day; Days 1, 2 and 3 and necropsied on Day 31; 20 mg/kg/day 1, 2, 3, 12, 19 and 26 and necropsied on Day 31; and 20 mg/kg/day 1, 2, 3, 10, 17 and 24 and necropsied on Day 31.

## Notes

### Competing Interest Statement

The authors are employees of Syngenta Ltd. The authors have no other competing interests.
The studies were conducted at 3rd party contract research organisations and funded by Syngenta Ltd.

## References

Agilent, 2018. RecoverEase™ DNA Isolation Kit, Instruction Manual. Agilent Document 720202-12, Revision D0. Santa Clara, CA. Available online at: www.agilent.com/cs/library/usermanuals/public/720202.pdf (last accessed 10 September 2025)

Agilent (2015a) Transpack Packaging Extract for Lambda Transgenic Shuttle Vector Recovery Instruction Manual. Agilent Document 200220-12. Revision B.0, La Jolla, CA. Available online at; www.agilent.com/cs/library/usermanuals/public/200220.pdf (last accessed 10 September 2025)

Agilent (2015b) λ Select-cII Mutation Detection System for Big Blue® Rodents”, 720120-12Revision B.0 La Jolla, CA. Available online at; www.agilent.com/cs/library/usermanuals/public/720120.pdf (last accessed 10 September 2025)

Ames B, McCann J and Yamasaki E (1975) Methods for Detecting Carcinogens and Mutagens with the Salmonella/Mammalian-Microsome Mutagenicity Test. Mutation Research/Environmental Mutagenesis and Related Subjects 31, 347–364

ANSES-French Agency for Food, Environmental and Occupational Health & Safety (2023) Fludioxonil Draft Renewal Assessment Report Volume 3 – B.6 (AS). Available online (Background documents) https://open.efsa.europa.eu/questions/EFSA-Q-2016-00481?search=fludioxonil; accessed 27 August 2025

Benigni R, Bossa C, Tcheremenskaia O and Worth A (2009) Development of Structural alerts for the in vivo micronucleus assay in rodents EUR 23844 EN – Joint Research Centre – Institute for Health and Consumer Protection EUR – Scientific and Technical Research series – ISSN 1018-5593

Benigni R, Bossa C and Worth A (2010) Structural analysis and predictive value of the rodent in vivo micronucleus assay results. Mutagenesis 25, 335–341

Guerard M, Marchand C and Plappert-Helbig U (2014) Influence of Experimental Conditions on Data Variability in the Liver Comet Assay. Environmental and Molecular Mutagenesis 55,114–121.

Guttenplan J, Spratt T, Khmelnitsky M, Kosinska W, Desai D, El-Bayoumy K (2004) Effects of 3H-1,2-dithiole-3-thione, 1,4-phenylenebis(methylene)selenocyanate, and selenium-enriched yeast individually and in combination on benzo[a]pyrene-induced mutagenesis in oral tissue and esophagus in lacZ mice. Mutation Research 559, 199–210.

Dertinger S, Li D, Beevers C, Douglas R, Heflich R H, Lovell DP, Roberts DJ,| Smith R, Uno Y, Williams A, Witt K L,| Zeller A, Zhou C (2023) Assessing the quality and making appropriate use of historical negative control data: A report of the International Workshop on Genotoxicity Testing (IWGT). Environmental and Molecular Mutagenesis 1–22

European Chemicals Agency (ECHA). (2017). Read-Across Assessment Framework (RAAF) Publications Office of the EU [DOI: 10.2823/794394].

European Food Safety Authority (EFSA) (2011) Scientific opinion on genotoxicity testing strategies applicable to food and feed safety assessment. EFSA Journal 9(9):2379

EFSA (2016). Technical report on the outcome of the pesticides peer review meeting on general recurring issues in mammalian toxicology. EFSA Journal 13(8): 1074

EFSA (2017) EFSA Scientific Committee, Hardy A, Benford D, Halldorsson T, Jeger M, Knutsen HK, More S, Naegeli H, Noteborn H, Ockleford C, Ricci A, Rychen G, Silano V, Solecki R, Turck D, Younes M, Aquilina G, Crebelli R, Gurtler R, Hirsch-Ernst KI, Mosesso P, Nielsen E, van Benthem J, Carf_ M, Georgiadis N, Maurici D, Parra Morte J and Schlatter J, 2017. Scientific Opinion on the clarification of some aspects related to genotoxicity assessment. EFSA Journal 15(12):5113

EFSA (2019) Pesticide Peer Review Meeting 01 – session 2 (01 – 05 April 2019) within document; Fludioxonil expert meeting reports August 2024_redacted_public. Available online (Background documents) https://open.efsa.europa.eu/questions/EFSA-Q-2016-00481?search=fludioxonil; accessed 27 August 2025

EFSA (2024) Peer review of the pesticide risk assessment of the active substance fludioxonil. EFSA Journal 22(11):9047

EFSA (2025). Guidance on the use of read-across for chemical safety assessment in food and feed. EFSA Journal, 23(7) 9586.

EU (2013) Commission Regulation (EU) No 283/2013 of 1 March 2013 setting out the data requirements for active substances, in accordance with Regulation (EC) No 1107/2009 of the European Parliament and of the Council concerning the placing of plant protection products on the market

Ferrari T and Gini G (2010) An open source model to predict mutagenicity from statistical analysis and relevant structural alerts. Chemistry Central Journal, 4(Suppl 1):S2

Henderson L, Wolfreys A, Fedyk J, Bourner C and Windebank S (1998) The ability of the Comet assay to discriminate between genotoxins and cytotoxins. Mutagenesis, 13, 89–94.

JMPR-Joint FAO/WHO Meeting on Pesticide Residues-Report 2022: pesticide residues in food. ISBN No.0-904147-72-X, 269–282

Judson P, Marchant CA, and Vessey J (2003) Using Argumentation for Absolute Reasoning about the Potential Toxicity of Chemicals. Journal of Chemical Information and Computer Sciences 43, 1364–1370

Judson P, Stalford SA and Vessey J (2013) Assessing confidence in predictions made by knowledge-based systems. Toxicology Research 2, 70–79.

Kirkland D, Levy D, LeBaron M, Aardema M, Beevers C, Bhalli J, Douglas G, Escobar P, Farabaugh C, Guerard M, Johnson G, Kulkarni R, Le Curieux F, Long A, Lott J, Lovell D, Luijten M, Marchetti F, Nicolette J, Pfuhler S, Roberts, Stankowski Jr. L, Thybaud V, Weiner S, Williams A, Witt K, Young R (2019) A comparison of transgenic rodent mutation and in vivo comet assay responses for 91 chemicals. Mutation research. Genetic toxicology and environmental mutagenesis 839, 21–35.

Klimisch H.-J., Andreae M, and Tillmann U (1997) A Systematic Approach for Evaluating the Quality of Experimental Toxicological and Ecotoxicological Data. Regulatory Toxicology and Pharmacology 25, 1–5

Lorenzo Y, Costa S, Collins A and Azqueta A (2013) The comet assay, DNA damage, DNA repair and cytotoxicity: hedgehogs are not always dead. Mutagenesis 28, 427–432

Mackay J and Elliot B (1992) Series: “Current Issues in Mutagenesis and Carcinogenesis” No. 29. Dose-ranging and dose setting for in vivo genetic toxicology studies. Mutation research 271, 97–99

Marchant CA, Briggs KA & Long A (2003). In silico tools for sharing data and knowledge on toxicity and metabolism: Derek for Windows, Meteor, and Vitic. Toxicology Mechanisms and Methods 18, 177–187.

Maron D and Ames B (1983). Revised methods for the Salmonella mutagenicity test. Mutation Research 113, 173–215

Mortelmans K, and Zeiger, E (2000) The Ames Salmonella/microsome mutagenicity assay. Mutation Research 455, 29–60

Mortelmans K. and Riccio E (2000) The bacterial tryptophan reverse mutation assay with *Escherichia coli* WP2. Mutation Research 455, 61–69.

OECD Guidelines for the Testing of Chemicals, Test Guideline No. 471. Bacterial Reverse Mutation Test 1997. Adopted: 21 July 1997

OECD Guideline for the testing of chemicals. Test Guideline No. 489 In Vivo Mammalian Alkaline Comet Assay. Adopted: 29 July 2016

OECD Guideline for the testing of chemicals. Test Guideline No. 488 Transgenic Rodent Somatic and Germ Cell Gene Mutation Assays Adopted: 26 June 2020

OECD Guideline for the testing of chemicals. Test Guideline No. 487 In Vitro Mammalian Cell Micronucleus Test. Adopted: 29 July 2016

OECD Guideline for the testing of chemicals. Test Guideline No. 474 Mammalian Erythrocyte Micronucleus Test. Adopted: 29 July 2016

OECD (2017a) Overview of the set of OECD Genetic Toxicology Test Guidelines and updates performed in 2014-2015. Series on Testing & Assessment No. 238 - 2nd edition. ENV/JM/MONO(2016)33/REV1

OECD (2017b), Guidance on Grouping of Chemicals, Second Edition, OECD Series on Testing and Assessment, No. 194, OECD Publishing, Paris, 10.1787/9789264274679-en

OECD (2024), (Q)SAR Assessment Framework: Guidance for the regulatory assessment of (Quantitative) Structure Activity Relationship models and predictions - Second edition, OECD Series on Testing and Assessment No. 405, OECD Publishing, Paris.

Powley M (2015) (Q)SAR assessments of potentially mutagenic impurities: A regulatory perspective on the utility of expert knowledge and data submission. Regulatory Toxicology and Pharmacology 71, 295–300

Robison TW, Heflich RH, Manjanatha MG, Elespuru R, Atrakchi A, Mei N, Ding W. (2021) Appropriate in vivo follow-up assays to an in vitro bacterial reverse mutation (Ames) test positive investigational drug candidate (active pharmaceutical ingredient), drug-related metabolite, or drug-related impurity. Mutation Research - Genetic Toxicology and Environmental Mutagenesis, 868–869,.

Sanderson DM and Earnshaw CG. (1991). Computer prediction of possible toxic action from chemical structure; the DEREK system. Human & Experimental Toxicology 10, 261–273

SANTE (2020). Guidance Document on Pesticide Analytical Methods for Risk Assessment and Post-approval Control and Monitoring Purposes SANTE/2020/12830 rev 1, 24 Feb 2021

Smith, C., Adkins, D. J., Martin, E. A., O’Donovan, M. R. (2008), “Recommendations for design of the rat comet assay”, Mutagenesis, Vol. 23/3, pp. 233–40

Vasquez MZ, Dewhurst NE (2024) Reducing risk of false positives in the in vivo comet assay and improving result reliability. Mutation Research - Genetic Toxicology and Environmental Mutagenesis 895, 503–750.

Venitt S, Crofton-Sleigh C and Forster R (1984). Bacterial Mutation Assays Using Reverse Mutation. In: Venitt S and Parry JM, Eds. Mutagenicity Testing a Practical Approach, -IRL Press Oxford, England, ISBN No.0-904147-72-X, 45–98.

Zeller A, Pfuhler S, Albertini S, Bringezu F, Czich A, Dietz Y, Fautz R, Hewitt NJ, Kirst and Kasper P (2018) A critical appraisal of the sensitivity of in vivo genotoxicity assays in detecting human carcinogens. Mutagenesis, 33, 179–193.

Zeller A, Brigo A, Brink A, Guerard M, Lang D, Muster W, Runge F, Sutter A, Vock E, Wichard J, and Schadt S, (2020) Genotoxicity Assessment of Drug Metabolites in the Context of MIST and Beyond Chemical Research in Toxicology, 33, 10–19.

